# Peri-Locus Coeruleus Controls Endothelial Cell Heterogeneity in Enriched Environments via Region-Specific Norepinephrine Signaling

**DOI:** 10.64898/2026.06.23.733358

**Authors:** Yingzi Zhao, Botai Li, Furong Ju, Jiesi Feng, Ji-an Wei, Jie Shao, Weikang Sun, Shuai Chen, Yue Qiu, Dashuang Gao, Lu Zhang, Yuewen Chen, Yulong Li, Liping Wang, Jie Tu, Fan Yang

## Abstract

The brain acts as a central integrator that enables organisms to interpret environmental stimuli and coordinate adaptive responses across the body. While enriched environments are known to enhance neural plasticity and cognitive function, their impact on peripheral organ systems remains less understood. Here, we show that enriched environmental conditions activate a peri-locus coeruleus (peri-LC) neuronal ensemble that preserves CD31^hi^EMCN^hi^ type H endothelium, a specialized bone-associated endothelial subtype, against age-associated decline, thereby enhancing femoral bone density. Chemogenetic activation and ablation experiments further revealed that this peri-LC ensemble is functionally distinct from other sympathetic regulatory brain regions in its control of bone endothelial heterogeneity and skeletal mass. Three-dimensional bone clearing and spatial analyses showed that CD31^hi^EMCN^hi^ endothelium is closely associated with sympathetic nerve terminals. The peri-LC mediates this process through sympathetic signaling and β2-adrenergic receptors on bone vascular endothelial cells. *In vivo* imaging further demonstrated that norepinephrine release is spatially restricted, with sprouting endothelial cells near the growth plate exhibiting the highest sensitivity. Together, these findings reveal a refined brain–periphery regulatory logic in which distinct sympathetic brain circuits engage spatially restricted norepinephrine signaling to coordinate organ-specific vascular and skeletal adaptations.

## Introduction

The ability to perceive and adapt to environmental changes is essential for the survival of all organisms. As a dynamic and adaptive organ, the brain plays a pivotal role in sensing and processing external information, with psychological states guiding responses to environmental cues ^1^. Exposure to enriched environments (EE), characterized by increased sensory, cognitive, and social stimulation compared to standard conditions, has been shown to enhance neural plasticity, improve cognitive function, and support overall brain health ^2^. Beyond its role in sensory and cognitive functions, the brain has long been recognized as a master regulator of systemic physiological processes, including cardiovascular function, respiratory control, immune responses ^3–5^, nutrient preferences, and metabolic pathways ^6,7^. However, the mechanisms by which enriched environments influence the body through the brain remain underexplored. In the current study, we investigated the effects of enriched environments on brain activity and its potential to mediate adaptive changes in peripheral organs via neural-vascular interactions.

The central and peripheral nervous systems are closely integrated with the vascular system, as evidenced by the considerable spatial and morphological alignment between the peripheral nervous and vascular networks ^8^. Beyond structural parallels, these systems exhibit reciprocal influences in growth, development, and function ^8–10^. Neurotransmitters such as norepinephrine (NE), acetylcholine, and adenosine triphosphate (ATP), released by the peripheral nervous system, modulate vascular smooth muscle and endothelial cell activity, thereby regulating critical physiological parameters, including heart rate, blood pressure, and endothelial function ^11,12^. Dysregulation of these processes is closely associated with pathologies such as heart failure, hypertension, and atherosclerosis ^13–15^. Sympathetic nervous system activation has also been implicated in tumor progression by promoting angiogenesis, which increases cancer cell proliferation and tumor growth ^16^. These findings emphasize the critical role of the peripheral nervous system in regulating vascular functions. However, how the nervous system governs endothelial heterogeneity and organ specificity remains relatively unexplored.

Endothelial cells exhibit extensive heterogeneity and organ-specific adaptations ^10^. These cells secrete angiocrine factors that participate in diverse physiological processes, including neural development and regeneration, reproduction, bone marrow hematopoiesis, liver and lung regeneration, and bone and lipid metabolism ^17^. The skeletal system contains a specialized vascular endothelial subtype, characterized by high expression of platelet endothelial cell adhesion molecule (PECAM-1, also known as CD31) and endomucin (EMCN), termed CD31^hi^EMCN^hi^ or type H endothelium. With their unique morphological, molecular, and functional properties, these vessels localize to specific regions to facilitate bone vasculature growth, create distinct metabolic and molecular microenvironments, support perivascular osteoprogenitors, and link angiogenesis with osteogenesis ^18^. As aging progresses, however, the expression of CD31 and EMCN in type H endothelial cells decreases, leading to a transition to type L endothelium (CD31^lo^EMCN^lo^), which lacks active metabolic growth and osteogenesis properties ^18^. Given the critical role of CD31^hi^EMCN^hi^ endothelium in bone growth and development, it represents a promising therapeutic target for osteoporosis ^19,20^. Despite advances in understanding endothelial function, the mechanisms through which the central and peripheral nervous systems influence the phenotypic plasticity of CD31^hi^EMCN^hi^ and CD31^lo^EMCN^lo^ bone endothelial phenotypes remain largely unexplored. Investigating these mechanisms holds the potential to reveal novel insights into the neural regulation of vascular specialization and bone health.

This study elucidates a brain–body interaction through which enriched environments are translated into neural signals that shape peripheral endothelial phenotypes. We identify an EE-responsive peri-locus coeruleus (peri-LC) neuronal ensemble that regulates skeletal vascular endothelial heterogeneity through the peripheral sympathetic nervous system, thereby promoting bone formation. This peri-LC ensemble is functionally distinct from other sympathetic regulatory brain regions in its control of bone endothelium and skeletal mass, highlighting the regional specificity of central sympathetic circuits in peripheral organ regulation. Three-dimensional bone clearing and two-photon imaging further revealed close neurovascular interactions in bone, with peri-LC stimulation triggering spatially restricted norepinephrine release preferentially received by sprouting endothelial cells. Together, these findings suggest that the peri-LC acts beyond its classical role in arousal to orchestrate long-term peripheral vascular and skeletal adaptations, revealing a refined brain–periphery regulatory logic by which distinct brain regions control specific organ phenotypes through sympathetic signaling.

## Results

### Enriched environments enhance peri-LC activity

Six-week-old mice were housed in an enriched environment (actual design was showed in Figure S1A, B and methods) characterized by larger, more complex spaces, diverse objects, and higher cage population density compared to standard conditions ^2,21^. To identify brain regions responsive to enriched environments exposure, transgenic TRAP2 mice were used. These mice enable the labeling of neurons with increased immediate early gene *Fos* expression using tdTomato as a marker for neural activation, facilitating brain-wide screening to identify regions affected by environmental enrichment (Figure 1A). Brain-wide analysis was conducted using tissue clearing, alignment to a reference brain atlas, and automated cell counting. This comprehensive approach revealed a significant increase in neuronal activity across the brains of EE-housed mice compared to standard-housed controls, as indicated by heightened tdTomato expression (Figure S1C-G, Video 1, Figure 1B, and Table 1). Notable increases in neural activity were observed in multiple brain regions, including the basolateral amygdala (BLA), basomedial amygdala (BMA), ventral striatum (STRv), medial hypothalamic zone (MEZ), pons (P), medulla oblongata (MY), and hemispheric region (HEM) (Figure 1B, S1F and Table 1). These findings suggest that enriched environments robustly activate neural circuits across key areas responsible for information processing, emotional regulation, motor coordination, and homeostatic control.

**Figure 1:**
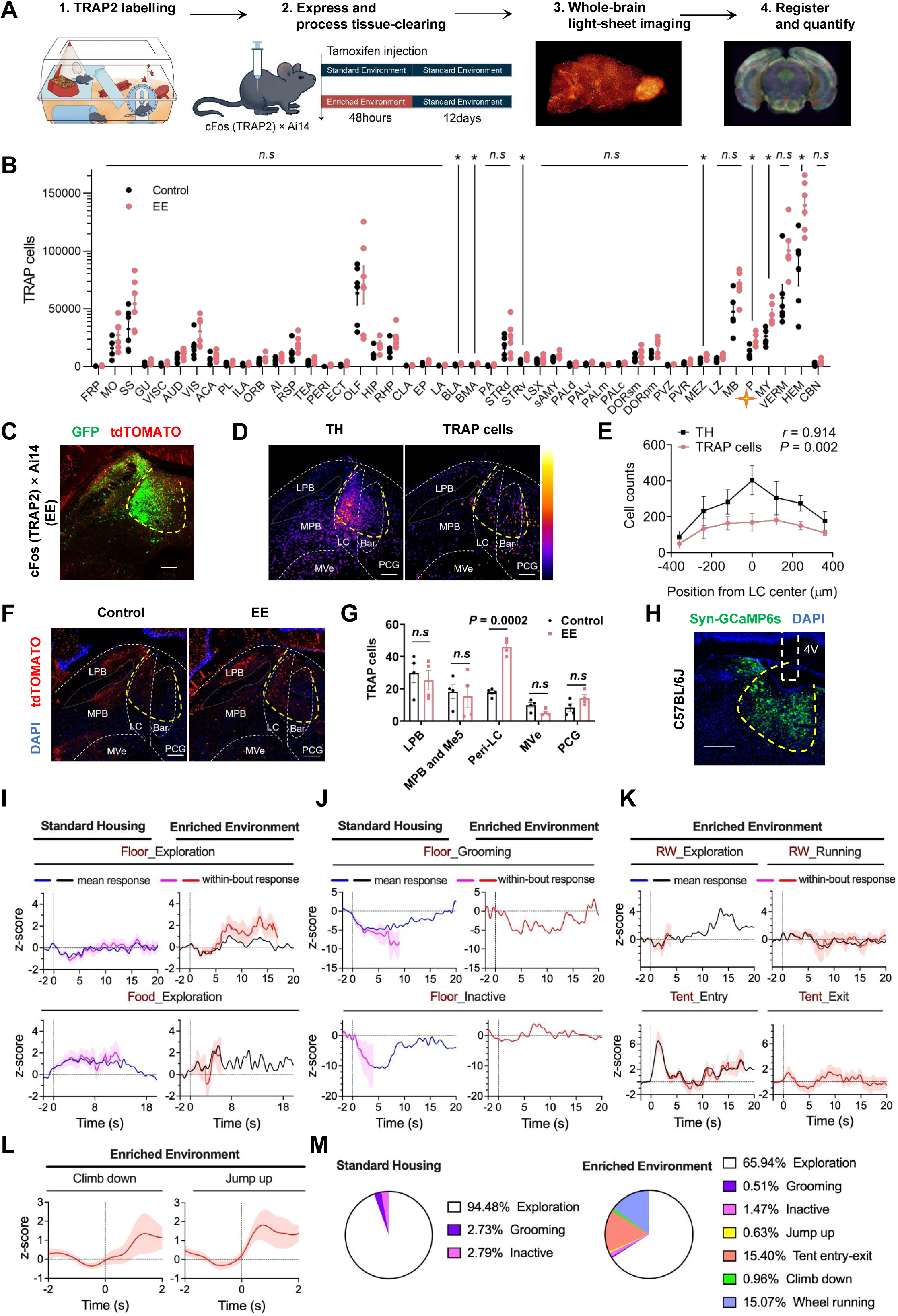
Activation of peri-LC neuronal populations by environmental enrichment. (A) Schematic of whole-brain activity mapping to identify regions activated by standard and enriched environments. Double-transgenic cFos (TRAP2) × Ai14 reporter mice were injected with tamoxifen and housed under standard or enriched environments for 48 hours. After 12 days of tdTomato reporter gene expression, mice were euthanized and processed with tissue-clearing. Whole brains were imaged using a light-sheet microscope, followed by automated registration to the Allen Brain Reference Atlas, cell segmentation, and quantification to identify brain regions with differential accumulation of TRAP-labeled (tdTomato^+^) cells. (B) Regional TRAP cell counts of enriched (red) versus standard environment (black) cohorts sorted from anterior to posterior in all anatomical regions (n = 6 per group, multiple unpaired *t*-tests corrected for multiple comparisons using the two-stage linear step-up procedure of Benjamini, Krieger, and Yekutieli, *false discovery rate (FDR) = 10%). Adjusted *p*-values: FRP *p* = 0.19878, MO *p* = 0.10801, SS *p* = 0.09162, GU *p* = 0.17950, VISC *p* = 0.07051, AUD *p* = 0.09133, VIS *p* = 0.09162, ACA *p* = 0.15636, PL *p* = 0.19878, ILA *p* = 0.25312, ORB *p* = 0.10801, AI *p* = 0.09162, RSP *p* = 0.09162, TEA *p* = 0.10801, PERI *p* = 0.11258, ECT *p* = 0.09133, OLF *p* = 0.46888, HIP *p* = 0.05196, RHP *p* = 0.07554, CLA *p* = 0.24852, EP *p* = 0.05177, LA *p* = 0.09133, BLA *p* = 0.04160, BMA *p* = 0.04160, PA *p* = 0.05177, STRd *p* = 0.19878, STRv *p* = 0.04160, LSX *p* = 0.20968, sAMY *p* = 0.07077, PALd *p* = 0.39897, PALv *p* = 0.05177, PALm *p* = 0.09162, PALc *p* = 0.05177, DORsm *p* = 0.15636, DORpm *p* = 0.05787, PVZ *p* = 0.05787, PVR *p* = 0.11012, MEZ *p* = 0.04160, LZ *p* = 0.05177, MB *p* = 0.05177, P *p* = 0.04160, MY *p* = 0.04160, VERM *p* = 0.05177, HEM *p* = 0.04160, CBN *p* = 0.05787. Acronyms for each brain region are listed in Figure S1E and Table 1. (C), cFos (TRAP2) × Ai14 mice (TRAP-labeled in enriched environments) were injected with AAV-TH-GFP virus, and coronal brain sections were subsequently collected. Scale bar, 200 μm. (D), Map of density of TH and activated TRAP (tdTomato^+^) neurons within a 360 μm radius around the LC region (anterior to posterior projection). n = 3. Scale bar, 200 μm. (E) Distributions of TH and activated TRAP (tdTomato^+^) cells within a 360 μm radius around the LC region from posterior to anterior. n = 3. (F) Representative coronal section showing activated TRAP (tdTomato^+^) cells in peri-LC (yellow dotted line) in mice exposed to standard or enriched environments. Scale bar, 200 μm. (G) Quantification of activated TRAP (tdTomato^+^) in peri-LC and surrounding area. n = 4. (H) Representative images of viral injection and optical fiber implantation in the peri-LC. Scale bar, 200 μm. (I–L) Peri-LC calcium responses aligned to the onset of indicated behaviors in standard housing (SH) and enriched environment (EE) mice. Blue and black traces indicate the mean event-aligned responses in SH and EE mice, respectively, whereas purple and red traces indicate the within-bout responses in SH and EE mice, respectively. Mean response was calculated as the average calcium signal within a fixed 20-s window after behavioral onset. Within-bout response was calculated using only the period during which the corresponding behavior was continuously expressed. Thus, the mean response reflects the overall onset-associated response, whereas the within-bout response reflects calcium activity during the actual behavioral bout. Shaded areas indicate SEM, shown only for the within-bout responses. The vertical dashed line marks behavior onset. (I) Floor exploration or food-top exploration: SH floor exploration, 87 trials, n = 7; EE floor exploration, 113 trials, n = 7; SH food-top exploration, 69 trials, n = 7; EE food-top exploration, 29 trials, n = 7. (J) Grooming or inactive state: SH grooming, 6 trials, n = 5; EE grooming, 1 trial, n = 1; SH inactive state, 3 trials, n = 2; EE inactive state, 1 trial, n = 1. (K) Running wheel (RW) exploration, 15 trials, n = 5; RW running, 18 trials, n = 5; tent entry, 24 trials, n = 7; tent exit, 22 trials, n = 7. (L) Climb down, 26 trials, n = 7; jump up, 21 trials, n = 7. (M) Pie charts showing the distribution of annotated behavioral events during calcium photometry recordings in SH and EE mice. Percentages indicate the proportion of each behavioral category among all annotated events. n = 7.

**Table 1:**
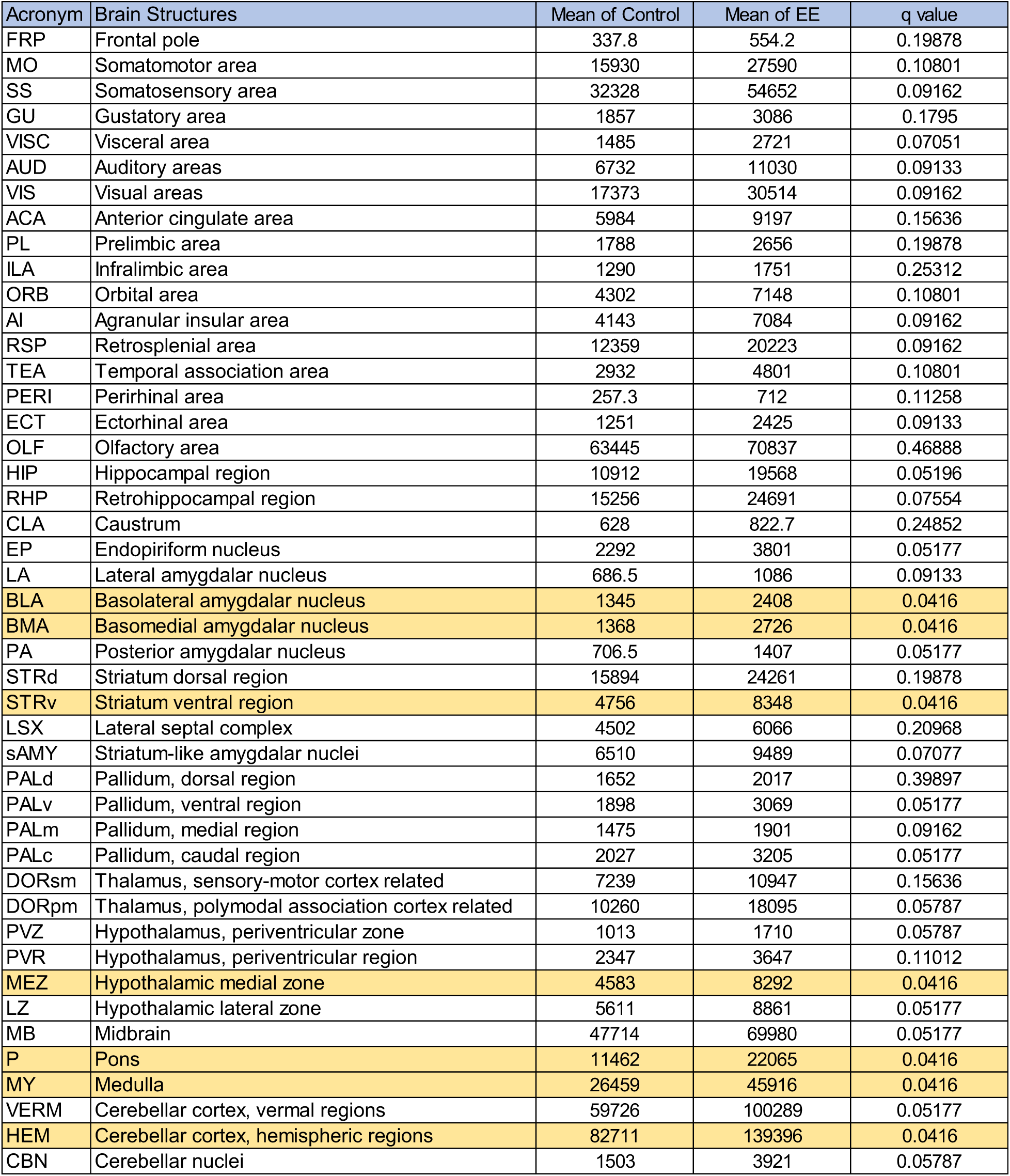
Brain-wide analysis of neural activation in EE.

| Acronym | Brain Structures | Mean of Control | Mean of EE | q value |
| --- | --- | --- | --- | --- |
| FRP | Frontal pole | 337.8 | 554.2 | 0.19878 |
| MO | Somatomotor area | 15930 | 27590 | 0.10801 |
| SS | Somatosensory area | 32328 | 54652 | 0.09162 |
| GU | Gustatory area | 1857 | 3086 | 0.1795 |
| VISC | Visceral area | 1485 | 2721 | 0.07051 |
| AUD | Auditory areas | 6732 | 11030 | 0.09133 |
| VIS | Visual areas | 17373 | 30514 | 0.09162 |
| ACA | Anterior cingulate area | 5984 | 9197 | 0.15636 |
| PL | Prelimbic area | 1788 | 2656 | 0.19878 |
| ILA | Infralimbic area | 1290 | 1751 | 0.25312 |
| ORB | Orbital area | 4302 | 7148 | 0.10801 |
| AI | Agranular insular area | 4143 | 7084 | 0.09162 |
| RSP | Retrosplenial area | 12359 | 20223 | 0.09162 |
| TEA | Temporal association area | 2932 | 4801 | 0.10801 |
| PERI | Perirhinal area | 257.3 | 712 | 0.11258 |
| ECT | Ectorhinal area | 1251 | 2425 | 0.09133 |
| OLF | Olfactory area | 63445 | 70837 | 0.46888 |
| HIP | Hippocampal region | 10912 | 19568 | 0.05196 |
| RHP | Retrohippocampal region | 15256 | 24691 | 0.07554 |
| CLA | Caustrum | 628 | 822.7 | 0.24852 |
| EP | Endopiriform nucleus | 2292 | 3801 | 0.05177 |
| LA | Lateral amygdalar nucleus | 686.5 | 1086 | 0.09133 |
| BLA | Basolateral amygdalar nucleus | 1345 | 2408 | 0.0416 |
| BMA | Basomedial amygdalar nucleus | 1368 | 2726 | 0.0416 |
| PA | Posterior amygdalar nucleus | 706.5 | 1407 | 0.05177 |
| STRd | Striatum dorsal region | 15894 | 24261 | 0.19878 |
| STRv | Striatum ventral region | 4756 | 8348 | 0.0416 |
| LSX | Lateral septal complex | 4502 | 6066 | 0.20968 |
| sAMY | Striatum-like amygdalar nuclei | 6510 | 9489 | 0.07077 |
| PALd | Pallidum, dorsal region | 1652 | 2017 | 0.39897 |
| PALv | Pallidum, ventral region | 1898 | 3069 | 0.05177 |
| PALm | Pallidum, medial region | 1475 | 1901 | 0.09162 |
| PALc | Pallidum, caudal region | 2027 | 3205 | 0.05177 |
| DORsm | Thalamus, sensory-motor cortex related | 7239 | 10947 | 0.15636 |
| DORpm | Thalamus, polymodal association cortex related | 10260 | 18095 | 0.05787 |
| PVZ | Hypothalamus, periventricular zone | 1013 | 1710 | 0.05787 |
| PVR | Hypothalamus, periventricular region | 2347 | 3647 | 0.11012 |
| MEZ | Hypothalamic medial zone | 4583 | 8292 | 0.0416 |
| LZ | Hypothalamic lateral zone | 5611 | 8861 | 0.05177 |
| MB | Midbrain | 47714 | 69980 | 0.05177 |
| P | Pons | 11462 | 22065 | 0.0416 |
| MY | Medulla | 26459 | 45916 | 0.0416 |
| VERM | Cerebellar cortex, vermal regions | 59726 | 100289 | 0.05177 |
| HEM | Cerebellar cortex, hemispheric regions | 82711 | 139396 | 0.0416 |
| CBN | Cerebellar nuclei | 1503 | 3921 | 0.05787 |

The locus coeruleus (LC), a compact nucleus in the pons of the brainstem, participates in the regulation of arousal and wakefulness, sensory and emotional processing, cognitive functions, and autonomic responses ^22^. Whole-brain cfosTRAPed cell mapping (Figure 1B, the orange four-point stars labeled ‘P’ (Pon) denote the brain region where the LC and peri-LC are located) and functional analyses of brain regions identified the LC as central to the neural adaptations observed in this study. To label the precise location of LC-noradrenergic (NA) neurons, tyrosine hydroxylase (TH)-GFP virus was injected into EE-labeled transgenic TRAP2 × Ai14 mice, with subsequent examination of coronal sections of the entire LC (Figure 1C and Figure S1H). Quantification of neuronal density in the LC and peri-LC region revealed a significant positive correlation between c-Fos-positive and TH-positive neurons (Figures 1D and 1E). Environmental enrichment led to a marked increase in the density of c-Fos-positive neurons in the peri-LC compared to standard housing conditions (Figures 1C, 1D, 1F, yellow dashed circle, and 1G).

The peri-LC, defined as the region surrounding the anterior LC (Shipley et al., 1996), is anatomically flanked laterally by the mesencephalic trigeminal nucleus (Me5), medially by the laterodorsal tegmental nucleus (LDT), and dorsally by the fourth ventricle ^6^. In our study, environmental enrichment-induced activation of the peri-LC encompassed both the LC and areas innervated by LC-TH neuronal terminals (Figure 1C and D, yellow dashed circle). For clarity, the term peri-LC is used to refer to this combined region. *In situ* hybridization revealed that neurons in the peri-LC expressed markers for TH, VGLUT, and GABA (Figure S4E and F).

Under environmentally enriched conditions, the number of active TH- and GABA-expressing neurons significantly increased, while VGLUT-expressing neurons showed no notable change (Figure S4E and F). GABAergic neurons in the peri-LC are known to regulate arousal by modulating LC-TH neuronal activity ^23^. These findings suggest that the peri-LC modulates brain states in response to enriched environments, enhancing adaptability to external stimuli.

To determine whether the elevated peri-LC neuronal activity under enriched environment conditions was transient or sustained, and to investigate which environmental factors might contribute to this increase, we performed *in vivo* calcium photometry. Neural activity in the peri-LC was measured in wild-type mice at 6 weeks of age (Figure 1H). Compared to mice housed under standard environments (stand housing, SH), mice exposed to enriched environments exhibited significantly elevated peri-LC neural activity (Figures 1I and 1M). The results showed that exploration on or around the food was associated with increased peri-LC calcium activity in both SH and EE conditions (Figure 1I, *lower*). During floor exploration, peri-LC activity showed an initial transient decrease within the first 5 s after behavioral onset in both SH and EE. After this initial phase, however, peri-LC activity markedly increased in EE, resulting in an overall higher calcium response than that observed under SH conditions (Figure 1I, *upper*). Grooming was associated with a significant reduction in peri-LC calcium activity in both SH and EE conditions (Figure 1j, *upper*). During inactive periods, peri-LC activity was markedly reduced in SH. In EE, peri-LC activity also showed an initial decrease followed by a subsequent rebound, but the magnitude of suppression was much smaller than that observed in SH (Figure 1J, *lower*).

Under EE conditions, when mice jumped onto the running wheel (RW) and began exploring it, peri-LC calcium activity showed a modest transient decrease followed by an increase (Figure 1K, *upper*), a response pattern similar to that observed during floor exploration (Figure 1I, *upper,* RW_Exploration). When mice left the running wheel and transitioned to other locations or behaviors, peri-LC activity increased further (Figure 1I, *upper,* RW_Exploration). In contrast, when mice ran on the wheel for more than 4 s, peri-LC calcium activity was overall suppressed (Figure 1K, *upper,* RW_Running). Entry into or exit from the tent, particularly tent entry, was associated with a robust increase in peri-LC calcium activity (Figure 1K, *lower*, Tent entry and exit). In SH, mice rarely showed visually identifiable upward jumping or downward climbing due to the lack of spatial structure. In EE, however, both upward jumping and downward climbing were associated with increased peri-LC calcium activity (Figure 1L).

Videos 2-3 further illustrate the peri-LC calcium dynamics associated with running, tent entry and exit, jumping, exploration of confined spaces, and grooming. Taken together, these event-aligned analyses suggest that increased spatial and structural complexity in the EE may be a key factor contributing to elevated peri-LC neuronal activity. Based on the data, together with the proportion of time spent in different behavioral states under each housing condition (Figure 1M), peri-LC neuronal activity is overall higher in EE than in SH.

### Enriched environments modulate endothelial cell heterogeneity and enhance bone density

To investigate whether enriched environments coordinate osteogenic CD31^hi^EMCN^hi^ endothelium to maintain bone formation, the levels of these endothelial cells were assessed in mice exposed to environmentally enriched conditions. Mice exposed to enriched environments displayed significantly enhanced bone density compared to standard controls (Figure 2E and F *lower*). CD31^hi^EMCN^hi^ endothelial cells were predominantly localized to the bone marrow beneath the growth plate and near the endosteum, consistent with prior findings ^18^. Notably, the abundance of CD31^+^EMCN^+^ endothelial cells was significantly higher in EE-exposed mice compared to the standard controls (Figures 2A-C). Although endothelial cell heterogeneity markers differ across organs, analysis of CD31^+^EMCN^+^ endothelial cells in the brain, heart, kidney, liver, and lung revealed no significant changes (Figure S2E), indicating that the effect on CD31 and EMCN expression is specific to the skeletal system. Even within skeletal sites such as the vertebrae and skull, CD31^+^EMCN^+^ vessel levels showed no significant differences between EE-housed and control mice (Figure S2E). However, flow cytometry confirmed the pronounced increase in CD31^hi^EMCN^hi^ vascular endothelium in the long bones of EE-housed mice compared to controls (Figures 2D and 2F *upper*). CD31^hi^EMCN^hi^ endothelium promotes bone formation through the release of angiocrine factors. Noggin has been identified as a relevant endothelial-derived angiocrine factor ^24^. We therefore examined Noggin expression in the long bones of mice exposed to enriched environment. Compared with mice housed under standard conditions, EE mice showed a significant increase in Noggin signal within and around the vasculature (Fig. S2A and Fig. S2D, *left*). These findings indicate that environmental enrichment selectively regulates skeletal CD31^hi^EMCN^hi^ endothelial cells and enhances the release of angiocrine factors, thereby increasing long bone density. Consistent with the osteogenic role of CD31^hi^EMCN^hi^ endothelium ^18,20^, mice exposed to enriched environment for four weeks showed significant increases in BV/TV, Ct.Th, Tb.Th, and Tb.N, whereas Tb.Sp showed a non-significant decreasing trend (Figure 2F and Table 2). These findings indicate that enriched environment increases long-bone mass and improves bone microarchitecture, consistent with enhanced bone accrual associated with increased CD31^hi^EMCN^hi^ endothelium.

**Table 2-19:** Micro-CT Analysis of Bone Structural Parameters. Table 2:

| C57BL/6J Standard housing vs. Enriched environment |  |  |  |  |  |  |  |  |
| --- | --- | --- | --- | --- | --- | --- | --- | --- |
| Group | SH | EE | SH | EE |  |  |  |  |
| Descriptive statistics | Mean |  | SEM |  | n | Direction vs. control | P value | Significance |
| Ct.Th (mm) | 0.2162 | 0.2276 | 0.003098 | 0.003648 | 22 | ↑ | 0.0216 | * |
| Tb.Th (mm) | 0.08964 | 0.09767 | 0.001951 | 0.001633 | 22 | ↑ | 0.0029 | ** |
| Tb.N (1/mm) | 3.265 | 3.537 | 0.07693 | 0.07837 | 22 | ↑ | 0.0175 | * |
| Tb.Sp (mm) | 0.1845 | 0.1775 | 0.004411 | 0.004445 | 22 | ↓ | 0.2699 | n.s |

**Table 3:**

| Cfos (TRAP2) Peri-LC Control vs. DTR |  |  |  |  |  |  |  |  |
| --- | --- | --- | --- | --- | --- | --- | --- | --- |
| Group | Control | DTR | Control | DTR |  |  |  |  |
| Descriptive statistics | Mean |  | SEM |  | n | Direction vs. control | P value | Significance |
| Ct.Th (mm) | 0.2265 | 0.2158 | 0.005426 | 0.004691 | 8-9 | ↓ | 0.156 | n.s |
| Tb.Th (mm) | 0.09913 | 0.08787 | 0.005158 | 0.002505 | 8-9 | ↓ | 0.0598 | n.s |
| Tb.N (1/mm) | 3.329 | 2.903 | 0.07541 | 0.1212 | 8-9 | ↓ | 0.0111 | * |
| Tb.Sp (mm) | 0.1796 | 0.1979 | 0.003741 | 0.005077 | 8-9 | ↑ | 0.0122 | * |

**Table 4:**

| Cfos (TRAP2) Peri-LC Control vs. hM3Dq |  |  |  |  |  |  |  |  |
| --- | --- | --- | --- | --- | --- | --- | --- | --- |
| Group | Control | hM3Dq | Control | hM3Dq |  |  |  |  |
| Descriptive statistics | Mean |  | SEM |  | n | Direction vs. control | P value | Significance |
| Ct.Th (mm) | 0.2299 | 0.2426 | 0.004986 | 0.00363 | 11-13 | ↑ | 0.0578 | n.s |
| Tb.Th (mm) | 0.08921 | 0.1054 | 0.002307 | 0.001747 | 11-13 | ↑ | <0.0001 | **** |
| Tb.N (1/mm) | 3.217 | 4.064 | 0.1336 | 0.1086 | 11-13 | ↑ | <0.0001 | **** |
| Tb.Sp (mm) | 0.1807 | 0.1543 | 0.004495 | 0.004066 | 11-13 | ↓ | 0.0003 | *** |

**Table 5:**

| Cfos (TRAP2) Peri-LC Control vs . hM4Di |  |  |  |  |  |  |  |  |
| --- | --- | --- | --- | --- | --- | --- | --- | --- |
| Group | Control | hM4Di | Control | hM4Di |  |  |  |  |
| Descriptive statistics | Mean |  | SEM |  | n | Direction vs. control | P value | Significance |
| Ct.Th (mm) | 0.223 | 0.228 | 0.007396 | 0.002763 | 6 | ↑ | 0.5342 | *n.s* |
| Tb.Th (mm) | 0.08732 | 0.07604 | 0.00338 | 0.001276 | 6 | ↓ | 0.0108 | * |
| Tb.N (mm) | 4.158 | 2.993 | 0.1034 | 0.1088 | 6 | ↓ | <0.0001 | **** |
| Tb.Sp (mm) | 0.1526 | 0.1925 | 0.005555 | 0.006024 | 6 | ↑ | 0.0007 | *** |

**Table 6:**

| Cfos (TRAP2) RVLM Control vs . DTR |  |  |  |  |  |  |  |  |
| --- | --- | --- | --- | --- | --- | --- | --- | --- |
| Group | Control | DTR | Control | DTR |  |  |  |  |
| Descriptive statistics | Mean |  | SEM |  | n | Direction vs. control | P value | Significance |
| Ct.Th (mm) | 0.2355 | 0.2269 | 0.004974 | 0.007766 | 9 | ↓ | 0.3651 | *n.s* |
| Tb.Th (mm) | 0.09522 | 0.09374 | 0.002861 | 0.002461 | 9 | ↓ | 0.7001 | *n.s* |
| Tb.N (1/mm) | 3.684 | 3.434 | 0.08619 | 0.1593 | 9 | ↓ | 0.1861 | *n.s* |
| Tb.Sp (mm) | 0.1635 | 0.1717 | 0.003217 | 0.006309 | 9 | ↑ | 0.2619 | *n.s* |

**Table 7:**

| Cfos (TRAP2) RVLM Control vs . hM3Dq |  |  |  |  |  |  |  |  |
| --- | --- | --- | --- | --- | --- | --- | --- | --- |
| Group | Control | hM3Dq | Control | hM3Dq |  |  |  |  |
| Descriptive statistics | Mean |  | SEM |  | n | Direction vs. control | P value | Significance |
| Ct.Th (mm) | 0.2269 | 0.2136 | 0.004094 | 0.003376 | 9 | ↓ | 0.0233 | * |
| Tb.Th (mm) | 0.08992 | 0.08482 | 0.001562 | 0.001676 | 9 | ↓ | 0.0406 | * |
| Tb.N (1/mm) | 3.258 | 2.67 | 0.1384 | 0.1022 | 9 | ↓ | 0.0035 | ** |
| Tb.Sp (mm) | 0.1772 | 0.1962 | 0.006211 | 0.004938 | 9 | ↑ | 0.0289 | * |

**Table 8:**

| C57BL/6J Peri-LC Control vs. hM3Dq |  |  |  |  |  |  |  |  |
| --- | --- | --- | --- | --- | --- | --- | --- | --- |
| Group | Control | hM3Dq | Control | hM3Dq |  |  |  |  |
| Descriptive statistics | Mean |  | SEM |  | n | Direction vs. control | P value | Significance |
| Ct.Th (mm) | 0.2242 | 0.2351 | 0.003878 | 0.005157 | 8-10 | ↑ | 0.1017 | *n.s* |
| Tb.Th (mm) | 0.09053 | 0.09907 | 0.002782 | 0.001917 | 8-10 | ↑ | 0.029 | * |
| Tb.N (1/mm) | 3.255 | 3.74 | 0.1066 | 0.1114 | 8-10 | ↑ | 0.0066 | ** |
| Tb.Sp (mm) | 0.1815 | 0.1675 | 0.00475 | 0.00352 | 8-10 | ↓ | 0.0389 | * |

**Table 9:**

| GAD2-Cre Peri-LC Control vs. DTR |  |  |  |  |  |  |  |  |
| --- | --- | --- | --- | --- | --- | --- | --- | --- |
| Group | Control | DTR | Control | DTR |  |  |  |  |
| Descriptive statistics | Mean |  | SEM |  | n | Direction vs. control | P value | Significance |
| Ct.Th (mm) | 0.2274 | 0.2298 | 0.004017 | 0.004138 | 11-12 | ↑ | 0.6875 | *n.s* |
| Tb.Th (mm) | 0.1046 | 0.09852 | 0.005136 | 0.002119 | 11-12 | ↓ | 0.2748 | *n.s* |
| Tb.N (1/mm) | 3.535 | 3.549 | 0.1841 | 0.1248 | 11-12 | ↑ | 0.9497 | *n.s* |
| Tb.Sp (mm) | 0.1706 | 0.1746 | 0.006619 | 0.004824 | 11-12 | ↑ | 0.617 | *n.s* |

**Table 10:**

| GAD2-Cre Peri-LC Control vs. hM3Dq |  |  |  |  |  |  |  |  |
| --- | --- | --- | --- | --- | --- | --- | --- | --- |
| Group | Control | hM3Dq | Control | hM3Dq |  |  |  |  |
| Descriptive statistics | Mean |  | SEM |  | n | Direction vs. control | P value | Significance |
| Ct.Th (mm) | 0.2075 | 0.2158 | 0.005684 | 0.004631 | 7-8 | ↑ | 0.2734 | *n.s* |
| Tb.Th (mm) | 0.09148 | 0.09643 | 0.003252 | 0.004032 | 7-8 | ↑ | 0.3653 | *n.s* |
| Tb.N (1/mm) | 3.111 | 3.162 | 0.1782 | 0.176 | 7-8 | ↑ | 0.8431 | *n.s* |
| Tb.Sp (mm) | 0.1916 | 0.1857 | 0.006306 | 0.007469 | 7-8 | ↓ | 0.5625 | *n.s* |

**Table 11:**

| TH-Cre Peri-LC Control vs . DTR |  |  |  |  |  |  |  |  |
| --- | --- | --- | --- | --- | --- | --- | --- | --- |
| Group | Control | DTR | Control | DTR |  |  |  |  |
| Descriptive statistics | Mean |  | SEM |  | n | Direction vs. control | P value | Significance |
| Ct.Th (mm) | 0.2282 | 0.2162 | 0.005742 | 0.002127 | 8-9 | ↓ | 0.0568 | *n.s* |
| Tb.Th (mm) | 0.09832 | 0.08648 | 0.002919 | 0.001587 | 8-9 | ↓ | 0.0023 | **\*\*** |
| Tb.N (1/mm) | 3.402 | 2.43 | 0.311 | 0.2076 | 8-9 | ↓ | 0.0181 | **\*** |
| Tb.Sp (mm) | 0.1816 | 0.2252 | 0.01318 | 0.009798 | 8-9 | ↑ | 0.0167 | **\*** |

**Table 12:**

| TH-Cre Peri-LC Control vs . hM3Dq |  |  |  |  |  |  |  |  |
| --- | --- | --- | --- | --- | --- | --- | --- | --- |
| Group | Control | hM3Dq | Control | hM3Dq |  |  |  |  |
| Descriptive statistics | Mean |  | SEM |  | n | Direction vs. control | P value | Significance |
| Ct.Th (mm) | 0.217 | 0.224 | 0.002341 | 0.003805 | 9-10 | ↑ | 0.1441 | *n.s* |
| Tb.Th (mm) | 0.08637 | 0.09092 | 0.001046 | 0.001976 | 9-10 | ↑ | 0.0655 | *n.s* |
| Tb.N (1/mm) | 2.795 | 2.894 | 0.1353 | 0.1485 | 9-10 | ↑ | 0.631 | *n.s* |
| Tb.Sp (mm) | 0.2012 | 0.204 | 0.008326 | 0.01008 | 9-10 | ↑ | 0.835 | *n.s* |

**Table 13:**

| Vglut2-Cre Peri-LC Control vs . DTR |  |  |  |  |  |  |  |  |
| --- | --- | --- | --- | --- | --- | --- | --- | --- |
| Group | Control | DTR | Control | DTR |  |  |  |  |
| Descriptive statistics | Mean |  | SEM |  | n | Direction vs. control | P value | Significance |
| Ct.Th (mm) | 0.2301 | 0.2177 | 0.006232 | 0.005458 | 9 | ↓ | 0.1537 | *n.s* |
| Tb.Th (mm) | 0.09617 | 0.09398 | 0.00255 | 0.002337 | 9 | ↓ | 0.5356 | *n.s* |
| Tb.N (1/mm) | 3.689 | 3.644 | 0.1704 | 0.1528 | 9 | ↓ | 0.8484 | *n.s* |
| Tb.Sp (mm) | 0.1707 | 0.1767 | 0.006495 | 0.007333 | 9 | ↑ | 0.5465 | *n.s* |

**Table 14:**

| Vglut2-Cre Peri-LC Control vs. hM3Dq |  |  |  |  |  |  |  |  |
| --- | --- | --- | --- | --- | --- | --- | --- | --- |
| Group | Control | hM3Dq | Control | hM3Dq |  |  |  |  |
| Descriptive statistics | Mean |  | SEM |  | n | Direction vs. control | P value | Significance |
| Ct.Th (mm) | 0.2299 | 0.225 | 0.004597 | 0.006025 | 9 | ↓ | 0.526 | *n.s* |
| Tb.Th (mm) | 0.09386 | 0.09463 | 0.001559 | 0.003226 | 9 | ↑ | 0.8336 | *n.s* |
| Tb.N (1/mm) | 3.295 | 3.195 | 0.1407 | 0.1103 | 9 | ↓ | 0.585 | *n.s* |
| Tb.Sp (mm) | 0.1841 | 0.1874 | 0.005017 | 0.005641 | 9 | ↑ | 0.6664 | *n.s* |

**Table 15:**
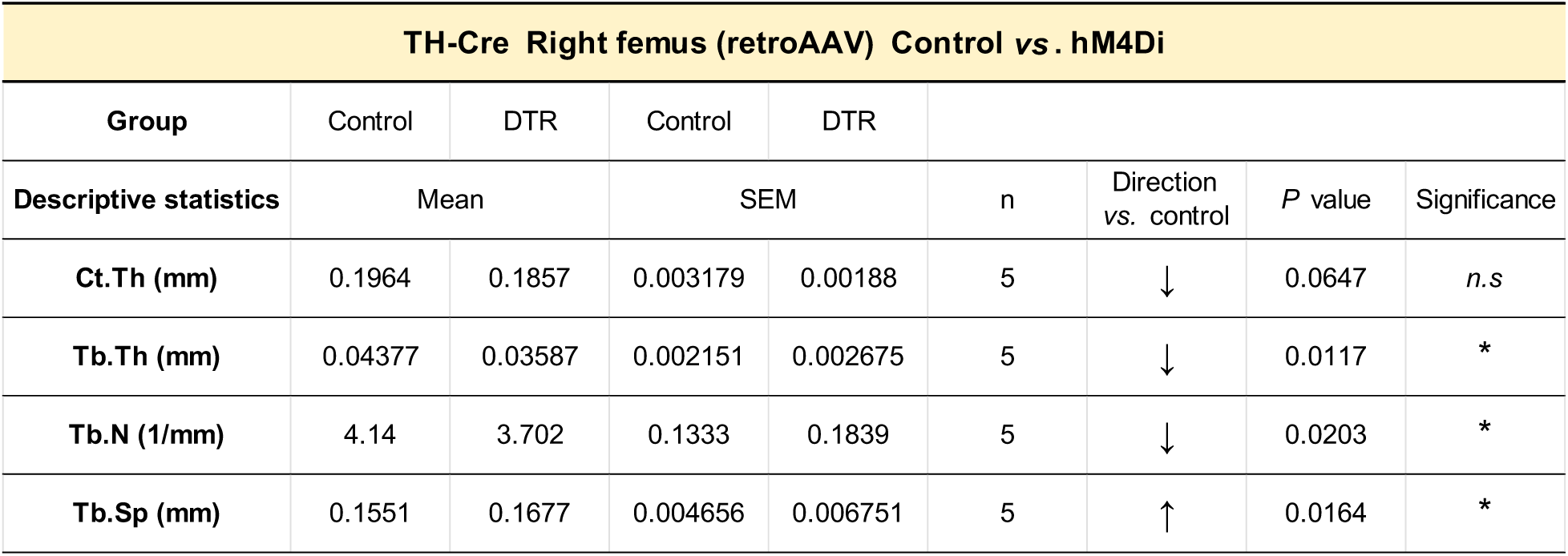

| TH-Cre Right femus (retroAAV) Control vs. hM4Di |  |  |  |  |  |  |  |  |
| --- | --- | --- | --- | --- | --- | --- | --- | --- |
| Group | Control | DTR | Control | DTR |  |  |  |  |
| Descriptive statistics | Mean |  | SEM |  | n | Direction vs. control | P value | Significance |
| Ct.Th (mm) | 0.1964 | 0.1857 | 0.003179 | 0.00188 | 5 | ↓ | 0.0647 | *n.s* |
| Tb.Th (mm) | 0.04377 | 0.03587 | 0.002151 | 0.002675 | 5 | ↓ | 0.0117 | * |
| Tb.N (1/mm) | 4.14 | 3.702 | 0.1333 | 0.1839 | 5 | ↓ | 0.0203 | * |
| Tb.Sp (mm) | 0.1551 | 0.1677 | 0.004656 | 0.006751 | 5 | ↑ | 0.0164 | * |

**Table 16:**

| TH-Cre Right femus (retroAAV) Control vs. hM3Dq |  |  |  |  |  |  |  |  |
| --- | --- | --- | --- | --- | --- | --- | --- | --- |
| Group | Control | hM3Dq | Control | hM3Dq |  |  |  |  |
| Descriptive statistics | Mean (mm) |  | SEM |  | n | Direction vs. control | P value | Significance |
| Ct.Th (mm) | 0.1841 | 0.1759 | 0.003357 | 0.007452 | 5 | ↓ | 0.2299 | *n.s* |
| Tb.Th (mm) | 0.0971 | 0.1071 | 0.001906 | 0.003318 | 5 | ↑ | 0.0057 | * |
| Tb.N (1/mm) | 3.802 | 4.25 | 0.155 | 0.1258 | 5 | ↑ | 0.0305 | * |
| Tb.Sp (mm) | 0.161 | 0.1501 | 0.006755 | 0.004922 | 5 | ↓ | 0.101 | *n.s* |

**Table 17:**

| C57BL/6J ICI118551 |  |  |  |  |  |  |  |  |
| --- | --- | --- | --- | --- | --- | --- | --- | --- |
| Group | Control | DTR | Control | DTR |  |  |  |  |
| Descriptive statistics | Mean |  | SEM |  | n | Direction vs. control | P value | Significance |
| Ct.Th (mm) | 0.2249 | 0.2217 | 0.00245 | 0.004884 | 9 | ↓ | 0.5647 | *n.s* |
| Tb.Th (mm) | 0.1024 | 0.09551 | 0.001364 | 0.001916 | 9 | ↓ | 0.0101 | * |
| Tb.N (1/mm) | 3.657 | 3.38 | 0.1125 | 0.1289 | 9 | ↓ | 0.124 | *n.s* |
| Tb.Sp (mm) | 0.1768 | 0.1862 | 0.006858 | 0.006292 | 9 | ↑ | 0.3266 | *n.s* |

**Table 18:**

| Adrb2 flox Control vs. Cdh5-ERT2-Cre |  |  |  |  |  |  |  |  |
| --- | --- | --- | --- | --- | --- | --- | --- | --- |
| Group | Control | hM3Dq | Control | hM3Dq |  |  |  |  |
| Descriptive statistics | Mean (mm) |  | SEM |  | n | Direction vs. control | P value | Significance |
| Ct.Th (mm) | 0.2026 | 0.2096 | 0.009054 | 0.00857 | 5-6 | ↑ | 0.5887 | *n.s* |
| Tb.Th (mm) | 0.07458 | 0.06535 | 0.001748 | 0.001703 | 5-6 | ↓ | 0.0045 | ** |
| Tb.N (1/mm) | 3.016 | 2.188 | 0.1494 | 0.1328 | 5-6 | ↓ | 0.0025 | ** |
| Tb.Sp (mm) | 0.1964 | 0.2511 | 0.00844 | 0.009157 | 5-6 | ↑ | 0.0019 | ** |

**Table 19:**

| Adrb2 flox Femus (AAV) OC-Cre vs. Tie2-Cre |  |  |  |  |  |  |  |  |
| --- | --- | --- | --- | --- | --- | --- | --- | --- |
| Group | Control | DTR | Control | DTR |  |  |  |  |
| Descriptive statistics | Mean |  | SEM |  | n | Direction vs. control | P value | Significance |
| Ct.Th (mm) | 0.2422 | 0.2432 | 0.007647 | 0.01177 | 7 | ↑ | 0.9163 | *n.s* |
| Tb.Th (mm) | 0.08985 | 0.08973 | 0.002415 | 0.003334 | 7 | ↓ | 0.9589 | *n.s* |
| Tb.N (1/mm) | 3.742 | 3.48 | 0.107 | 0.1082 | 7 | ↓ | 0.0054 | ** |
| Tb.Sp (mm) | 0.1678 | 0.1762 | 0.004414 | 0.00416 | 7 | ↑ | 0.1204 | *n.s* |

**Figure 2:**
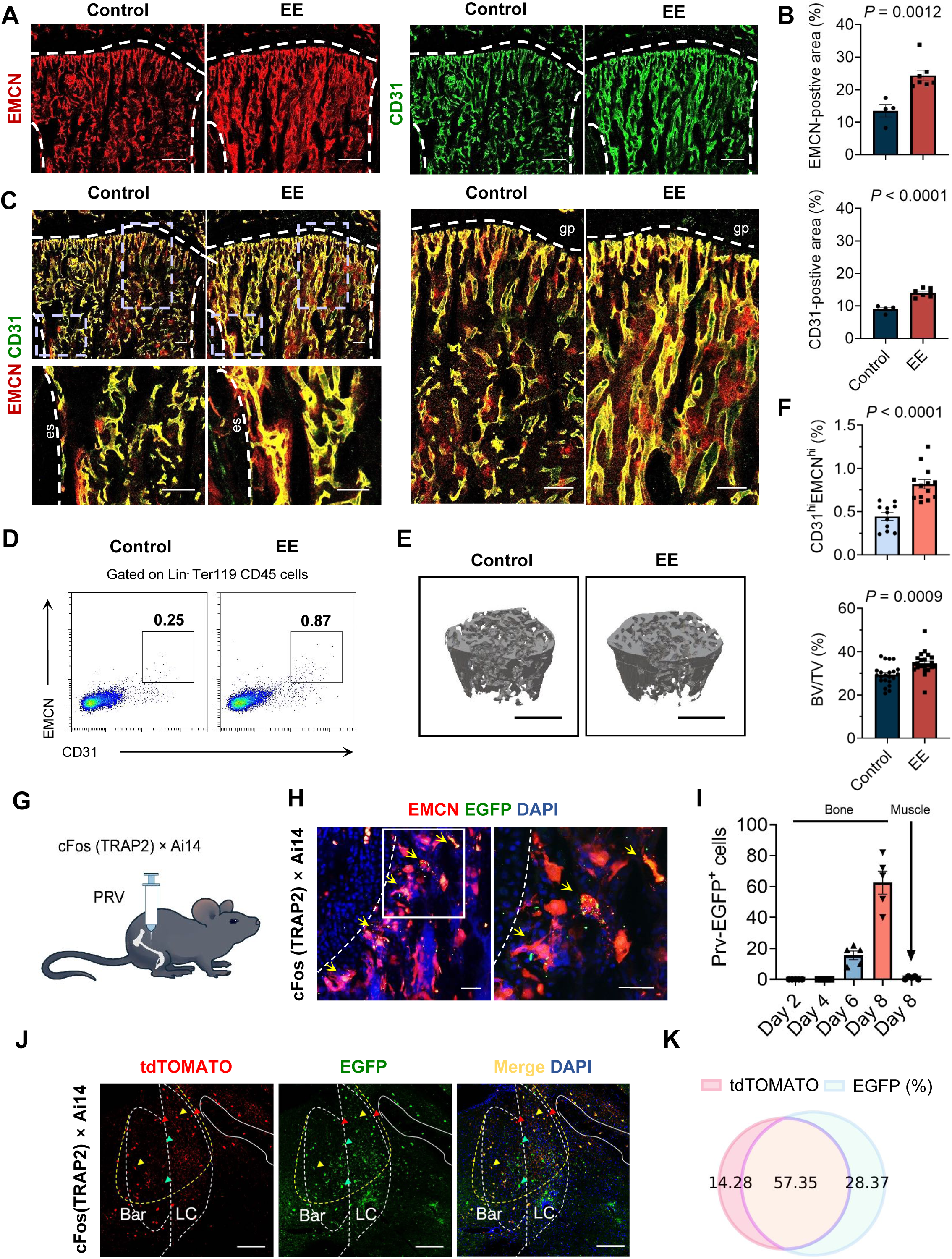
Regulation of endothelial heterogeneity and promotion of bone formation by enriched environments, and transsynaptic tracing from femur with PRV. (A) Representative confocal images of 7-week-old male mouse femurs stained with EMCN (left, red) and CD31 (right, green), EE-group mice were housed in an enriched environment for 4 weeks (3–7 weeks of age). Growth plate and cortical bone are marked with a dashed line. Scale bars, 200 μm. (B) Quantification of relative EMCN^+^ (*upper*) and CD31^+^ (*lower*) vessel area in bone marrow cavity of femur sections in 7-week-old standard- and EE-housed mice. n = 4–7. (C) Representative images of CD31 (green) and EMCN (red) dual-immunostained femur sections of 7-week-old male mice. Growth plate (gp) and endosteum (es) are marked. Scale bars, 100 μm. (D) and (F) *upper*, Representative flow cytometry plots (D) with quantification (F u*pper*) of CD31^hi^EMCN^hi^ endothelial cells from femurs and tibiae of 7-week-old standard- and EE-housed wild-type mice. n = 11–13. (E) and (F) *lower*, Representative μCT images of trabecular bone in distal femur (E) and bone volume/total volume (BV/TV) (F l*ower*) of 7-week-old standard- and EE-housed male wild-type mice. Additional micro-CT parameters, including Ct.Th, Tb.Th, Tb.N, and Tb.Sp, are provided in Table 2. n = 22. Scale bars, 1 mm. (G) Schematic representation of transsynaptic tracing from femur infected with PRV in EE-activated cFos (TRAP2) × Ai14 mice. (H) Representative images of metaphysis of PRV-infected femur stained with EMCN, growth plate is marked. Scale bars, 50 μm. (I) Quantification of PRV-tracing cells in the peri-LC of PRV-infected mice at series time points. n = 5. (J) Representative images of the peri-LC in EE-treated cFos (TRAP2) × Ai14 mice with PRV-infected femur. Scale bars, 200 μm. (K) Quantification of colocalization between femur-PRV-tracing and EE-activated cells in the peri-LC. n = 3

Physical activity, particularly weight-bearing and resistance exercise, is known to increase or maintain bone density ^25,26^. Because the running wheel in the EE cage increases locomotor activity relative to standard housing, we designed a series of experiments to determine whether locomotor activity itself accounted for the EE-induced increase in bone density.

A locked-wheel condition was established to control for the potential contribution of voluntary exercise. Mice were housed under the corresponding conditions for 4 weeks, and locking the running wheel in the EE cage did not prevent the EE-induced increases in CD31^hi^EMCN^hi^ endothelial cells or bone mass. Both parameters remained comparable to those observed under the full EE condition (Figure S3A, EE-LW). An empty EE cage containing only food and water was also included in our experiments. The results showed that simply increasing cage space was not sufficient to increase CD31^hi^EMCN^hi^ endothelial cells or bone mass (Figure S3, Empty-EE).

To determine whether locomotor activity was associated with bone mineral density in our study, we recorded nighttime videos of mice, quantified the duration of active behaviors, and correlated these measurements with bone mineral density. The results showed that the duration of active behavior during the sampling period was not correlated with bone mineral density after 4 weeks of housing under the corresponding conditions (Figure S3B). To exclude the potential influence of different housing environments on the analysis, we further examined the correlation between active time and bone mineral density within each housing condition separately. Consistently, no significant correlation was observed in any of the four housing conditions (Figure S3C). Moreover, active time during the sampling period did not differ significantly among mice housed under the four different conditions (Figure S3D).

These findings suggest that general locomotor activity, as defined by displacement, climbing, hanging, and wheel running during the active-phase sampling window, is unlikely to account for the EE-induced increase in bone mineral density. However, because wheel running in the enriched environment may represent a more intense form of exercise than general movement within the cage, we further quantified wheel-running activity using a wheel counter and designed a forced treadmill running experiment based on the wheel-running activity observed under the EE condition. The daily number of wheel revolutions was measured in individually housed mice, and the median value, 2,137 revolutions per day (Figure S3E), corresponding to approximately 1,200 m of running distance, was used as the reference daily exercise load for designing the forced treadmill running experiment. Mice were subjected to forced running for 4 weeks (6 days/week), from 4 to 8 weeks of age. The results showed that 4 weeks of forced running did not increase bone mineral density (Figure S3F).

Taken together, several lines of evidence argue against locomotor activity as the driver of the EE-induced skeletal phenotype. First, locking the running wheel did not prevent the increase in bone mineral density under EE conditions. Second, active time did not differ significantly among the four housing conditions, and active time during the sampling period was not correlated with bone mineral density. Third, 4 weeks of forced running which designed based on locomotor activity in EE failed to increase bone mineral density.

### Peri-LC regulates CD31^hi^EMCN^hi^ endothelium

To investigate whether enriched environments influence skeletal endothelial cells via the central nervous system, pseudorabies virus (PRV) was injected into the bone marrow cavity near the femoral growth plate in mice (Figure 2G). The PRV-infected mice exhibited morphological changes in vascular endothelial cells, and expression of PRV-encoded enhanced green fluorescent protein (EGFP) was detected (Figure 2H). Symptoms such as tremors appeared around day 7 post-injection. Analysis of PRV-EGFP expression in the peri-LC on days 2, 4, 6 and 8 revealed progressively increasing levels (Figure 2I). In contrast to bone marrow injections, which consistently labeled neurons in the peri-LC region, PRV tracing from skeletal muscle produced little to no labeling in peri-LC neurons, with only 0–2 labeled neurons detected per animal (Figure 2I and S4B). Notably, PRV tracing from both bone marrow and skeletal muscle resulted in labeling within the M1/M2 cortical areas, but the labeled regions within the motor cortex were spatially distinct between the two injection sites (Figure S4B). In cFosTRAP2 mice, which express tdTomato in response to enriched environments, PRV injections were administered post-EE exposure, followed by whole-brain section analysis. Co-expression of PRV-EGFP and tdTomato was observed in multiple brain regions, indicating neural connections and neuronal activation driven by environmental enrichment (Figure 4SA and Figure 2J). Among these regions, the peri-LC displayed the highest co-expression rate (57.35%) (Figure 2J and K), establishing it as a focal point of the study. Retrograde tracing experiments in TH-Cre × Ai14, Vglut2-Cre × Ai14, and Gad2-Cre × Ai14 mice confirmed that neurons projecting from the distal femurs to the peri-LC were predominantly TH-expressing (co-expression rate: 35.78%, TH-co-expression cells in total EGFP-expressed: 58.25%) and GABAergic neurons (co-expression rate: 17.73%, GAD2-co-expression cells in total EGFP-expressed: 42.45%) (Figure S4C and D).

In cFosTRAP2 mice, we injected an adeno-associated virus (AAV) carrying diphtheria toxin receptor (DTR) into the peri-LC to enable diphtheria toxin (DT)-mediated ablation of EE-labeled peri-LC neurons (Figure 3A). This ablation resulted in a marked decrease in CD31^hi^EMCN^hi^ endothelium and long-bone density in EE-exposed mice (Figure 3B–E). We next used chemogenetic manipulation based on designer receptors exclusively activated by designer drugs (DREADDs) to modulate the activity of EE-labeled peri-LC neurons. Chemogenetic inhibition using hM4Di, an inhibitory DREADD, reduced CD31^hi^EMCN^hi^ endothelium and long-bone density in EE-exposed mice (Figure S5E–I). Conversely, chemogenetic activation using hM3Dq, an excitatory DREADD, increased CD31^hi^EMCN^hi^ endothelium and bone density even under standard housing conditions (Figure 3F–J). All chemogenetic activation and inhibition experiments included matched CNO administration across experimental groups. In addition, we performed a separate control experiment to evaluate the effect of CNO alone in this system. In a control paradigm matched to the chemogenetic activation design, CNO injection alone did not alter bone endothelial heterogeneity, bone density, or other parameters measured in this study (Figure S5A–D).

**Figure 3:**
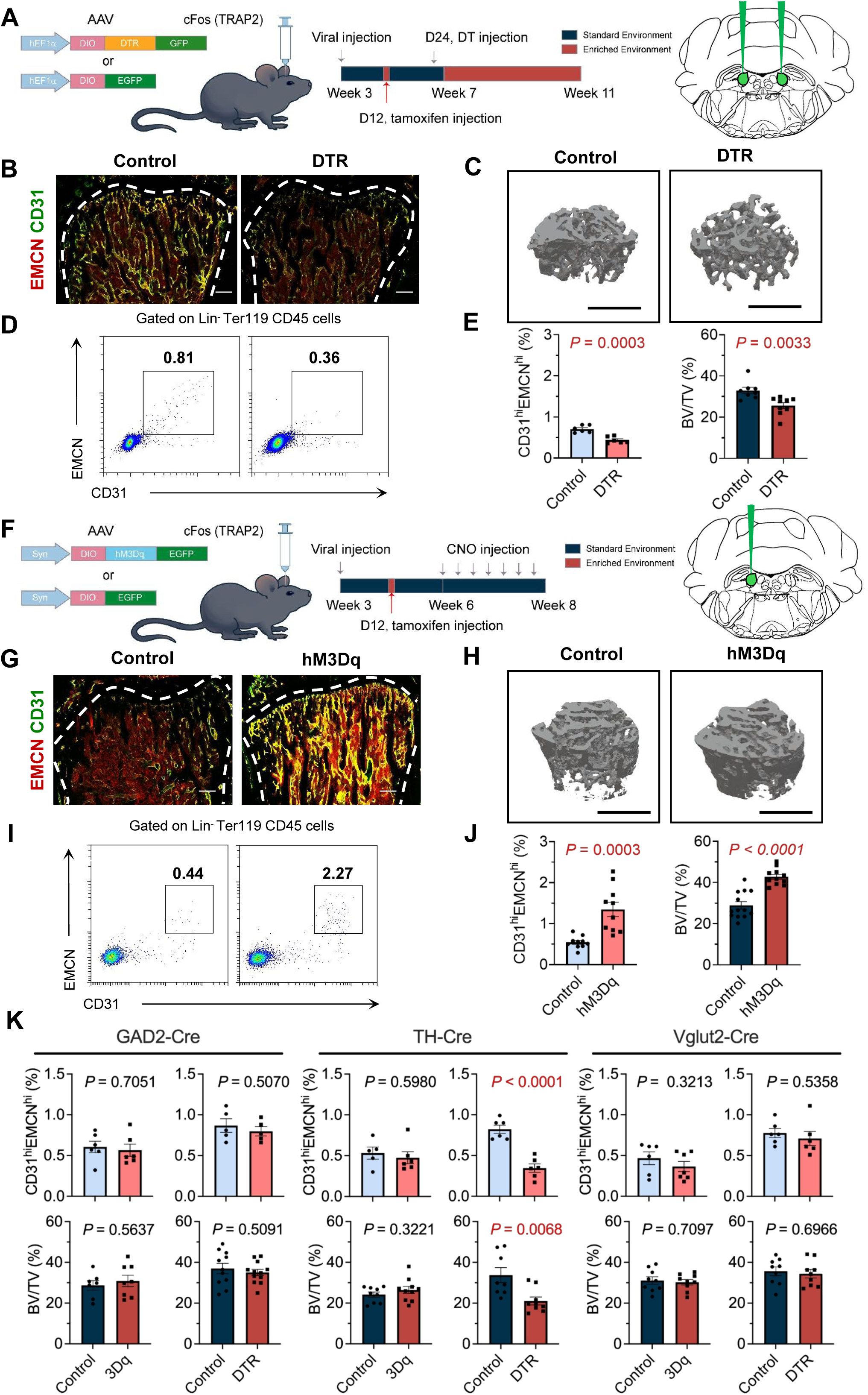
Functional manipulation and cell-type characterization of EE-labeled peri-LC neurons in the regulation of skeletal endothelial heterogeneity and bone density. (A) Schematic representation of DTR-mediated ablation of EE-labeled peri-LC neurons in cFos (TRAP2) mice. Virus was injected at 3 weeks of age, followed by tamoxifen administration on day 12 and a single diphtheria toxin injection on day 24 after viral injection. (B) Representative images of CD31 (green) and EMCN (red) dual-immunostained femur sections from EE-housed male mice with or without DTR-mediated ablation. Growth plate and endosteum are marked. Scale bars, 200 μm. (C) Representative μCT images of trabecular bone in the distal femur, Scale bars, 1 mm. (D) Representative flow cytometry plots showing CD31^hi^EMCN^hi^ endothelial cells from the femurs and tibiae of EE-housed male mice with or without DTR-mediated ablation. (E) *left*, Flow cytometry quantification of CD31^hi^EMCN^hi^ endothelial cells, n = 6, and *right*, bone volume/total volume (BV/TV) of EE-housed male mice with or without DTR-mediated ablation. Additional micro-CT parameters, including Ct.Th, Tb.Th, Tb.N, and Tb.Sp, are provided in Table 3. n = 8-9. (F) Schematic representation of chemogenetic activation of the peri-LC in cFos (TRAP2) mice. CNO was administered every two days from week six to week eight, seven times in total. (G) Representative images of CD31 (green) and EMCN (red) dual-immunostained femur sections from standard-housed male mice with or without chemogenetic activation. Growth plate and endosteum are marked. Scale bars, 200 μm. (H) Representative μCT images of trabecular bone in the distal femur, Scale bars, 1 mm. (I) Representative flow cytometry plots showing CD31^hi^EMCN^hi^ endothelial cells from femurs and tibiae of standard-housed male mice with or without chemogenetic activation. (J) *left*, Flow cytometry quantification of CD31^hi^EMCN^hi^ endothelial cells, n = 10, and *right*, bone volume/total volume (BV/TV) of standard-housed male mice with or without chemogenetic activation. Additional micro-CT parameters, including Ct.Th, Tb.Th, Tb.N, and Tb.Sp, are provided in Table 4. n = 11-13. (K) Flow cytometric quantification of CD31^hi^EMCN^hi^ endothelial cells and micro-CT analysis of BV/TV after DTR-mediated ablation or 3Dq-mediated activation of peri-LC neurons in Gad2-Cre, TH-Cre, and Vglut2-Cre mice. Additional micro-CT parameters, including Ct.Th, Tb.Th, Tb.N, and Tb.Sp, are provided in Table 9-14. Sample sizes for panel K: Gad2-Cre, 3Dq: n = 6 for flow cytometry and n = 7–8 for BV/TV; Gad2-Cre, DTR: n = 5 for flow cytometry and n = 11–12 for BV/TV; TH-Cre, 3Dq: n = 5–6 for flow cytometry and n = 9–10 for BV/TV; TH-Cre, DTR: n = 6 for flow cytometry and n = 8–9 for BV/TV; Vglut2-Cre, 3Dq: n = 6–7 for flow cytometry and n = 9 for BV/TV; Vglut2-Cre, DTR: n = 6 for flow cytometry and n = 9 for BV/TV.

To determine whether peri-LC activation altered skeletal endothelium through increased physical activity, we measured locomotor distance after activating EE-labeled peri-LC neurons. Rather than increasing locomotion, activation of these neurons reduced locomotor distance (Figure S6B), arguing against physical activity as the sole driver of endothelial changes. In addition, ablation of EE-labeled peri-LC neurons did not impair locomotor activity (Figure S6B), indicating that the skeletal phenotype was not secondary to altered motor capacity. To further assess whether locomotor activity activates peri-LC neurons, we introduced a running wheel under standard housing conditions and examined active neurons using cFos (TRAP2) × Ai14 mice. Running wheel exposure did not increase peri-LC neuronal activity compared with standard housing (Figure S6D and E). Consistently, calcium photometry showed reduced peri-LC neuronal activity during wheel running compared with exploratory behaviors (Figure 1K, *upper*, and Video 2). These results indicate that locomotor activity does not increase peri-LC neuronal activity under the experimental conditions used in this study. Additionally, co-labeling with PRV-EGFP and cFosTRAP2 × Ai14 identified activation in M1 and M2 brain regions (Figure S4A). However, activating M1 and M2 neurons in mice exposed to standard environments did not alter CD31^hi^EMCN^hi^ endothelium levels or bone density (Figure S7A-D). These findings suggest that the observed changes in skeletal endothelial heterogeneity and bone density are not mediated by physical activity or motor cortex activation but are instead regulated through the peri-LC. Collectively, these results highlight the peri-LC as a critical neural hub through which enriched environments exert systemic effects on skeletal CD31^hi^EMCN^hi^ endothelial cell heterogeneity and bone density. This brain-body regulatory pathway underscores the pivotal role of the peri-LC in translating environmental stimuli into physiological adaptations.

Previous studies have reported PRV labeling of LC neurons following injections into visceral organs such as the kidney and heart ^27,28^, indicating that LC neurons are involved in visceral autonomic regulation. Consistent with this notion, chemogenetic manipulation of peri-LC neurons modulated cardiovascular parameters, decreasing heart rate and diastolic blood pressure without significantly altering systolic blood pressure (Figure S8B and D). Acute dynamic exercise typically increases heart rate and systolic blood pressure ^29^, whereas diastolic blood pressure remains unchanged or decreases slightly ^30^. Therefore, the reduced heart rate and diastolic blood pressure, together with unchanged systolic blood pressure, observed after peri-LC activation do not resemble an exercise-like cardiovascular response. However, peri-LC manipulation did not alter renal calcium or phosphate homeostasis (Figure S9A-D), suggesting that the skeletal phenotypes observed in our study are unlikely to arise from indirect systemic changes in mineral metabolism.

### An EE-defined peri-LC neuronal ensemble is required for the regulation of CD31^hi^EMCN^hi^ endothelium

Under environmentally enriched conditions, the number of active TH- and GABA-expressing neurons was significantly increased, whereas VGLUT-expressing neurons showed no notable change (Figure S4E and F). Retrograde tracing experiments in TH-Cre × Ai14, Vglut2-Cre × Ai14, and Gad2-Cre × Ai14 mice further showed that neurons projecting from the distal femur to the peri-LC region were predominantly TH-expressing and GABAergic neurons (Figure S4C and D). Given that the peri-LC is anatomically and functionally distinct from the LC proper^31,32^, we next sought to determine whether these LC TH- and peri-LC GABA-expressing neuronal populations contribute to the regulation of CD31^hi^EMCN^hi^ endothelium and bone density.

TH-Cre, GAD2-Cre, and Vglut2-Cre mouse lines to examine how manipulating each subtype influences the phenotypes observed in our system (Figure S7E). Manipulation of neuronal subtypes within the peri-LC revealed a more complex regulatory pattern than expected. Selective activation or ablation of peri-LC GABAergic or glutamatergic neurons did not alter bone vascular endothelial heterogeneity or bone mass (Figure 3K, *left* and *righ*t). In contrast, ablation of LC-NE neurons significantly reduced the proportion of CD31^hi^EMCN^hi^ endothelial cells and led to a marked decrease in bone mass, whereas activation of LC-NE neurons did not produce the opposite phenotype (Figure 3K, *middle*). These findings indicate that the coordinated activity of the EE-defined LC/peri-LC neuronal ensemble, rather than isolated activation of a single neuronal subtype, is required for the regulation of skeletal endothelium and bone mass.

We also measured locomotor activity, heart rate, and blood pressure after activation or ablation of peri-LC neuronal subtypes in Gad2-Cre, TH-Cre, and Vglut2-Cre mice. In contrast to manipulation of EE-labeled peri-LC neurons in cFos (TRAP2) mice, subtype-specific manipulations did not affect locomotor activity (Figure S6A). Only Gad2-Cre-mediated ablation decreased diastolic blood pressure, and only Vglut2-Cre-mediated activation decreased heart rate, whereas other subtype-specific manipulations did not significantly alter cardiovascular parameters (Figure S8A and D). These findings suggest that EE-associated peri-LC activation engages multiple neuronal subtypes to coordinate peripheral physiological responses.

### Peri-LC regulates NE release in femurs

To explore how signals from the peri-LC are transmitted to the femurs, fluorescent staining in wild-type and transgenic mice was conducted. Results revealed dense sympathetic nerve terminals encircling CD31^hi^EMCN^hi^ endothelium (Figures S10A and B). The density of sympathetic nerve terminals was significantly greater around CD31^hi^EMCN^hi^ endothelium than around CD31^lo^EMCN^lo^ endothelium (Figure S9C and D). To better visualize the distribution of sympathetic nerves within long bones, 3D imaging of the femur was performed (Video 4). Results showed that sympathetic nerves entered the bone marrow through the nutrient foramen at the proximal femur and branched in three directions: upward, horizontally, and downward (Figure 4A and Video 4). EMCN expression was concentrated near the femoral head, endosteum, and growth plate (Figure 4B and Video 4). No significant sympathetic nerve terminals were detected near CD31^hi^EMCN^hi^ regions adjacent to the growth plate at 6.3× magnification (Figure 4A and B, yellow arrow). However, higher magnification (25×) revealed that sympathetic nerve extended toward endothelial cells next to growth plate, forming swollen, foot-like structures at their tips (Figure 4D cyan arrow, E, and Video 4, 5). Modeling analysis of sympathetic nerve in the diaphysis and epiphysis of the bone (Figure 4F) indicated that lateral branches in the diaphysis terminated near the endosteum (Figure 4G), while downward branches in the epiphysis extended progressively toward the growth plate (Figure 4H-J). We performed quantitative 3D proximity analysis by measuring the nearest-neighbor distance from the CD31^hi^EMCN^hi^ endothelial surface to the closest TH^+^ sympathetic fiber (Figure S10F). The distance between TH^+^ sympathetic fibers and the CD31^hi^EMCN^hi^ endothelial surface ranged from 0.0067 μm to 18.27 μm, with a median distance of 2.2 μm. This close spatial association indicates that sympathetic fibers are closely apposed to the vascular endothelial surface. These observations suggest that the peri-LC regulates bone vascular endothelial heterogeneity via the peripheral sympathetic nervous system.

**Figure 4:**
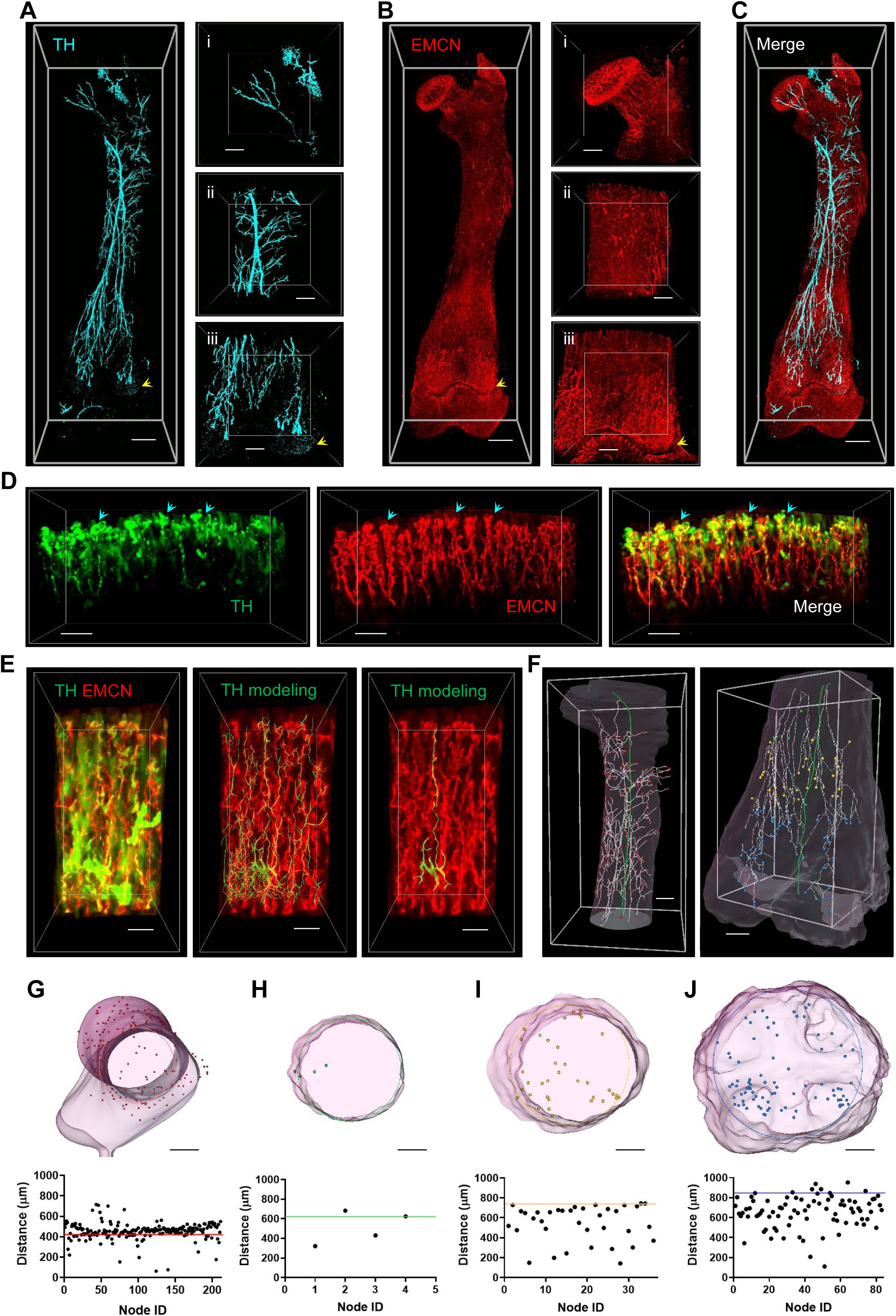
Sympathetic innervation and heterogeneous distribution of sympathetic nerve terminals in femurs of juvenile mice. (A) Three-dimensional bone clearing images of TH-immunostained femur of 3-week-old mice. Scale bars, left 600 μm, right 300 μm. (B) Three-dimensional bone clearing images of EMCN-immunostained femur 3-week-old mice. Scale bars, left 600 μm, right 300 μm. (C) Three-dimensional bone clearing images of EMCN and TH dual-immunostained femur 3-week-old mice. Scale bars, 600 μm. (D) Three-dimensional bone clearing image of EMCN (red) and TH (green) dual-immunostained femur near grow plate, cyan arrows indicate foot-like nerve endings. Scale bars, 100 μm. (E) *Left*, three-dimensional bone clearing image of EMCN (red) and TH (green) dual-immunostained femur near grow plate. *Middle* and *right*, sympathetic fiber modeling based on TH images (green). Scale bars, 50 μm. (F) Sympathetic nerve terminal (white line) and ending (ball) modeling based on TH image (Figure 4A). The green line represents the model-fitted bone central axis, which is used to calculate the shortest distance from the ending to the bone central axis. Left, diaphysis. Right, epiphysis. Scale bars, 300 μm. (G)-(J) *Upper*, Modeling of endosteum, growth plate, and sympathetic nerve endings in segmentations. Different colors correspond to locations of endings shown in Figure 4F. A cylinder was fitted based on the endosteum position, and its radius was used to draw circles in the corresponding figure. Scale bars, 300 μm. (G)-(J) *Lower*, Quantification of shortest distance from sympathetic nerve endings to central axis of the bone (green midline in Figure 4F). Positions of colored horizontal lines in the lower panel correspond to radii of circles in the upper panel.

Dual-NECa mice ^33^ crossed with Cdh5-Cre mice were used to express the GRAB_NE2m NE sensor specifically in vascular endothelial cells. Simultaneously, viruses carrying optogenetic or chemogenetic genes were injected into the peri-LC (Figure 5A), enabling real-time observations of NE release within the femur. To investigate the role of the peri-LC in NE release and endothelial heterogeneity near the femoral growth plate, blood vessels were imaged under optogenetic and chemogenetic stimulation. This endothelial-focused approach directly assessed peri-LC-evoked NE signaling at the vascular interface, without excluding additional NE actions on other bone-resident cells. Prior to optogenetic stimulation, green fluorescence was detected in the vascular endothelium (Figure 5B). Upon blue light stimulation of the peri-LC, green fluorescence rapidly illuminated and accumulated at distal endothelial cells closest to the growth plate (Figure 5B, red arrows). In contrast, fluorescence intensity remained largely unchanged at vascular bifurcations further from the growth plate (Figure 5B, blue arrows). Quantification of fluorescence at sprouting endothelial cells and bifurcation points confirmed these observations (Figure 5C). The entire process, from baseline to 10 minutes post-stimulation, was recorded (Video 7).

**Figure 5:**
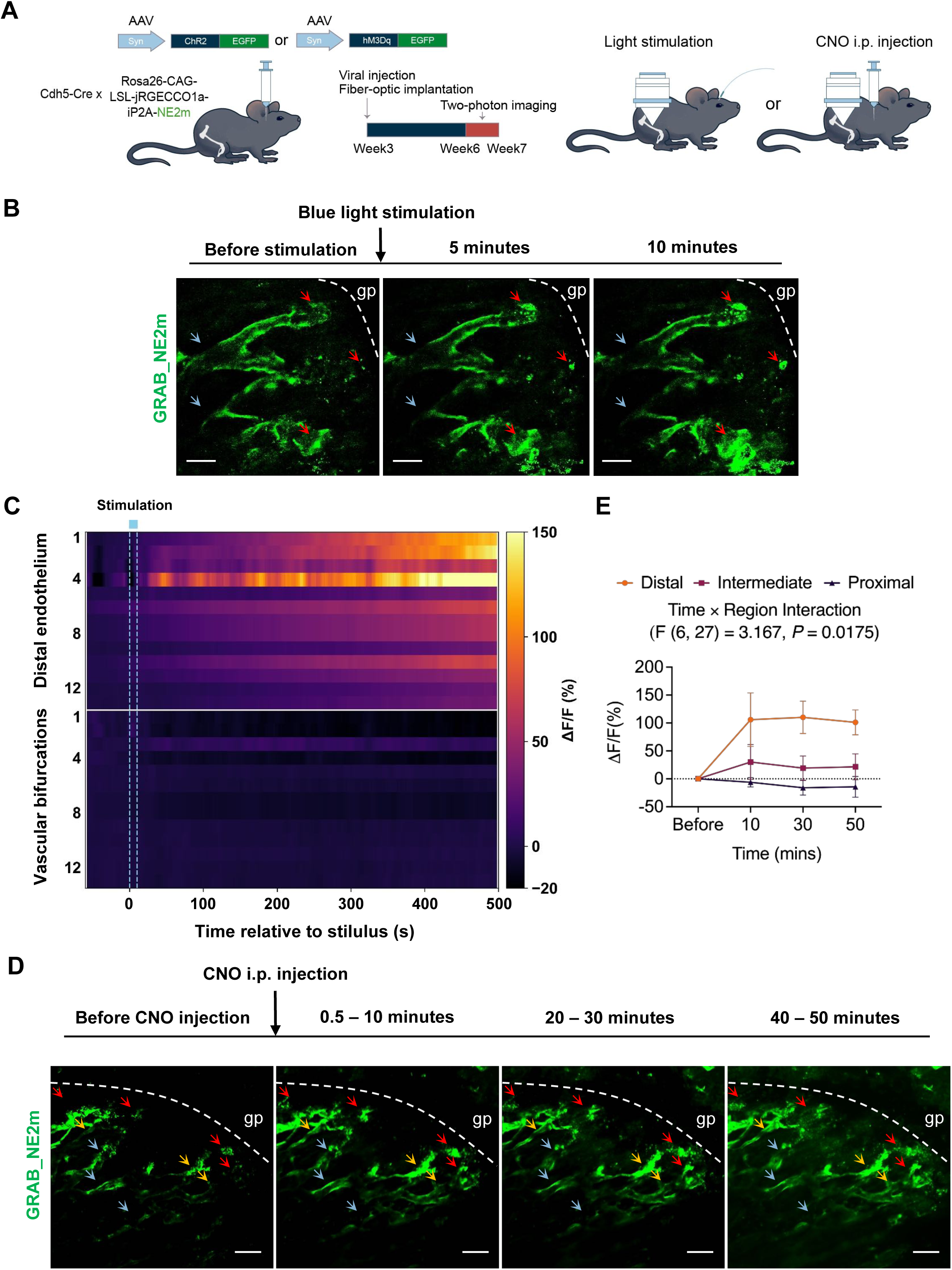
Real-time regulation of intraosseous NE release by the peri-LC. (A) Schematic of *in vivo* real-time two-photon imaging. Double-transgenic Rosa26-CAG-LSL-jTGECCO1a-iP2A-NE2m × Chd5-Cre mice were injected with AAV-Syn-ChR2 or AAV-Syn-hM3Dq virus targeting the peri-LC. NE sensors were specifically expressed in endothelial cells. Three weeks post-virus expression and recovery, anesthetized mice were positioned for two-photon imaging of CD31^hi^EMCN^hi^ endothelium near femoral growth plate. (B) Representative *in vivo* images of vascular sprouting endothelium (CD31^hi^EMCN^hi^) with optogenetic activation of the peri-LC. Growth plate is marked. Scale bars, 50 μm. (C) Quantification of simultaneously collected two-photon images of endothelial cells near growth plate with optogenetic activation of the peri-LC, n = 3. (D) Representative *in vivo* images (maximum intensity projection) of vascular sprouting endothelium (CD31^hi^EMCN^hi^) with chemogenetic activation of the peri-LC. Growth plate is marked. Scale bars, 100 μm. (E) Quantification of simultaneously collected two-photon images of endothelial cells near growth plate with chemogenetic activation of the peri-LC. Fluorescence changes in distal, intermediate, and proximal endothelial regions before and after CNO stimulation. n = 4. Data were analyzed using Two-way ANOVA with the Geisser-Greenhouse correction followed by Dunnett’s multiple comparisons test for comparisons with the pre-CNO baseline within each vascular region and Tukey’s multiple comparisons test for comparisons among vascular regions at each time point. The Two-way ANOVA model revealed significant effects of time (F(2.163, 83.64) = 9.289, P = 0.0002), vascular region (F(2, 39) = 13.15, P < 0.0001), and time × region interaction (F(6, 116) = 7.187, P < 0.0001). n = 4.

Chemogenetic stimulation of the peri-LC, achieved by intraperitoneal injection of clozapine-N-oxide (CNO), elicited changes in NE levels within the femur comparable to those observed during optogenetic stimulation (Figure 5D and E). Statistical analysis revealed significant effects of time (F (1.759, 15.83) = 4.005, *P* = 0.0435), vascular region (F (2, 9) = 6.867, *P* = 0.0155), and time × region interaction (F (6, 27) = 3.167, *P* = 0.0175), indicating region-specific fluorescence responses following CNO stimulation. Post hoc Dunnett’s multiple comparisons test showed that fluorescence in distal endothelial regions was significantly increased at 50 min after CNO stimulation compared with the pre-CNO baseline (Figure 5D, red arrows, and E), whereas intermediate and proximal regions showed no significant changes (Figure 5D, gold and blue arrows, and E). Consistently, Dunnett’s multiple comparisons test showed that distal endothelial regions exhibited higher fluorescence than proximal regions at 30 and 50 min after CNO stimulation. These results indicate that peri-LC activation preferentially induces fluorescence responses in distal endothelial regions near the growth plate.

Both optogenetic and chemogenetic stimulations induced fluorescence changes that persisted for more than 10 minutes (Figure 5C and E). During optogenetic stimulation, fluorescence transitioned from a point to a circular area (Figure 5B, bottom right red arrow), likely reflecting NE neurotransmitter diffusion. These results indicate that peri-LC activation preferentially induces NE release around distal endothelial cells near the growth plate, where foot-like sympathetic fiber structures were observed at the light-microscopy level and corresponded to regions of high sympathetic nerve density (Figure 4D, Figure S9C and Figure 5C). Due to technical limitations, these experiments were performed in anesthetized mice and were therefore designed to assess relative changes and spatial patterns of norepinephrine (NE) signaling under controlled conditions, rather than absolute sympathetic tone. This spatial heterogeneity in sympathetic innervation may contribute to the heterogeneous distribution of vascular endothelial subtypes in bone. These experiments demonstrated that the peri-LC directly regulates NE dynamics in the skeletal system, the peri-LC appears to regulate this endothelial heterogeneity via the peripheral sympathetic system.

Due to limitations of the transgenic mouse models, the entire peri-LC brain region was activated to observe its control over NE release within the femur. To verify whether whole-region activation yields similar effects to specific activation of EE-labeled neurons in long bones, similar procedures were performed on wild-type C57 mice (Figures S10A). Consistent with specific activation, activation of the whole peri-LC, even in mice housed under standard environments, led to an increase in CD31^hi^EMCN^hi^ endothelium and bone density (Figure S10B-E).

### Sympathetic nervous system regulates endothelial cell heterogeneity

Next, we utilized retroAAV to regulate the local sympathetic nervous system in the femur and validate our findings. RetroAAVs carrying chemogenetic genes were injected into the bone marrow cavity near the femoral growth plate of TH-Cre mice, enabling inhibition of local sympathetic nerve activity (Figure 6A and F). Inhibition of sympathetic nerve activity resulted in a marked reduction in CD31^hi^EMCN^hi^ endothelium and bone density (Figures 6B-E).

**Figure 6:**
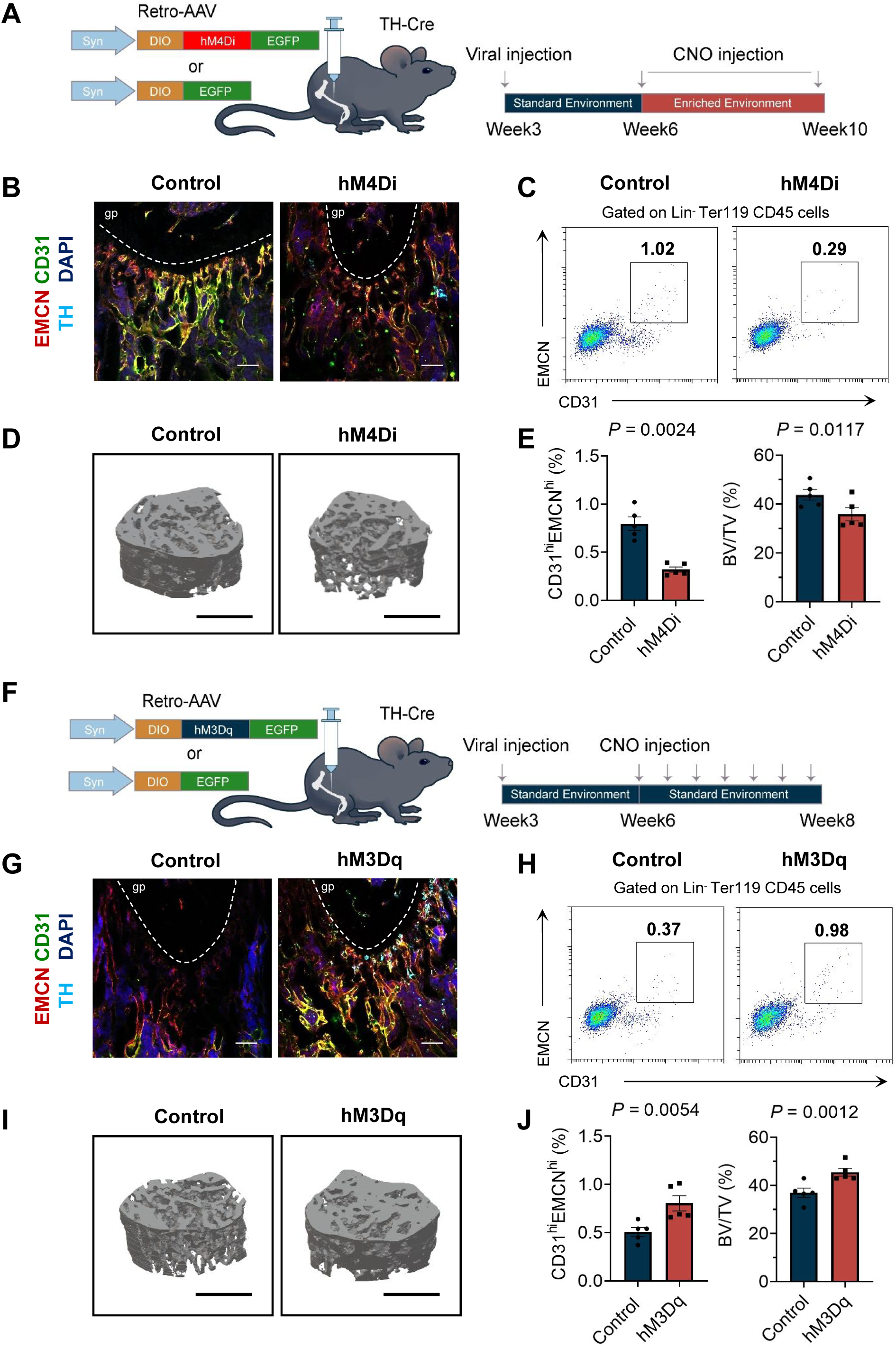
Sympathetic regulation of endothelial cell heterogeneity in femurs. (A) Schematic representation of chemogenetic inactivation of local sympathetic nerves in the femurs of EE-housed TH-Cre mice. CNO was administered every two days from week six to week ten. (B) Representative images of immunostained-femur sections from mice with chemogenetic inactivation of local sympathetic nerves. Growth plate is marked. Scale bars, 100 μm. (C) Representative flow cytometry plots showing CD31^hi^EMCN^hi^ endothelial cells from the femurs of EE-housed male mice with or without chemogenetic inactivation of local sympathetic nerves. (D) Representative μCT images of trabecular bone in the distal femur, scale bars, 1 mm. (E) *left*, Flow cytometry quantification of CD31^hi^EMCN^hi^ endothelial cells, and *right*, bone volume/total volume (BV/TV) of EE-housed male mice with or without chemogenetic inactivation. Additional micro-CT parameters, including Ct.Th, Tb.Th, Tb.N, and Tb.Sp, are provided in Table 15. n = 5. (F) Schematic representation of chemogenetic activation of local sympathetic nerves in the femurs of standard-housed TH-Cre mice. CNO was administered every two days from week six to week eight. (G) Representative images of immunostained-femur sections from mice with chemogenetic activation of local sympathetic nerves. Growth plate is marked. Scale bars, 100 μm. (H) Representative flow cytometry plots showing CD31^hi^EMCN^hi^ endothelial cells from the femurs of standard-housed male mice with or without chemogenetic activation of local sympathetic nerves. (I) Representative μCT images of trabecular bone in the distal femur, scale bars, 1 mm.. (J) *left*, Flow cytometry quantification of CD31^hi^EMCN^hi^ endothelial cells, and *right*, bone volume/total volume (BV/TV) of standard-housed male mice with or without chemogenetic activation. Additional micro-CT parameters, including Ct.Th, Tb.Th, Tb.N, and Tb.Sp, are provided in Table 16. n = 5.

Specifically, we locally administered 6-hydroxydopamine (6-OHDA) to selectively ablate sympathetic nerve fibers within the bone marrow, thereby reducing local sympathetic activity (Figure S12A). Compared with the contralateral femur, the femur receiving 6-OHDA treatment exhibited a significant reduction in CD31^hi^EMCN^hi^ endothelial cells (Figure S12B), accompanied by a corresponding decrease in bone density (Figure S12B). Conversely, in mice housed under standard conditions, local activation of the sympathetic nerves increased the levels of CD31^hi^EMCN^hi^ endothelium and bone density (Figures 6G-J). These observations demonstrate that local sympathetic nerve activity in the femur can directly modulate the heterogeneity of bone vascular endothelial cells and associated bone density.

Previous studies examining the effects of beta-adrenergic receptor blockers, such as propranolol, on bone density have produced conflicting outcomes, with reports of both increases and decreases ^34–37^. To clarify this relationship, different doses of propranolol were administered to 4-week-old mice over a 4-week period via oral gavage. Analysis revealed a dose-dependent decrease in CD31^hi^EMCN^hi^ endothelial cell levels and bone density (Figures S12B-D). Notably, pharmacological inhibition of β2-adrenergic receptors with ICI-118551 also decreased CD31^hi^EMCN^hi^ endothelial cells and bone density. These findings suggest that the sympathetic nervous system regulates CD31^hi^EMCN^hi^ endothelial cells and bone density through β-adrenergic receptor signaling, with β2-adrenergic receptors likely playing a major role. Consistent with this possibility, we confirmed robust expression of β2-adrenergic receptors in bone vascular endothelial cells (Figure S13A). They further suggest that β-adrenergic blockade may contribute to bone loss.

The sympathetic nervous system has been shown to influence angiogenesis through β2-adrenergic receptors on endothelial cells^16^. To investigate the role of these receptors in regulating endothelial heterogeneity, we generated endothelial-specific β2-adrenergic receptor knockout mice. One week after tamoxifen-induced endothelial *adrb2* knockout, β2-adrenergic receptor expression in endothelial cells near the growth plate was significantly reduced (Figure S13B and C). This reduction was accompanied by decreased CD31^hi^EMCN^hi^ endothelium at the same location (Figure 7A and B, *upper*). Tamoxifen was administered daily from postnatal day 10 to 14, and samples were collected at postnatal day 21. This reduction impaired angiogenic sprouting, evidenced by diminished filopodia formation (Figures S13D and E). By the eighth week, knockout mice (knockout efficiency = 54.7%, Figure S13G-I) exhibited pronounced decreases in both CD31^hi^EMCN^hi^ endothelium and bone density compared with littermate controls (Figures 7B, *middle* and *lower*, 7C–E, and S13F). Consistently, immunofluorescence analysis showed that expression of the angiocrine factor Noggin was also significantly reduced in *adrb2* knockout mice compared with control mice (Figure S2C). Even after 4 weeks of environmental enrichment, knockout mice displayed significantly lower levels of CD31^hi^EMCN^hi^ endothelium and bone density compared to littermate controls (Figures 7F-H). Furthermore, unlike the effects achieved in wild type mice, chemogenetic activation of the peri-LC in knockout mice failed to increase CD31^hi^EMCN^hi^ endothelium levels or bone density (Figure S14A-E). These findings underscore the critical role of β2-adrenergic receptors in mediating angiogenesis, regulating endothelial heterogeneity, and maintaining bone density.

**Figure 7:**
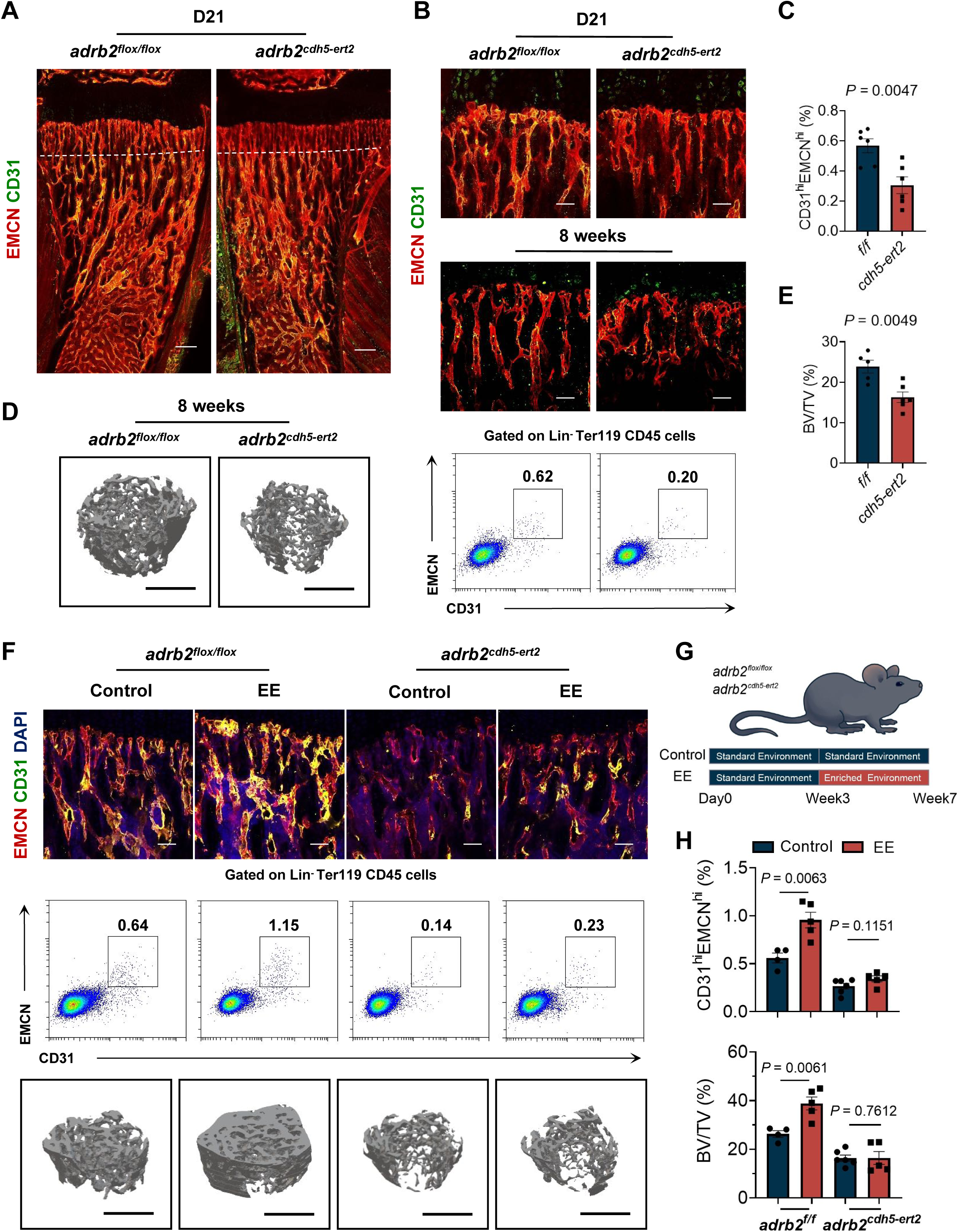
Beta2-adrenergic receptor-mediated regulation of endothelial heterogeneity by the sympathetic nervous system. (A) and (B) Upper, Representative images of D21 *adrb2*^flox/flox^ and *adrb2*^cdh5-ert2^ mouse femurs stained with CD31 (green) and EMCN (red). Scale bars, A, 200 μm, B, 50 μm (B) *Middle*, Representative images of 8-week-old *adrb2*^flox/flox^ and *adrb2*^cdh5-ert2^ mouse femurs stained with CD31 (green) and EMCN (red). Scale bars, 50 μm (B) *Lower*, Representative flow cytometry plots showing CD31^hi^EMCN^hi^ endothelial cells from femurs and tibiae of 8-week-old *adrb2*^flox/flox^ and *adrb2*^cdh5-ert2^ mice. (C) Flow cytometry quantification of CD31^hi^EMCN^hi^ endothelial cells in 8-week-old *adrb2*^flox/flox^ and *adrb2*^cdh5-ert2^ mice. n = 6. (D) Representative μCT images of trabecular bone in the distal femur, scale bars, 1 mm. (E) Bone volume/total volume (BV/TV) of 8-week-old *adrb2*^flox/flox^ and *adrb2*^cdh5-ert2^ mice. Additional micro-CT parameters, including Ct.Th, Tb.Th, Tb.N, and Tb.Sp, are provided in Table 18. n = 5-6. (F) *Upper*, Representative images of immunostained-femur sections from *adrb2*^flox/flox^ and *adrb2*^cdh5-ert2^ mice housed under standard or enriched environments. Scale bars, 50 μm. *Middle*, Representative flow cytometry plots showing CD31^hi^EMCN^hi^ endothelial cells from femurs and tibiae. *Lower*, Representative μCT images of trabecular bone in the distal femur, scale bars, 1 mm. (G) Schematic representation of standard- or EE-housed *adrb2*^flox/flox^ and *adrb2*^cdh5-ert2^ mice. (H) *Top*, Flow cytometry quantification of CD31^hi^EMCN^hi^ endothelial cells (n = 4–6), and bottom, bone volume/total volume (BV/TV) (n = 4–6) of standard- or EE-housed *adrb2*^flox/flox^ and *adrb2*^cdh5-ert2^ mice.

Previous studies have reported that β2-adrenergic receptor knockout in osteoblasts promotes bone formation in 6-month-old mice ^38^. However, when the levels of CD31^hi^EMCN^hi^ endothelium and bone density were assessed in 4-week-old osteoblast-specific β2-adrenergic receptor knockout mice, no significant changes were observed (Figures S15A-C). To ensure consistency with the osteoblast-specific knockout strategy, we used Cdh5-Cre, rather than Cdh5-CreER, for endothelial-specific β2-adrenergic receptor deletion in this comparison. At the same age, Cdh5-Cre-mediated endothelial-specific β2-adrenergic receptor knockout mice showed substantial reductions in CD31^hi^EMCN^hi^ endothelium and bone density (Figure S15D-F). These results indicate that enriched environment regulates CD31^hi^EMCN^hi^ endothelium levels through the peripheral sympathetic nervous system and the β2-adrenergic receptors on vascular endothelial cells. Furthermore, we performed local, cell type-specific deletion of β2-adrenergic receptors within the same animals, with endothelial deletion in one femur and osteoblastic deletion in the contralateral femur. This paired design allowed direct comparison of endothelial and osteoblastic β2-adrenergic receptor signaling in the regulation of endothelium and bone phenotypes. Viral expression and targeting were robust (Figure S15G), and paired analyses within the same animals revealed that bone density was significantly reduced on the endothelial *adrb2*–deleted side compared with the osteoblastic control femur (FigureS15H). These results indicate that endothelial β2-adrenergic signaling contributes locally to the maintenance of bone mass.

The rostral ventrolateral medulla (RVLM) is a classical brain region involved in both basal and reflex regulation of sympathetic outflow ^39^. We used the RVLM as a comparison region to determine whether brain regions broadly involved in the regulation of peripheral sympathetic output can modulate skeletal endothelial heterogeneity and bone density, or whether the peri-LC exerts a distinct regulatory effect. Compared to peri-LC manipulations, chemogenetic activation of TRAP-labeled RVLM neurons did not alter the abundance of CD31^hi^EMCN^hi^ endothelial cells (Figure S15I, *left*). However, did significantly reduced bone mass (Figure S15I, *left*). Notably, ablation of RVLM TRAP-labeled neurons did not produce detectable changes in either endothelial heterogeneity or bone mass (Figure S15I, *right*). Although the RVLM is a well-established autonomic center controlling peripheral sympathetic tone, our data suggests that it does not regulate bone vascular endothelial heterogeneity. The RVLM is a classical brain region involved in cardiovascular regulation. Consistent with this role, chemogenetic activation of the RVLM in C57BL/6J mice increased heart rate and blood pressure (Figure S8C and D). However, activation or ablation of EE-labeled RVLM neurons did not alter heart rate or blood pressure (Figure S8B and D). This response pattern differed from that observed after peri-LC activation, which decreased heart rate and diastolic blood pressure (Figure S8B and D). In addition, unlike peri-LC activation, RVLM activation did not affect locomotor activity (Figure S6B and C). Taken together, these results suggest that the central regulation of sympathetic output and its effects on peripheral organs are not uniform across brain regions. In contrast to other sympathetic regulatory nuclei, e.g., RVLM, the peri-LC exerts a unique influence on bone vascular endothelial heterogeneity and bone mass.

Chronic restraint stress (CRS) was found to alter emotional and behavioral responses in mice (Figure S16B and C) and to reduce bone mass (Figure S16D and E). To examine the potential restorative effects of environmental enrichment, stress-modeled mice were subsequently housed in either standard or enriched environments (Figure S16A). Mice in standard environments demonstrated a recovery in bone density (Figure S16G and H *right*, CRS-Sham group) to levels comparable to those of non-modeled mice (Figure S16G and H *right*, Control group), as well as similar CD31^hi^EMCN^hi^ endothelium levels (Figure S16F and H *left*). However, the EE-housed mice (CRS-EE group) showed further increases in both CD31^hi^EMCN^hi^ endothelium levels and bone density compared to standard-housed mice (Figure S16F-H). Enriched environments reversed stress-induced bone loss.

## Discussion

The ability of organisms to perceive and adapt to their external environment is fundamental for maintaining physiological homeostasis. In the present study, enriched environments elicited a widespread increase in neuronal activity in mice, particularly within the pons and medulla. These brainstem regions play essential roles in the relay of signals between different brain areas and between the brain and body, underpinning essential physiological processes and systemic homeostasis ^40^. The observed enhancement in neuronal activity within these regions suggests that environmental enrichment may exert broad physiological effects through central neural pathways. The increased peri-LC neuronal activity may be driven, at least in part, by the greater spatial complexity of the enriched environment. Indeed, our results demonstrated that environmental enrichment activates the LC and peri-LC brain regions, with neural signals originating from these regions subsequently transmitted to the skeletal system via the peripheral sympathetic nervous system, where they modulated vascular endothelial heterogeneity. This regulation is mediated by β2-adrenergic receptor signaling, which helps maintain CD31^hi^EMCN^hi^ endothelial characteristics in regions associated with sympathetic innervation, promotes the release of angiocrine factors such as Noggin, and ultimately contributes to increased bone density. These findings highlight the dual role of the LC and peri-LC regions in regulating both brain states and long-term physiological processes, revealing a novel brain-body interaction mechanism and the intricate regulation of peripheral innervation.

### Peri-LC and LC influence physiological states

The LC-NE system is a key modulator within the brain, influencing sensory processing, arousal, attention, mood, memory, and cognition across various behavioral contexts ^22^. The peri-LC and LC receive extensive projections from multiple brain regions, establishing their importance as integrative hubs for neural regulation ^23,41^. LC activity is widely described as following an inverted-U functional pattern, in which low LC activity is associated with quiescent or sleep-like states, whereas excessive LC activation is linked to stress-related conditions such as anxiety and PTSD ^22^. Only when LC output remains within an intermediate range does it support optimal physiological function ^22^. GABAergic neurons within the peri-LC have been shown to modulate peri-LC-NE neuronal activity, effectively regulating arousal levels^23,32^. In the current study, environmental enrichment was shown to significantly enhance the activity of both TH-expressing and GABAergic neurons in the LC/peri-LC region. However, activation of any single neuronal subtype alone did not reproduce the physiological and skeletal effects observed under enriched environment conditions. Based on these findings, we propose that peri-LC GABAergic neurons may help maintain LC-NE neuronal activity within an optimal physiological range, thereby supporting a peripheral bone-vascular state that favors CD31^hi^EMCN^hi^ endothelial cells and bone accrual. The circuit mechanisms by which peri-LC GABAergic neurons fine-tune LC-NE neuronal activity to maintain this optimal range remain an important direction for future investigation. Disrupting this balance—either by driving LC activity beyond its optimal range or by eliminating TH neurons—may impair the regulatory effect on bone vasculature and abolish the associated bone-mass benefits. This ensemble-level organization is essential for linking environmental enrichment to peripheral physiological regulation.

We propose that enriched environments, by increasing spatial complexity and imposing complex sensory, cognitive, and physiological demands, drive coordinated neuronal activity within the LC/peri-LC region. This heightened activity not only modulates arousal within the central nervous system but also influences peripheral physiological processes. CD31^hi^EMCN^hi^ endothelium represents a highly proliferative and metabolically active endothelial subtype that promotes bone formation by secreting angiocrine factors^18^. However, the abundance of these vessels and associated osteoprogenitors diminishes significantly in aged skeletal systems, contributing to reduced bone regeneration and metabolic capacity ^18^. The peri-LC plays a critical role in regulating the heterogeneity of skeletal vascular endothelium, preserving the unique characteristics of CD31^hi^EMCN^hi^ endothelium. This regulation enhances bone formation, contributing to improved skeletal health and broader physiological resilience.

The role of bone in maintaining body homeostasis extends beyond its structural and protective functions ^42^. The skeleton serves as a dynamic reservoir for essential minerals such as calcium and phosphorus, which are essential for the functioning of other physiological systems (e.g., nerve signaling transmission, muscle contraction, enzyme activation, and cell signaling). During periods of deficiency, bones release these minerals into the bloodstream to restore homeostatic balance ^43^. Maintaining high bone density provides a significant adaptive advantage by ensuring sufficient mineral reserves and robust skeletal integrity. Peri-LC activation in enriched environments enhances bone density through endothelial heterogeneity regulation, supporting systemic stability and improving adaptation to environmental challenges (e.g. starvation and temperature extremes). This highlights a novel pathway linking neural and vascular systems to long-term physiological adaptation.

Our results suggest that locomotor activity is unlikely to be the driver of EE-induced changes in skeletal endothelial heterogeneity and bone density. In our experimental setting, locomotor activity neither directly increased bone density nor accounted for the peri-LC neuronal activation associated with skeletal endothelial and bone changes. Although exercise is generally considered beneficial for bone mass, its skeletal effects depend on multiple factors, including exercise type, intensity, duration, stress level, metabolic status, age, and sex^44–46^. Thus, physical activity and bone accrual do not necessarily show a simple positive relationship. Consistent with this idea, a previous study showed that the same duration and form of physical activity produced distinct neural outcomes depending on environmental context: a walk in a natural environment decreased amygdala activity, whereas an equivalent walk in an urban environment did not^47^. Similarly, our findings suggest that EE-induced skeletal effects depend not on locomotor activity alone, but on the broader environmental context and its engagement of specific neural circuits.

### Neural-vascular EC coupling mediates brain-bone control

The intricate relationship between peripheral nerves and blood vessels is essential for their development, spatial organization, and functional coordination, particularly through the actions of endothelial cells ^8^. While the role of the sympathetic nervous system in promoting angiogenesis is well-established ^16^, the mechanisms by which the brain regulates peripheral vascular endothelial heterogeneity remain largely unknown. This study provides evidence that signals from the central nervous system are transmitted to the vasculature of long bones via the peripheral sympathetic nervous system. Notably, 3D imaging revealed that sympathetic fibers were concentrated near the endosteum and growth plate, where they ran immediately adjacent to vascular endothelial cells and formed enlarged, foot-like structures near distal sprouting endothelial cells. These structures corresponded to sites of real-time NE release in femurs following peri-LC stimulation. Upon peri-LC activation in the brain, neurotransmitters released by sympathetic nerves were primarily captured by distal endothelial cells near the growth plate, while vascular branches further from this region remained largely unaffected. Additionally, the spatial gradient of NE sensor fluorescence increased from the proximal to distal region, with the strongest response near the growth plate, suggesting that the NE signal was unlikely to be derived from systemic circulation or other organs. Together with the decreases in heart rate and diastolic blood pressure after peri-LC activation, as well as the absence of changes in circulating calcium and phosphate levels, these findings do not support a systemic increase in NE signaling after peri-LC activation, but instead suggest a preferential induction of local sympathetic NE release within bone. Indeed, the distribution of sympathetic nerve terminals closely aligned with the localization of CD31^hi^EMCN^hi^ endothelial cells within long bones, highlighting the spatial specificity of peri-LC-induced sympathetic activation. Moreover, endothelial heterogeneity was not significantly altered in other skeletal sites, such as vertebrae or skull, likely due to differences in their embryonic origins and developmental mechanisms ^48,49^. Although both the RVLM and peri-LC can influence peripheral sympathetic output, our findings suggest that they engage distinct central-to-peripheral regulatory programs. RVLM activation increased heart rate and blood pressure and reduced bone density, but did not alter skeletal endothelial heterogeneity or locomotor activity. In contrast, peri-LC activation decreased heart rate and diastolic blood pressure, regulated skeletal endothelial heterogeneity, increased bone density, and altered locomotor activity. These findings underscore the specificity of peri-LC regulation in modulating long bone density via peripheral neural-vascular coupling, achieved through distinct central brain regions, varied sympathetic connections, and differential nerve terminal distributions.

Previous studies have explored the neural regulation of bone metabolism ^35^, with research showing that osteoblast-specific knockout of β2-adrenergic receptors increases bone density ^38^, raising the possibility of beta blockers as a treatment for osteoporosis. However, clinical outcomes with beta blockers have been inconsistent, with reports of both beneficial and neutral effects ^34,36,37^. This study demonstrates that the sympathetic nervous system enhances bone density through β2-adrenergic receptors on endothelial cells. Our findings suggest that these inconsistencies may stem from the influence of the sympathetic nervous system on bone vascular endothelial heterogeneity.

Endothelial cells, which line the interior surface of blood vessels, establish an extensive network of capillary branches that form connections with almost all cell types across different organs ^17^. These cells exhibit organ-specific heterogeneity, characterized by unique gene expression profiles and functional markers ^10,50^. Our study focused on the skeletal system, leaving open the question of whether the peri-LC or other brain regions regulate vascular endothelial heterogeneity in peripheral organs such as the liver, lung, or spleen. To the best of our knowledge, this is the first study to demonstrate central nervous system control over peripheral vascular endothelial heterogeneity.

### Enriched environments reverse stress-induced bone loss

Previous research has demonstrated that activation of the peripheral sympathetic nervous system can exert diverse effects on the physiological and pathological states of organs ^16,51–56^. Our findings revealed that localized activation of sympathetic nerve terminals in long bones sustains CD31^hi^EMCN^hi^ vasculature, promoting increased bone density and enhancing systemic resilience. Targeting this neural-vascular interaction represents a promising therapeutic strategy for managing bone-related conditions, including osteoporosis and fractures. Our findings also indicated that chronic stress, which leads to bone density loss, can be effectively mitigated through exposure to enriched environments. Enriched environments were shown to reverse stress-induced bone loss, offering a natural and non-invasive approach for restoring bone health in stress-induced osteoporosis. By harnessing the intrinsic capacity of the body for adaptation, enriched environments could serve as a complementary therapy alongside conventional medical treatments, offering a holistic approach to bone health management.

### Limitation of the study

Our study has several limitations. We did not directly assess whether social interaction within the enriched environment affects peri-LC neuronal activity, skeletal endothelial heterogeneity, or bone density. In addition, the mechanisms by which the peri-LC connects with the peripheral sympathetic nervous system remain to be defined, including whether other brain nuclei participate in this regulatory pathway. Finally, although sympathetic nerves can directly regulate mesenchymal stromal cells ^57^, we did not examine whether direct sympathetic regulation of MSCs contributes to the bone-density phenotype observed in this study.

## Conclusions

This study provides compelling evidence that enriched environments exert positive effects not only on the brain but also on peripheral tissues and organs via neural mechanisms (Figure S17, summary graph). LC activity is traditionally associated with arousal states, with peri-LC GABAergic neurons playing coordinated roles in modulating LC activity. Mice exposed to enriched environments demonstrated the ability to maintain an optimal physiological state via LC and peri-LC activation. Our findings expand the regulatory role of the LC and peri-LC regions from central neural functions to peripheral physiological processes. Notably, the peri-LC was shown to regulate peripheral endothelial cell heterogeneity and bone health through unique and precise modulation of the sympathetic nervous system, underscoring the critical role of neural-vascular coupling in linking environmental stimuli to systemic physiological adaptations. By advancing our understanding of brain-body interactions, this study highlights the therapeutic potential of neural regulation and environmental enrichment for bone-related diseases.

## Supporting information

Video 1 Whole-Brain cFos

Video 2 Behavior-Aligned Calcium Signal Tracking During Running

Video 3 Behavior-Aligned Calcium Signal Tracking During Tent Entryexit And Grooming

Video 4 3D Femur Th Emcn

Video 5 25X Th Emcn

Video 6 25X Modeling

Video 7 Peri-LC NE ECs

## Supplemental information

**Supplemental Videos,** Video 1-7

Video 1_Whole-brain cFos related to Figure 1

Video 2_ Behavior-aligned calcium signal tracking during running

Video 3_ Behavior-aligned calcium signal tracking during tent entry/exit and grooming

Video 4_3D Femur TH EMCN

Video 5_25X TH EMCN and Video 6_25X modeling related to Video 5

Video 7_Peri-LC NE ECs related to Figure 5

## Acknowledgments

We thank Dr. Junlei Chang for providing Cdh5-ert2-Cre mouse line, Dr. Yingjie Zhu for providing cFos (TRAP2) mouse line. This work was supported by Science and Technology Innovation 2030 Major Project (STI2030-Major Projects 2022ZD0205700), Shenzhen Medical Research Fund (B2302011), Young Scientists Fund of the National Natural Science Foundation of China (31900829), Natural Science Foundation of Guangdong Province, China (2019A1515011122), and China Postdoctoral Science Foundation (2018M643244).

## Methods

### Experimental model and subject details

All experiments were conducted in accordance with the Guide for the Care and Use of Laboratory Animals (Eighth Edition, 2011) and were approved by the Institutional Animal Care and Use Committee (IACUC) at the Shenzhen Institute of Advanced Technology, Chinese Academy of Sciences. Experimental animals included mice under 10 weeks of age, including C57BL/6J and various transgenic strains, as specified below. Prior to any experimental procedures, mice were acclimated to handling by the experimenter for approximately 5 minutes per day for a minimum of 3 days to minimize stress and facilitate subsequent interventions.

Transgenic mouse lines, including adrb2 flox, Ai14, Chd5- Cre, Cdh5-ert2-Cre, cfosTRAP2, Gad2-Cre, OC-Cre, Rosa26-CAG-LSL-jRGECCO1a-iP2A-NE2m, TH-Cre, and Vglut2-Cre mice, were used to elucidate the mechanisms underlying the effects of enriched environments on endothelial cell heterogeneity and bone density through the peri-LC. To induce endothelial-specific *adrb2* knockout, tamoxifen (Sigma, USA) injections (500 μg per mouse) were performed daily from postnatal day D10 to D14 for five consecutive days.

Mice were housed under a standard 12-hour light/12-hour dark cycle with *ad libitum* access to food and water. The room temperature was maintained at 22 °C. To minimize the potential influence of sex-specific differences on study outcomes, only male mice were included in the bone density-related experimental cohorts.

Environmental enrichment was employed to examine its effects on mouse physiology and behavior, following a previously established framework ^21^. The enriched cages measured 55 cm × 39 cm × 27 cm (Figure S1A) and consisted of two levels: the upper level contained a small tent and a food bowl, while the lower level included a water bottle, running wheel, climbing pole, tunnel, ball, and soft tissue for nesting, with *ad libitum* access to food and water. These cages housed groups of 10–12 mice, providing greater sensory, cognitive, motor, and social stimulation compared to standard housing. In contrast, control mice were housed in standard cages in groups of 5–6, with limited space and *ad libitum* access to food and water. This comparative setup was designed to evaluate the impact of environmental complexity on biological and behavioral parameters, ensuring consistency and reproducibility by adhering to the established enrichment protocols ^21^.

## Method details

### TRAP2 labeling

Fos2A-iCreER (cfosTRAP2; JAX 030323) mice were backcrossed on a C57BL6/J background and bred with B6;129S6-Gt(ROSA)26Sortm14(CAG-tdTomato)/Hze/J (Ai14; JAX 007908) mice. Both male and female mice were included in the TRAP2 labeling experiments. At 6 weeks of age, mice were handled and acclimatized to fresh, clean cages (control group) or enriched environment (EE) cages (EE group) for 1 hour per day over 3 days prior to labeling. On the day of labeling, mice received an intraperitoneal injection of freshly prepared tamoxifen (Sigma, USA) at a dose of 100 mg/kg, dissolved in corn oil (MCE, USA). Mice in the EE group were housed in EE cages for 48 hours before being returned to their home cages. All mice were euthanized 12 days later to ensure full fluorophore expression.

### Whole-brain tissue clearing and analysis

#### Fixation

Mice were perfused with ice-cold phosphate-buffered saline (PBS) and 4% paraformaldehyde (PFA). After sampling, the brains were fixed in 4% PFA at 4 °C with gentle shaking overnight. After fixation, the brains were washed with PBS 2–3 times (2 hours each).

#### Clearing

Brains were processed using a Nohai Tissue Clearing Kit (Cat No: NH-CR-230701, Nohai Biotechnology, China). Fixed brains were placed in a centrifuge tube containing 20 mL of delipidation solution and gently shaken at 4 °C for 7–8 days, with the solution replaced every 2 days. After delipidation, the samples were transferred to PBS at room temperature and washed for 2 hours, repeated three times. The brains were then immersed in refractive index matching solution until fully cleared, after which optical 3D imaging was performed.

#### Three-dimensional (3D) imaging and image processing of mouse brain

3D fluorescence imaging of the brains was performed using a Nohai LS18 light-sheet microscope (Nohai Biotechnology, China). For whole-sample 3D image acquisition, each field of view was illuminated four times by the light sheet, and a 1×/0.25 NA air objective (Olympus MVPLAPO) was used^58^. Under the selected imaging conditions, the microscope magnification was set to 4×, with a spatial resolution of 3.3 × 3.3 × 7 µm^3^. Data were modeled and analyzed using Amira (Thermo Fisher Scientific, USA, v 2021.2).

### Viral vectors

Two adeno-associated virus (AAV) serotypes (AAV2/9, AAV9-retro) were utilized in this study. Viral particles were obtained from Taitool Bioscience Inc. (China) and Brain Case Inc. (China). The undiluted viral vector titers ranged from 0.8 to 1.5 × 10^13^ viral particles/mL, with a final injection titer of 5 × 10^12^ viral particles/mL.

### Surgery

#### Stereotaxic injection

Mice were deeply anaesthetized with 1% sodium pentobarbital (100 mg/kg body weight) and placed in a stereotactic frame (RWD Instruments, China). Virus was injected into the peri-LC (0.08 μL) (anterior-posterior (AP): −4.96 mm from bregma; medial-lateral (ML): ±0.74 mm; dorsal-ventral (DV): −3.71 mm from the dura) or somatomotor area (AP: 0.5 from bregma; ML: −1 mm; DV: −1 mm) using a Hamilton Ò microsyringe (Hamilton, USA) with a pressure microinjector (Picospritzer III, Parker, USA) at a rate of 0.01 μL/min. The injection needle was withdrawn 10 minutes after the end of the injection. For chemogenetic inactivation, injections were bilateral. For activation, injections were unilateral.

#### Optical fiber implantation

At 30 minutes after AAV injection, a ceramic ferrule with an optical fiber (200 mm OD, 0.37 NA, Newdoon, China) was implanted, positioning the optical fiber tip above the peri-LC (AP: −4.96 mm from bregma; ML: −0.74 mm; DV: −3.55 mm from the dura). The ferrule was secured on the skull with dental cement, and antibiotics were applied to the surgical wound to prevent infection. Fiber photometry recording and optogenetic stimulation were initiated at least 2 weeks post-implantation to ensure proper recovery and stable experimental conditions.

#### Distal femoral virus injection

Under anesthesia, the lower legs of each mouse were secured with medical tape. Using a microscope for precision, the skin was incised, and muscle tissue was carefully dissected along natural fiber lines with forceps. At the distal femur, a small hole was created using a 26G needle. A Hamilton Ò microsyringe (Hamilton, USA) attached to a pressure microinjector (Picospritzer III, Parker, USA) was used to inject 0.5 µL of PRV or AAV into the bone marrow cavity at a rate of 0.05 µL/min. Bone wax was applied to seal the junction between the syringe and the bone hole during the injection process. Ten minutes after completing the injection, the syringe was carefully withdrawn, and the bone hole was immediately sealed with bone wax. The skin was then sutured, and antibiotic gel was applied to the incision site.

Following surgery, mice were placed on a heat pad to recover from anesthesia. At the conclusion of the experiments, the mice were euthanized to confirm the accuracy of viral injection and optical fiber implantation. Brain sections were prepared at a thickness of 40 μm (Leica, CM1950, Germany) and counterstained with 4’,6-diamidino-2-phenylindole (DAPI). Fluorescent imaging was conducted using an Olympus VS120 (Japan) virtual microscopy slide scanning system. Only data from mice with confirmed injection accuracy were included in the analysis.

### Fiber photometry recording

Fiber photometry (ThinkerTech, China) was employed to record calcium activity in the peri-LC. GCaMP6s was delivered to neurons via intracranial injection of AAV-Syn-GCaMP6s. Calcium fluorescence signals were captured using an implanted optical fiber (200 µm OD, 0.37 NA, Newdoon, China) targeting the peri-LC. Calcium fluorescence recordings were conducted 2 weeks post-surgery. Mice were acclimatized to fresh enriched environment cages and standard cages for 1 hour per day over 3 days before data collection. Calcium fluorescence signals were collected using a photomultiplier tube and subsequently converted into electrical signals at a frequency of 40 Hz. GCaMP6s fluorescence was excited at 470 nm, and a 405 nm isosbestic channel was recorded as a calcium-independent reference. Before recording, dark offsets were measured under light-shielded conditions and subtracted from the raw signals. Offline preprocessing was performed in Python. The 405 nm reference trace was smoothed and fitted to the 470 nm signal using robust linear regression to estimate motion-related and non-calcium components. The corrected fluorescence signal was then converted to ΔF/F, interpolated only across brief artifact gaps, and low-pass filtered for downstream analysis. Behavioral events were manually annotated from synchronized overhead and lateral-view videos and aligned to the photometry time stamps. For event-aligned analyses, calcium traces were aligned to behavioral bout onset or offset and locally normalized using the 2-s pre-event baseline. For sustained behavioral bouts, duration-adjusted peri-event averaging was used to avoid contamination from subsequent behavioral states.

### RNA *in situ* hybridization

Specific RNA probes were commercially sourced from Spatial FISH Ltd..Mice were perfused with 0.1% diethylpyrocarbonate (DEPC)-treated PBS (Sigma, D5758, USA), followed by DEPC-treated PBS containing 4% PFA (PBS-PFA). After perfusion, brains were post-fixed overnight in DEPC-treated PBS-PFA, then transferred to DEPC-treated 30% sucrose solution at 4 °C for 30 hours. Brain sections (10 µm thick) were covered with reaction chambers for subsequent reactions. After dehydration and denaturation with methanol, hybridization buffer containing specific probes was applied, and the sections were incubated overnight at 37 °C. The sections were then washed with PBST three times and underwent target probe ligation in ligation mix for 3 hours at 25 °C. Following another round of washing, rolling circle amplification using Phi29 DNA polymerase was performed at 30 °C overnight. Fluorescent detection probes in hybridization buffer were then applied to the sections. Finally, the sections were dehydrated through a series of ethanol washes and mounted with a suitable medium. Imaging was performed using the Leica THUNDER Imaging System (Germany), and signal dots were decoded to obtain spatial RNA positioning information.

### Bone immunohistochemistry

Bone tissues freshly dissected from mice were immediately fixed overnight in ice-cold 4% PFA. Decalcification was performed using 10% EDTA (pH 7.2) at 4 °C with constant shaking. Once decalcified, the bones were submerged in a 20% sucrose and 2% polyvinylpyrrolidone (PVP) solution for 24 hours. The tissues were embedded and frozen in 8% porcine gelatin with 20% sucrose and 2% PVP. For immunofluorescent staining and analysis, 200-µm sections were obtained using low-profile blades on a Leica CM3050 cryostat (Germany). All bone tissues were processed, sectioned, stained, imaged, and analyzed under identical conditions. The sections were air-dried overnight, permeabilized with 0.3% Triton X-100, and blocked with 5% BSA at room temperature for 1 hour. Primary antibodies were diluted in 5% BSA in PBS and applied to the sections overnight at 4 °C. The following antibodies were used: Endomucin (sc-65495, Santa Cruz, USA, 1:50 dilution), Pecam1 (553370, BD Pharmingen, USA, 1:100), ADRB2 (ab182136, Abcam, UK, 1:100), TH (MAB318, Merck Millipore, USA, 1:500), and Tau-1 (MAB3420, Merck Millipore, 1:1 000). After incubation, the sections were washed three times with PBS, then incubated with Alexa Fluor-conjugated secondary antibodies (1:500, Abcam) for 2 hours at room temperature. Nuclei were counterstained with DAPI. The sections were thoroughly washed in PBS before mounting with Fluoromount-G (Southern Biotech, USA), with coverslips sealed using nail polish.

### Behavioral recording and analysis under different housing conditions

To analyze mouse behavior under different housing conditions, mice were individually placed in the corresponding environment and video-recorded from 9:00 PM to 12:00 AM. Active behavior was defined as any behavior involving body displacement, wheel running, or hanging upside down from the wire cage lid. The total duration of active behavior during the recording period was manually quantified for each mouse.

### Quantification of daily wheel-running activity

To quantify voluntary wheel-running activity, mice were individually housed in cages equipped with a running wheel for 24 h. Wheel-running activity was recorded using a wheel revolution counter system (ZNlogo, Tafit Co.Ltd., China) The total number of wheel revolutions was collected for each mouse over the 24-h recording period and used as the daily wheel-running revolution count.

### Forced treadmill running

Forced treadmill exercise was performed using a mouse treadmill system (SA101, SANS Biological Technology Co., Ltd., China). Mice were subjected to forced running for 4 weeks (6 days/week), from 4 to 8 weeks of age. The running distance, speed, and duration were gradually increased from the first day until the exercise duration reached 50 min per day on day 15; this regimen was then maintained through day 28. The final daily running distance (680 m/day, 5° incline) was set at approximately half of the daily voluntary wheel-running distance observed in EE mice, taking into consideration that forced treadmill running involves continuous exercise for 50 min, whereas voluntary wheel running under EE is intermittent.

### Measurement of plasma ionized calcium and Phosphate

Blood samples were collected from mice immediately after euthanasia into lithium heparin anticoagulant tubes. A portion of whole blood was analyzed within two hours for free calcium concentration using an i15 VET blood gas analyzer (EDAN Instruments, Shenzhen, China). The remaining blood was centrifuged to obtain plasma, and phosphate concentration was measured using a phosphate colorimetric assay kit (MAK308, Sigma-Aldrich, USA) according to the manufacturer’s instructions. Absorbance readings were acquired with a DLJ-100D microplate reader (Nanjing Dile Jia, China).

### Measurement of Blood Pressure and Heart Rate

Systolic and diastolic blood pressure, as well as heart rate, were measured in conscious mice using a BP2000 blood pressure analysis system (Visitech Systems, USA). All measurements were performed in a dedicated quiet room to minimize environmental stress. Prior to the formal recording session, mice underwent 3–5 days of acclimation to the apparatus and handling procedures. On the test day, data acquisition began only after five consecutive measurement cycles produced stable readings. Once this stabilization criterion was met, a total of 30 measurement cycles were recorded. Data were considered valid if at least 20 of the recorded cycles showed stable readings.

### Bone tissue clearing and analysis

#### Fixation

Mice were perfused with ice-cold PBS and 4% PFA. After sampling, the femurs were fixed in 4% PFA at 4 °C with gentle shaking overnight. After fixation, the femurs were washed with PBS 2–3 times (2 hours each).

#### Clearing and staining

Femurs were processed using a Nohai Tissue Clearing Kit (Cat No: NH240708-01, Nohai Biotechnology, China). Fixed femurs were placed in a centrifuge tube containing 15 mL of decalcification solution and gently shaken at 25 °C for 5 days, then placed in delipidation solution. After 20 days delipidation at 37 °C, the samples were transferred to PBS at room temperature and washed for 1 hour (repeated six times). The femurs were then stained with TH (AB152 or MAB318, Merck Millipore, 1:500) and EMCN (sc-65495, Santa Cruz, 1:50) antibodies (37 °C, 12 days), followed by relevant secondary antibodies (37 °C, 7 days). The samples were then immersed in refractive index matching solution until fully cleared, after which optical 3D imaging was performed.

#### 3D imaging and image processing of mouse femurs

Femurs were subjected to 3D fluorescence imaging using a Nohai LS18 light-sheet microscope (Nohai Biotechnology, China). For whole-femur 3D image acquisition, each field of view was illuminated six times by the light sheet, under a 1×/0.25 NA air objective (Olympus MVPLAPO)^58^. Based on the selected imaging conditions, the microscope magnification was set to 6.3×, with a spatial resolution of 2 × 2 × 5 µm^3^. When acquiring 3D fluorescence images of the epiphyseal region, each field of view was illuminated three times using a 10×/0.6 NA air objective lens (Olympus MVPLAPO). Under the selected imaging conditions, the microscope magnification was set to 5×, with a spatial resolution of 0.52 × 0.52 × 1.4 µm^3^. Data were modeled and analyzed using Amira (Thermo Fisher Scientific, USA, v 2021.2).

### Image acquisition and quantitative analysis

2D immunofluorescent staining was visualized at high resolution using a Zeiss laser scanning confocal microscope (LSM-880 and LSM-980, Germany). Z-stack images were acquired, processed, and 3D-reconstructed using Zeiss imaging software. Quantitative analyses were performed on high-resolution confocal images using ImageJ (NIH, USA).

### Micro-computed tomography (micro-CT) scanning and analysis

Femurs and tibiae were harvested from mice and immediately immersed in 4% PFA for fixation prior to micro-CT scanning. Scans were conducted using a Bruker Skyscan 1176/1276 micro-CT system (Kontich, Belgium) with the following parameters: 9 μm voxel size, medium resolution, 85 kV, 200 μA, 1 mm Al filter, and 384 ms integration time. Data analysis was conducted using the manufacturer’s proprietary evaluation software. Reconstruction of the scanned images was performed using NRecon (v1.7.4.2). Three-dimensional images were generated from contoured two-dimensional (2D) slices based on grayscale image transformation using CTvox software (v3.3.0). Both 3D and 2D analyses were performed using CT Analyser (v1.20.3.0).

Trabecular bone was identified as the region spanning 0.2–1.6 mm distal to the proximal growth plate of the femur, whereas cortical bone was analyzed in the region spanning 1.8–2.4 mm distal to the proximal growth plate. For trabecular bone analysis, the ROI was manually delineated slice-by-slice from 2D images to exclude cortical bone, followed by image binarization with a global gray level threshold (70–255) to isolate mineralized tissue. A Gaussian filter (radius = 1) was applied for 3D reconstruction. Quantitative analyses were performed on all bone regions within the ROI of the 3D images.

### Flow cytometry

Flow cytometry was conducted as described in previous research ^20^. Femurs and tibiae were dissected from mice, and the surrounding connective tissue was carefully removed. The metaphysis and diaphysis regions were crushed in Hank’s Balanced Salt Solution (Life Technologies, USA) supplemented with 10 mM HEPES (pH 7.2) (CellGro, USA). The bone fragments were enzymatically digested in a solution containing 2.5 mg/mL Collagenase A (Roche, Switzerland) and 1 U/mL Dispase II (Roche) for 15 minutes at 37 °C under gentle agitation.

The resulting cell suspensions were filtered through 40-μm mesh and washed in PBS (pH 7.2) with 0.5% BSA (Fraction V) and 2 mM EDTA. Following washing, equal numbers of cells from each mouse were blocked with Purified Rat Anti-Mouse CD16/CD32 (BD Biosciences) on ice for 30 minutes. Cells were then stained on ice for 45 minutes with the following antibodies: APC-conjugated EMCN (50-5851-80; eBioscience, USA), PE-conjugated CD31 (12-0311-81; eBioscience), FITC-conjugated CD45 (35-0451; Tonbo, USA), APC/Cy7-conjugated Ter119 (116223; BioLegend, USA), and PerCP/Cy5.5-conjugated CD146 (562231; BD Biosciences). After staining, cells were washed and resuspended in PBS (pH 7.2) containing 2 mM EDTA and 1 μg/mL DAPI for live/dead exclusion. For the bilateral Tie2-Cre and OC-Cre AAV experiments, the flow cytometry panel was modified to avoid spectral overlap with endogenous fluorescent reporters, including EGFP and tdTomato. Bone marrow cells were first incubated with anti-mouse CD16/CD32 to block Fc receptors and then stained with antibodies against CD31, Endomucin, CD45, and Ter119. The modified panel used APC-conjugated EMCN (50-5851-80; eBioscience, USA), BV421- conjugated CD31 (562939; BD Pharmingen, USA), PerCP-Cy5.5-conjugated CD45 (550994; BD Pharmingen, USA), and APC/Cy7-conjugated Ter119 (116223; BioLegend, USA), while endogenous EGFP or tdTomato fluorescence was detected directly without antibody staining. Flow cytometry analysis was performed using a CytoFLEX LX Flow Cytometer (Beckman Coulter, USA), and data were analyzed using CytExpert software.

### Two-photon imaging

Two-photon microscopy was used to visualize the vascular network within the distal femurs of anesthetized mice, providing detailed insights into endothelial cell morphology and behavior under varying experimental conditions. Mice were anesthetized with an intraperitoneal injection of 1% sodium pentobarbital (100 mg/kg body weight, Sigma) to minimize physiological stress and ensure stability during the imaging procedure. After anesthesia induction, mice were positioned on a custom-built imaging platform to provide stable access to the distal femur. Hair over the femoral region was removed using depilatory cream, and a small incision was made to expose the distal femur. The cortical bone was gently thinned using a bone drill. A window was carefully created in the bone without disturbing the bone marrow or vasculature, ensuring a clear view for imaging. Saline was periodically applied to prevent tissue desiccation.

Endothelial cell imaging was conducted using a two-photon resonant scanning microscope (Ultima Investigator, Bruker Nano, Germany) equipped with a 16× water-immersion objective (Nikon, 0.8 NA, Japan). Excitation was achieved using a femtosecond laser (Mai Tai, DeepSee, USA) operating at a 940 nm wavelength. Imaging was performed at 1× optical zoom with a resolution of 1 024 × 1 024 pixels using Prairie View software, enabling detailed visualization of endothelial cell morphology and their spatial arrangement within the vascular network. Quantitative analysis, focused on endothelial cell heterogeneity and vessel diameter in the distal femoral vasculature, was performed using ImageJ software (NIH, USA).

### Optogenetic light delivery

Optical fibers (200 µm diameter, NA 0.37, Newdoon, China) were connected to a blue LED light source (473 nm wavelength, Aurora-200, Newdoon, China). Optogenetic stimulation consisted of blue light pulses delivered at a frequency of 20 Hz, with a pulse width of 15 milliseconds, over a period of 10 seconds. Light intensity was set to 10 mW at the tip of the optical fiber.

### Oral gavage in mice

The propranolol (Sigma) doses were determined based on the clinical human dosage range (oral) ^59^ and human-to-mouse dose conversion formula ^60^. ICI-118551 was administered at a dose of 1.2 mg/kg/day for 4 weeks, from 4 to 8 weeks of age.

### Open field test

The open field apparatus consisted of a 50 cm × 50 cm square arena with 40 cm high walls, constructed from white plastic to provide a high-contrast background for the subjects. The apparatus was placed in a quiet room with dim lighting to minimize external distractions. The mice were habituated to the testing room for at least 1 hour prior to testing to reduce stress. Mice were tested within 1 hour following the administration of CNO (Sigma), rather than immediately after injection. Movements were tracked using an overhead camera, and locomotor activity, including total distance traveled, was automatically quantified using behavioral tracking software (ANY-maze, USA)

### Chronic restraint stress (CRS)

Mice were subjected to CRS by placement in 50-mL conical tubes with holes for air flow for 2–4 hours per day for 14 consecutive days.

### Sucrose preference test (SPT)

The SPT was employed to assess anhedonia, or the inability to experience pleasure ^61^. Mice were individually housed and acclimated to two bottles of water for 2 days, followed by 2 days of exposure to two bottles containing 2% sucrose solution. After a subsequent 24-hour period of water deprivation, mice were presented with one bottle of 2% sucrose solution and one bottle of water for 2 hours during the dark phase of their light cycle. The position of the bottles was switched 1 hour into the test to minimize side bias. Total fluid intake from each bottle was measured, and sucrose preference was defined as the average of the sucrose consumption ratio during the first and second hours. The sucrose consumption ratio was calculated by dividing the volume of sucrose consumed by total fluid intake (sucrose and water combined).

### Tail suspension test (TST)

The TST was conducted to evaluate behavioral despair in mice using the Bioseb Tail Suspension System (France). Each mouse was individually suspended by the tails with sterile tape affixed approximately 1 cm from the tail tip, which was then attached to the sensor. Mouse activity was recorded for 6 minutes, during which the system automatically quantified immobility time.

### Data analysis

All data are presented as mean ± standard error of the mean (SEM). For comparisons between two groups, paired or unpaired two-tailed Student’s *t*-tests were performed depending on the experimental design. One-way analysis of variance (ANOVA) was used for comparisons involving more than two groups, followed by appropriate *post-hoc* tests to determine group-specific differences. Two-way ANOVA with the Geisser-Greenhouse correction followed by Dunnett’s multiple comparisons test for comparisons with the pre-CNO baseline within each vascular region and Tukey’s multiple comparisons test for comparisons among vascular regions at each time point. Multiple comparisons were adjusted using the Benjamini, Krieger, and Yekutieli two-stage linear step-up procedure to control the false discovery rate (FDR) at 10%. For all tests, *p* < 0.05 was considered statistically significant. Statistical analyses were conducted using GraphPad Prism software (v8.0.2), and assumptions for normality and homogeneity of variances were checked prior to analysis.

### Declaration of generative AI and AI-assisted technologies in the writing process

During the preparation of this work, the author(s) used **ChatGPT** to check English grammar and enhance the fluency of the language. After using this tool, the author(s) reviewed and edited the content as needed and take(s) full responsibility for the content of the publication.

## Supplemental Figure Legends

**Figure S1:**
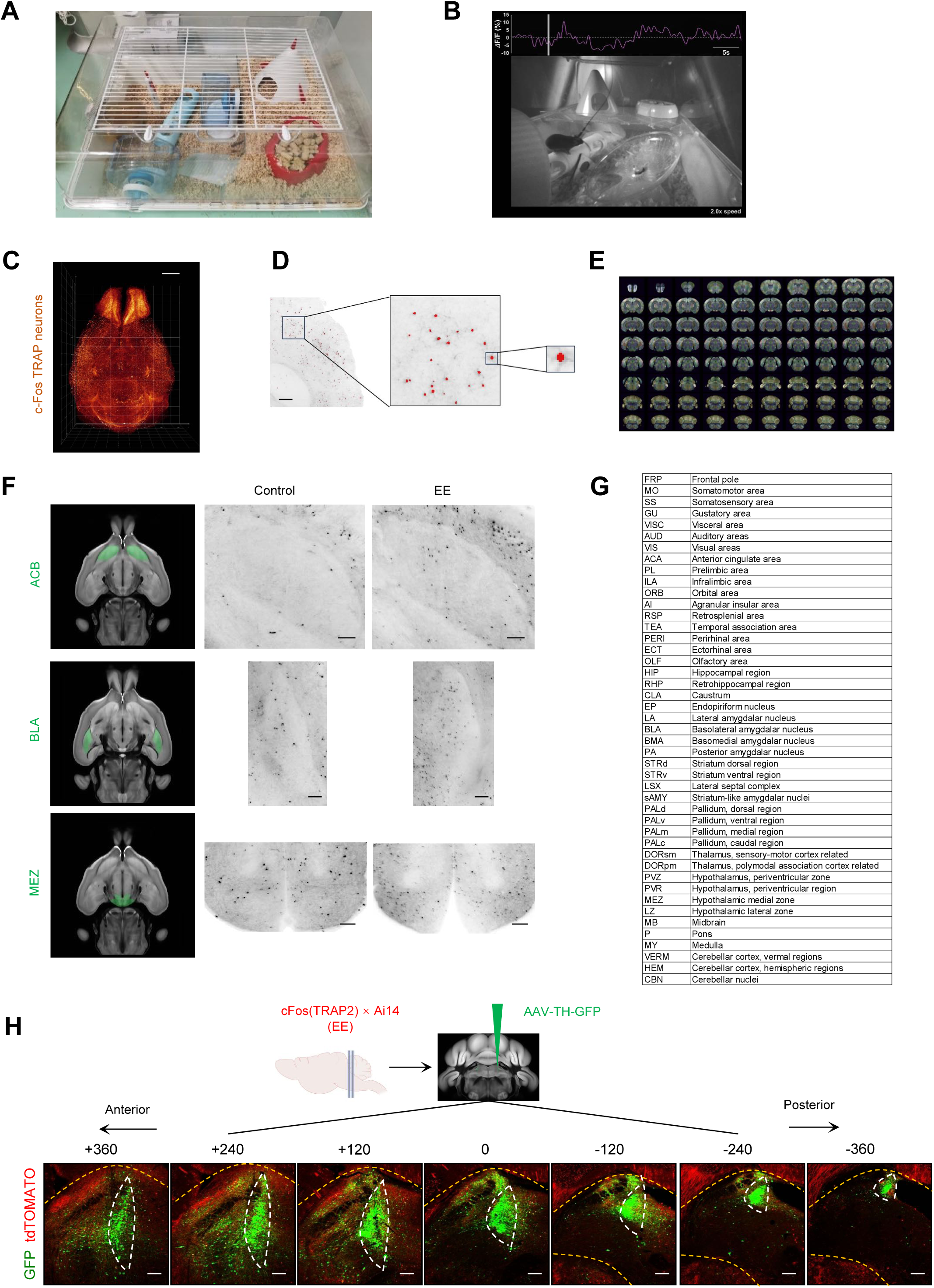
Activation of peri-LC neuronal populations by environmental enrichment. (A) Representative image of the enriched environment (EE) cage used in this study. The enriched cages measured 55 cm × 39 cm × 27 cm and consisted of two levels: the upper level contained a small tent and a food bowl, while the lower level included a water bottle, running wheel, climbing pole, tunnel, plastic ball, and soft tissue for nesting. Mice were housed at a density of 10–12 per cage, approximately double that of standard-housed mice. (B) Representative video still showing in vivo calcium photometry recording of a mouse in the EE cage. (C) Representative whole-brain clearing image of TRAP-labeled mouse brain. Scale bar, 2 mm. (D) Representative raw single-plane image of TRAP2-tdTomato-labeled brain acquired using a light-sheet microscope (left), alongside processed images with single cells outlined in red (*middle* and *right*). Scale bar, 400 μm. (E) Light-sheet image stacks registered to a common Allen Reference Atlas. (F) Representative light-sheet brain images of TRAP-labeled mice from standard and enriched environmental conditions, showing nucleus accumbens (ACB), basolateral amygdala nucleus (BLA), and hypothalamic medial zone (MEZ), outlined in green on axial reference slice. Scale bar, 200 μm. (G) Acronym table of Figure 1B. (H) cFos (TRAP2) × Ai14 mice (TRAP-labeled in enriched environments) injected with AAV-TH-GFP virus, followed by collection of coronal sections. Scale bar, 200 μm.

**Figure S2.**
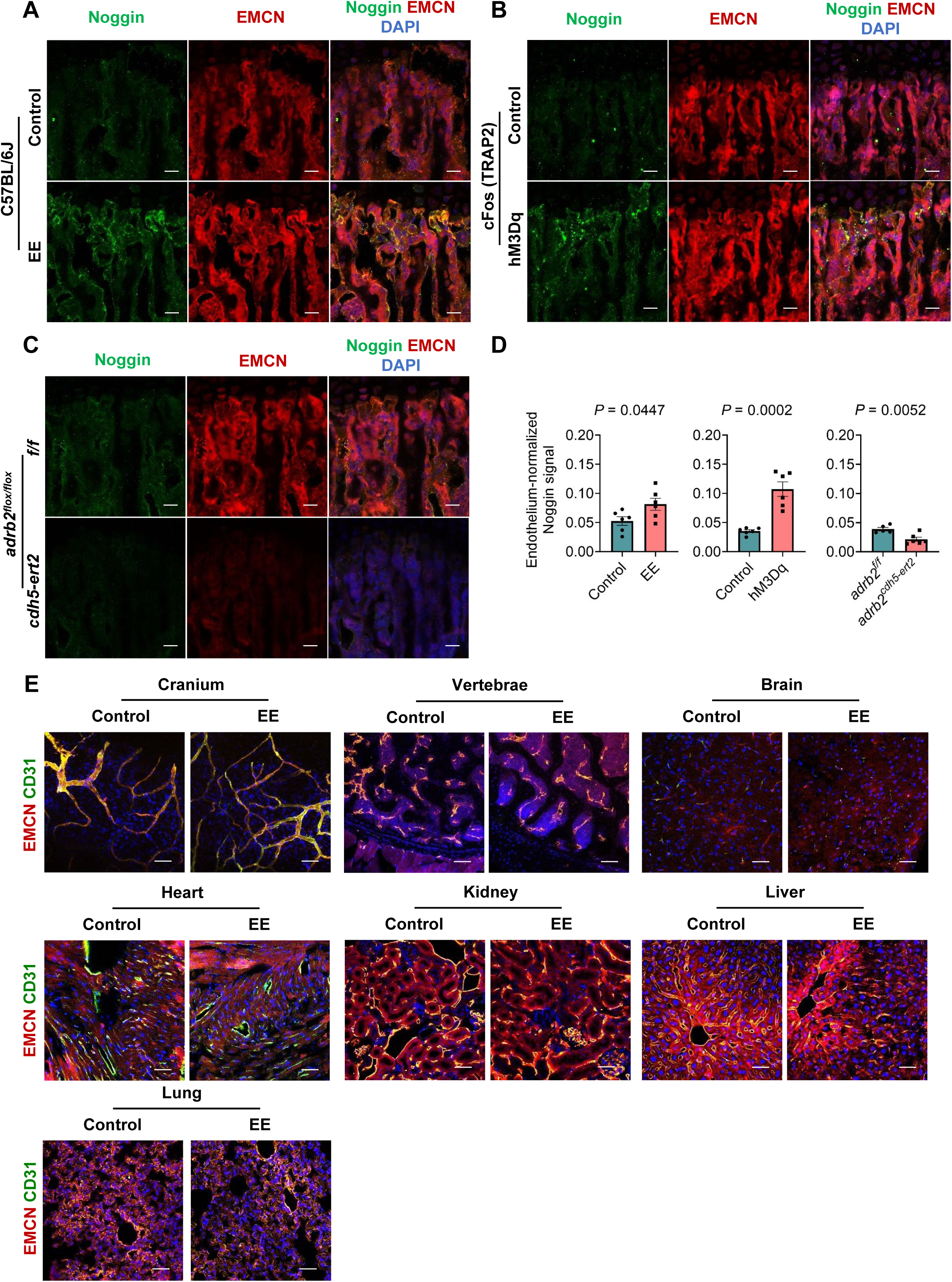
Expression of the endothelial-derived paracrine factor Noggin and tissue-specific endothelial responses under standard and enriched environment conditions. (A) Representative images showing the distribution of Noggin in femoral bone sections from C57BL/6J mice housed under standard housing (SH) or enriched environment (EE) conditions. Scale bars, 20 μm. (B) Representative images showing Noggin expression in cFos (TRAP2) mice following 3Dq-mediated activation of EE-labeled peri-LC neurons and in control mice. Scale bars, 20 μm. (C) Representative images showing Noggin expression in endothelial-specific Adrb2 knockout mice and control mice. Scale bars, 20 μm. (D) Quantification of Noggin expression under the experimental conditions shown in A–C. (E) Representative confocal images of cranium, vertebrate, brain, heart, kidney, liver, and lungs from 7-week-old mice housed in standard or enriched environments for 4 weeks, stained for EMCN (red), CD31 (green) and DAPI (blue). Scale bars, 50 μm.

**Figure S3.**
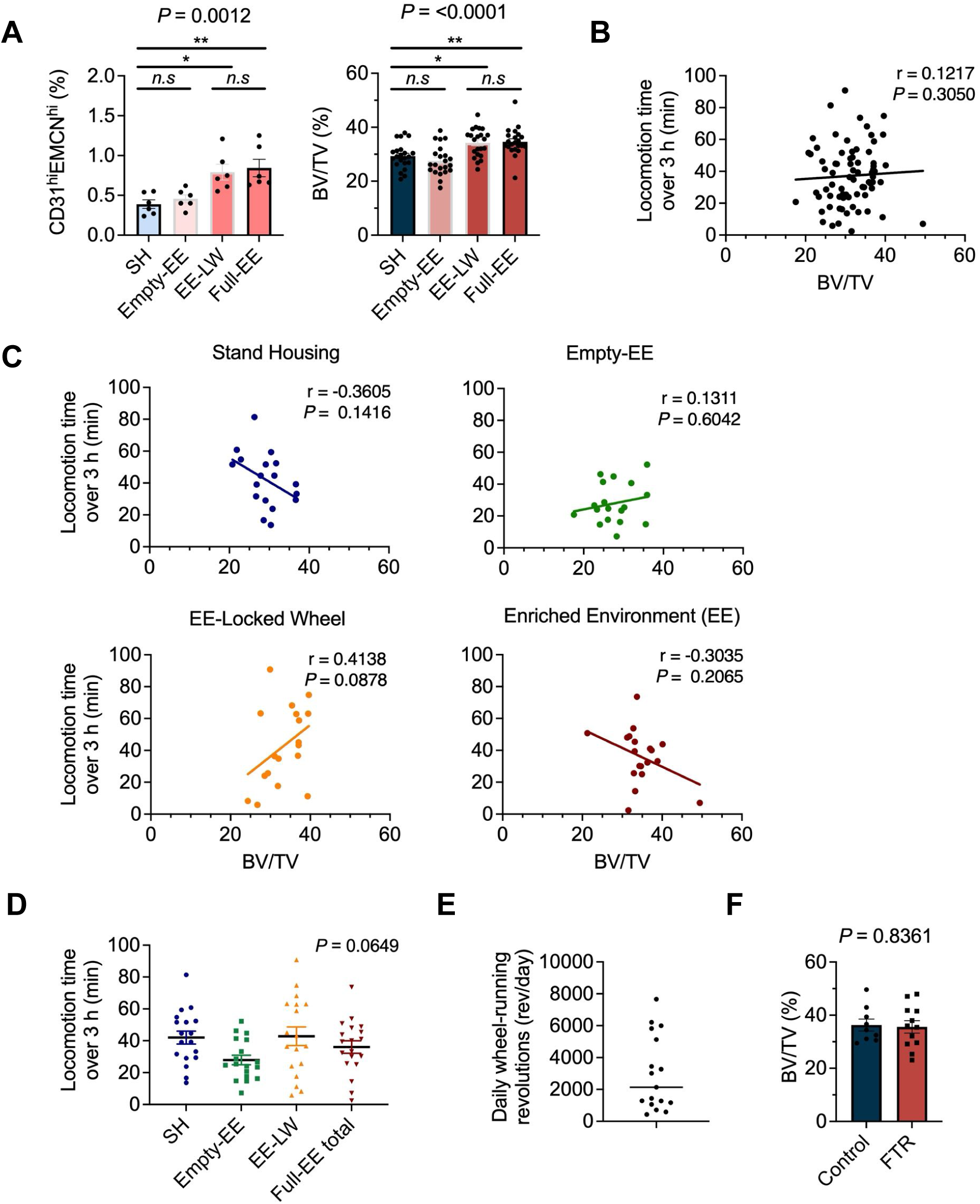
Analysis of the relationship between locomotor activity and bone mass changes under the experimental conditions used in this study. (A) Flow cytometric quantification of CD31^hi^EMCN^hi^ endothelial cells and micro-CT analysis of BV/TV in mice housed under SH, Empty-EE, EE-LW, or Full-EE conditions. Flow cytometric, n = 6, micro-CT, n = 22 (B) Correlation analysis between active time and BV/TV across all mice. n = 73. (C) Correlation analysis between active time and BV/TV within each housing condition. n = 18. (D) Quantification of active time under SH, Empty-EE, EE-LW, and Full-EE conditions. n = 18. (E) Quantification of daily wheel-running laps in EE-housed mice. n = 19. (F) Micro-CT analysis of BV/TV in forced treadmill running (FTR) and control mice. n = 9-12.

**Figure S4.**
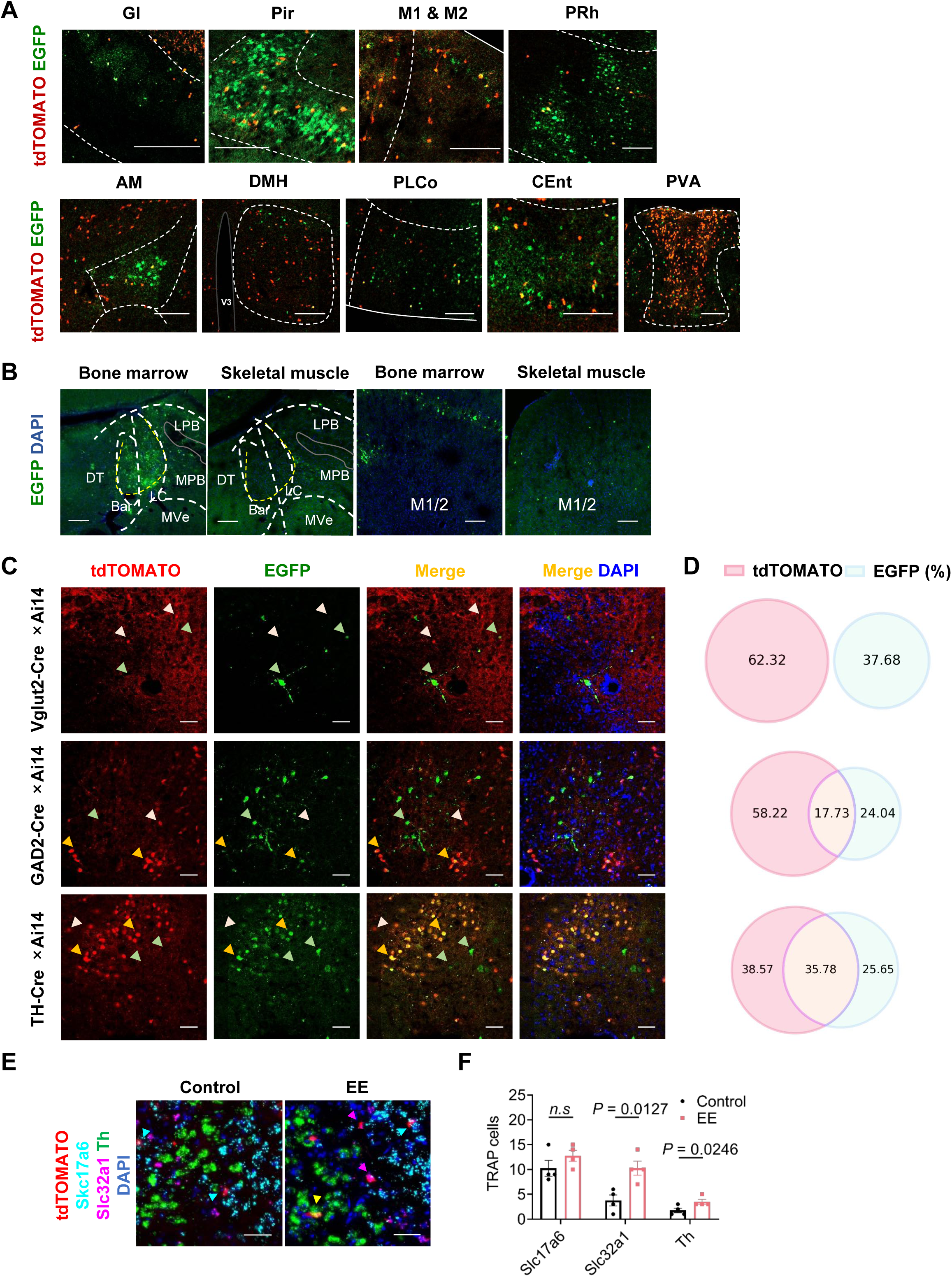
PRV-mediated transsynaptic tracing and analysis of cFos activation in TH-, GABA-, and VGLUT-expressing peri-LC neurons under enriched environment conditions. (A) Representative confocal images showing colocalization of PRV-EGFP-expressing neurons and TRAP tdTomato^+^ (EE-labeled) neurons in multiple brain regions. Gl, main olfactory bulb, glomerular layer; Pir, piriform cortex; M1&M2, primary motor area and secondary motor cortex; PRh, Perirhinal Cortex; AM, anterior medial nuclei; DMH, dorsomedial hypothalamic nucleus; PLCo, posterolateral cortical amygdala; CEnt, central entorhinal cortex, PVA, paraventricular nucleus of hypothalamus. Scale bars, 200 μm. (B) Representative image of PRV-mediated retrograde transsynaptic tracing from the bone marrow or adjacent skeletal muscle to the peri-LC and M1/M2 cortical regions. Scale bars, 100 μm. (C) Representative confocal images showing PRV-EGFP-expressing neurons in peri-LC in Vglut2-Cre × Ai14, GAD2-Cre × Ai14, and TH-Cre × Ai14 mice. Scale bars, 50 μm. (D) Quantification of colocalization between femur-PRV-tracing cells and tdTomato^+^ cells in the peri-LC of Vglut2-Cre × Ai14 (*upper*), GAD2-Cre × Ai14 (*right*), and TH-Cre × Ai14 mice (*lower*). n = 3. (E) Representative image of *in situ* hybridization for Slc17a6, Slc32a1, and Th in peri-LC in mice with TRAP-labeled neurons in standard or enriched environments. Colocalizations of tdTomato with Slc17a6 (cyan arrows), Slc32a1(magenta arrows), and Th (yellow arrow) are shown. Scale bar, 40 μm. (F) Quantification of neurons expressing Slc17a6, Slc32a1, and Th and colocalized with TRAP tdTomato in standard or enriched environments. n = 4.

**Figure S5.**
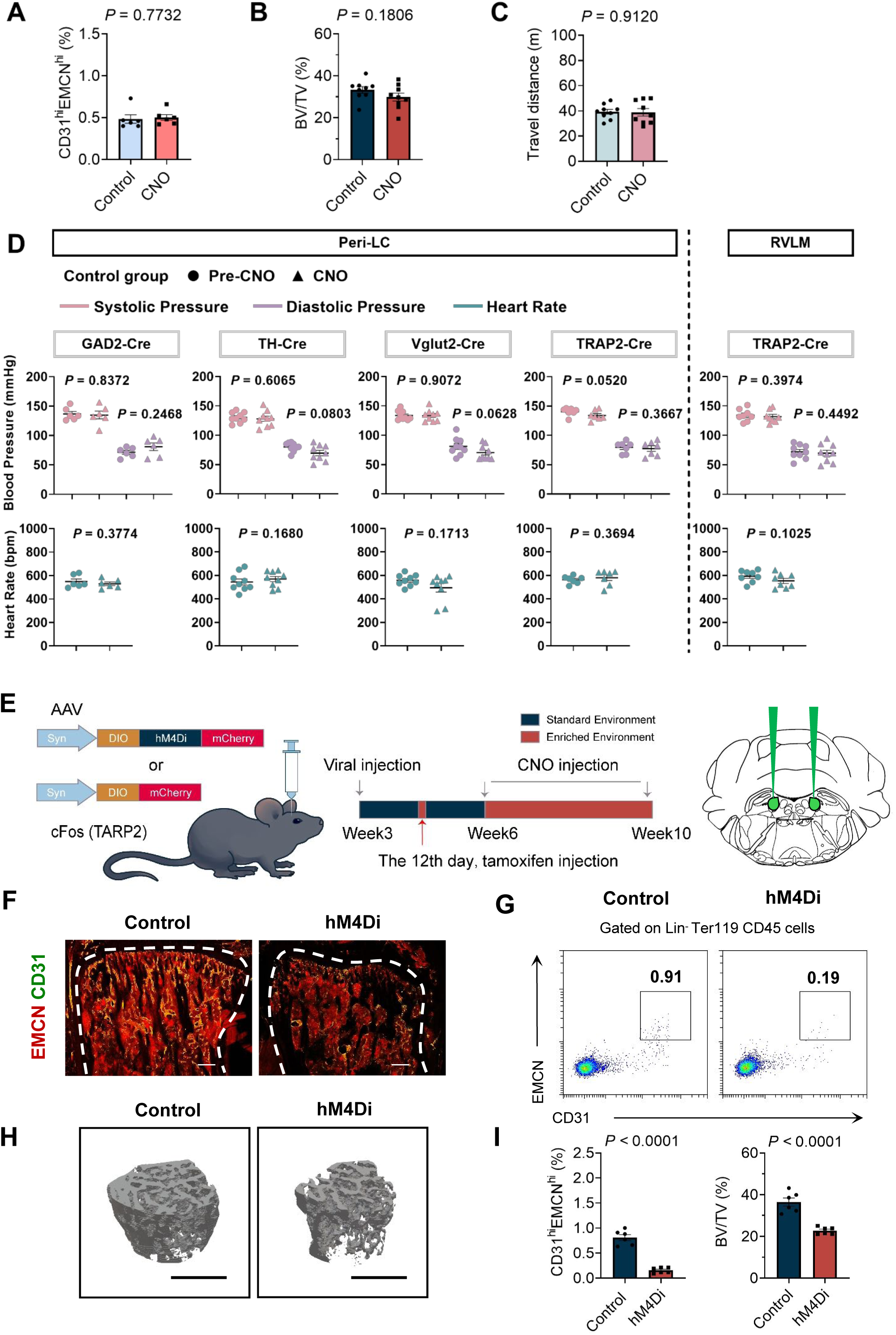
Effects of CNO administration alone and chemogenetic inhibition of EE-labeled peri-LC neurons. (A) Flow cytometry quantification of CD31^hi^EMCN^hi^ endothelial cells in C57BL/6J mice following saline or CNO administration every two days for two weeks, matching the dosing schedule and duration used in the neuronal activation experiments. n = 6. (B) Bone volume/total volume (BV/TV) in C57BL/6J mice following saline or CNO administration every two days for two weeks, matching the dosing schedule and duration used in the neuronal activation experiments. n = 9. (C) Total locomotor distance in the open field in mice injected with saline or CNO, measured within 0.5–2 h after injection. n = 9. (D) Blood pressure (*upper*) and heart rate (*lower*) before and after CNO administration in different transgenic mouse lines with control empty-virus injections into the indicated brain regions. n = 8-9. (E) Schematic representation of chemogenetic inhibition of EE-labeled peri-LC neurons in cFos (TRAP2) mice. Virus was injected at 3 weeks of age, followed by tamoxifen administration on day 12 after viral injection and CNO administration every two days during the experimental inhibition period. (F) Representative images of CD31 (green) and EMCN (red) dual-immunostained femur sections from EE-housed male mice with or without chemogenetic inhibition. Growth plate and endosteum are marked. Scale bars, 200 μm. (G) Representative flow cytometry plots showing CD31^hi^EMCN^hi^ endothelial cells from the femurs and tibiae of EE-housed male mice with or without chemogenetic inhibition. (H) Representative μCT images of trabecular bone in the distal femur. Scale bars, 1 mm. (I) *Left*, flow cytometric quantification of CD31^hi^EMCN^hi^ endothelial cells, n = 6; *right*, bone volume/total volume (BV/TV) of EE-housed male mice with or without chemogenetic inhibition, n = 6.

**Figure S6:**
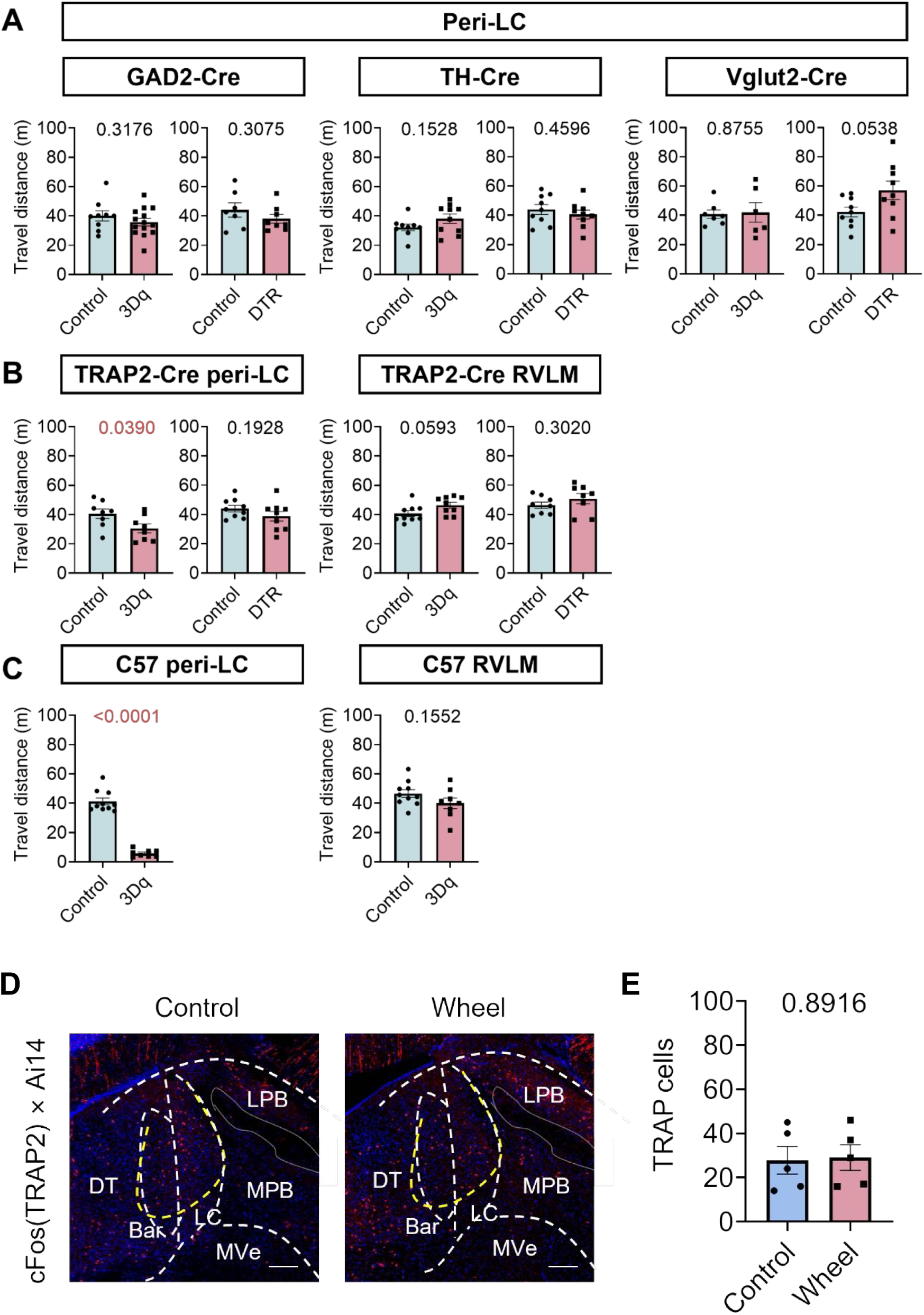
Effects of peri-LC, peri-LC neuronal subtype, and RVLM manipulations on locomotor activity and peri-LC response to running wheel. (A) Locomotor activity after activation or ablation of peri-LC neuronal subtypes. Gad2-Cre activation, n = 9-14; Gad2-Cre ablation, n = 7-8; TH-Cre activation, n = 9-10; TH-Cre ablation, n = 9; Vglut2-Cre activation, n = 6-7; Vglut2-Cre ablation, n = 9. (B) Locomotor activity after activation or ablation of TRAP2-labeled peri-LC or RVLM neurons. Peri-LC activation, n = 8; peri-LC ablation, n = 9; RVLM activation, n = 9; RVLM ablation, n = 8. (C) Locomotor activity after activation of peri-LC or RVLM neurons in C57BL/6J mice. Peri-LC activation, n = 8-10; RVLM activation, n = 8-10. (D) Representative confocal images showing activated TRAP (tdTomato^+^) cells in peri-LC (yellow dotted line) in mice exposed to standard with or without running wheel. Scale bar, 200 μm. (E) Quantification of TRAP (tdTomato^+^) cells in peri-LC (yellow dotted line) in mice exposed to standard with or without running wheel, n =5.

**Figure S7.**
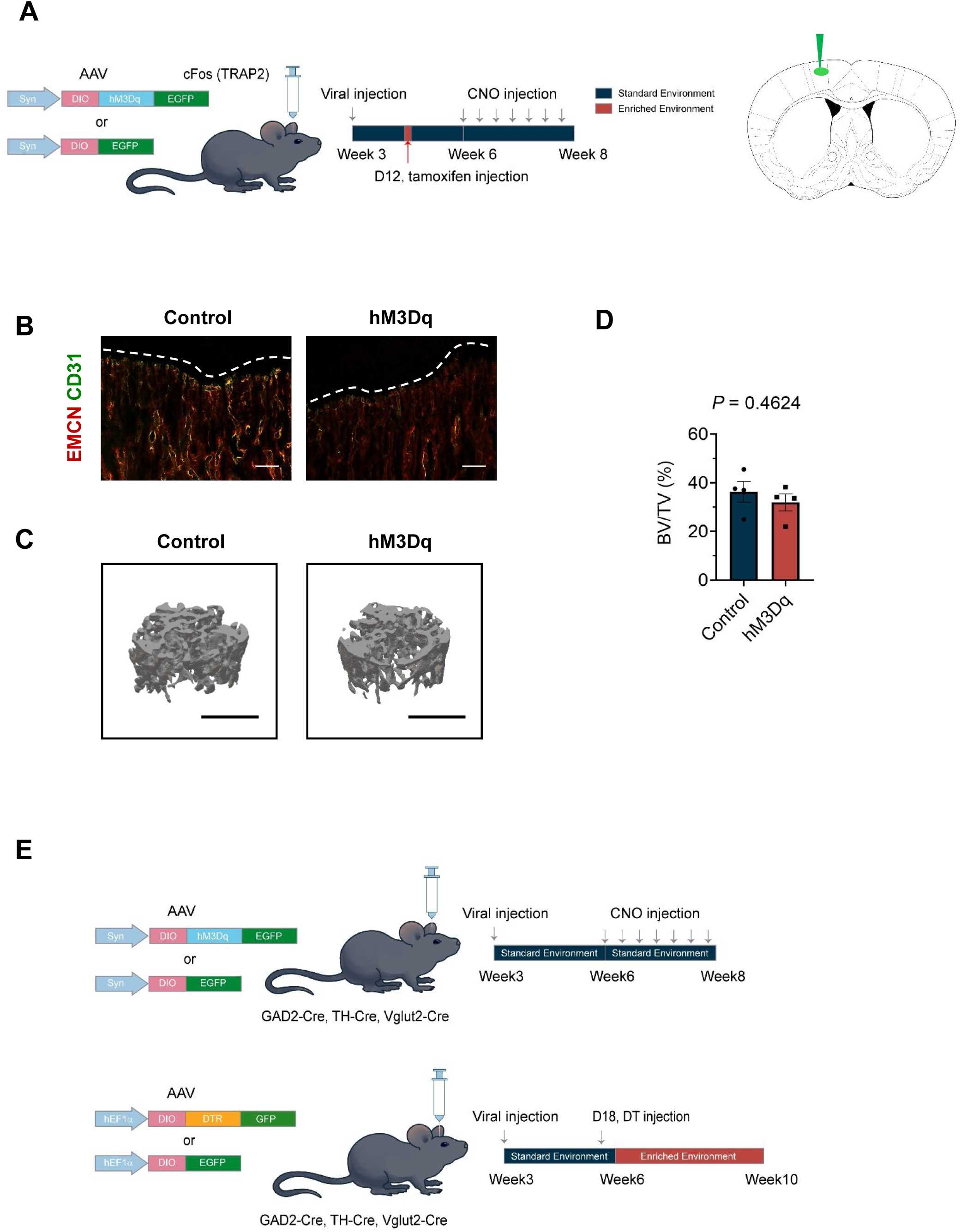
Effects of M1/M2 activation on skeletal endothelial heterogeneity and bone mass. (A) Schematic representation of chemogenetic activation of M1/M2 neurons in cFos (TRAP2) mice. CNO was administered every two days from week 6 to week 8. (B) Representative images of CD31 (green) and EMCN (red) dual-immunostained femur sections from standard-housed male mice with or without chemogenetic activation of M1/M2 neurons. The growth plate is marked. Scale bars, 100 μm. (C) Representative μCT images of trabecular bone in the distal femur. Scale bars, 1 mm. (D) Micro-CT analysis of bone volume/total volume (BV/TV) in standard-housed male mice with or without chemogenetic activation of M1/M2 neurons. n = 4. (E) Schematic representation of the experimental design for chemogenetic activation and DTR-mediated ablation of peri-LC neuronal subtypes. Corresponding results are shown in Figure 3K.

**Figure S8:**
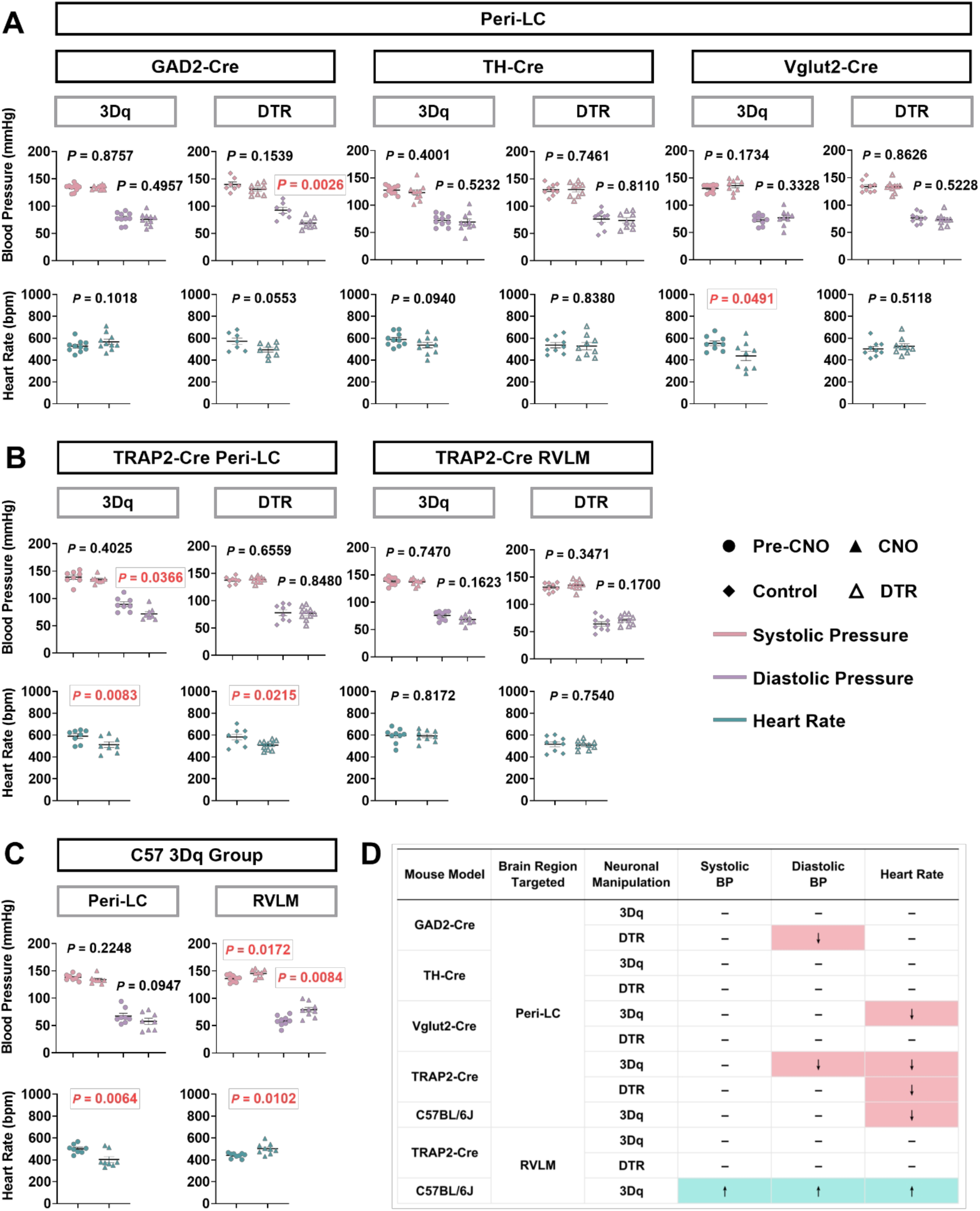
Effects of neuronal manipulation of the peri-LC and RVLM on blood pressure and heart rate across different mouse models. (A) Blood pressure and heart rate following activation or ablation of peri-LC neuronal subtypes. *Upper* panels, blood pressure; *lower* panels, heart rate. Gad2-Cre, activation, n = 10, ablation, n = 7-8; TH-Cre, activation, n = 10, ablation, n = 9; Vglut2-Cre, activation, n = 9, ablation, n = 9. (B) Blood pressure and heart rate following activation or ablation of TRAP2-labeled peri-LC or RVLM neurons. *Upper* panels, blood pressure; *lower* panels, heart rate. Peri-LC, activation, n = 8, ablation, n = 8-10; RVLM, activation, n = 9, ablation, n = 9. (C) Blood pressure and heart rate following activation of peri-LC or RVLM neurons in C57BL/6J mice. Peri-LC, activation, n = 8; RVLM, activation, n = 9. (D) Summary table showing statistically significant increases or decreases in blood pressure and heart rate across mouse models, brain regions, and neuronal manipulations.

**Figure S9:**
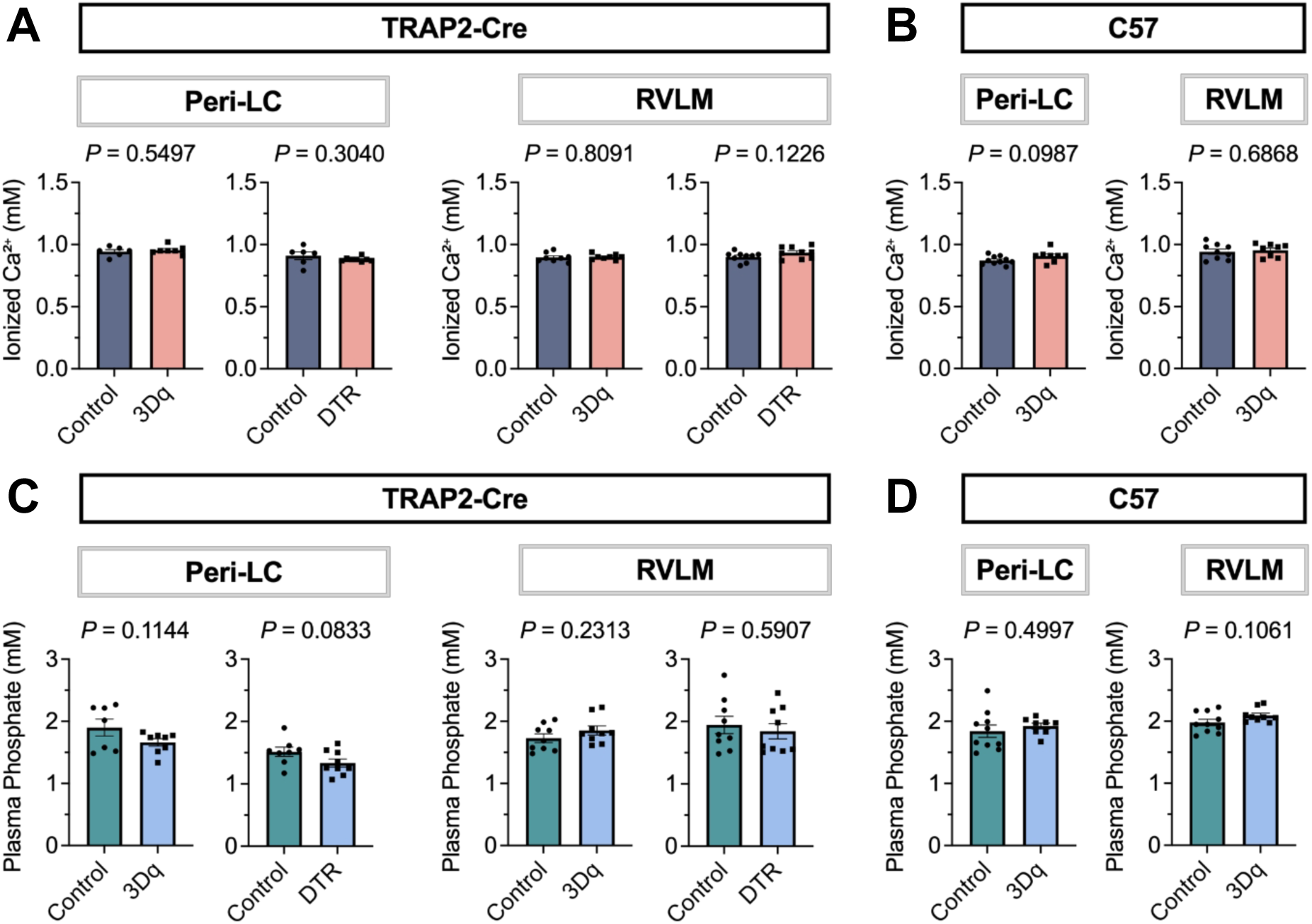
Effects of peri-LC and RVLM neuronal manipulation on circulating calcium and phosphate levels. (A) Circulating ionized calcium levels after activation or DTR-mediated ablation of TRAP2-labeled peri-LC or RVLM neurons. Peri-LC activation, n = 6-7; peri-LC ablation, n = 6-8; RVLM activation, n = 8; RVLM ablation, n = 9. (B) Circulating ionized calcium levels after activation of peri-LC or RVLM neurons in C57BL/6J mice. Peri-LC activation, n = 8-10; RVLM activation, n = 8-9. (C) Circulating phosphate levels after activation or DTR-mediated ablation of TRAP2-labeled peri-LC or RVLM neurons. Peri-LC activation, n = 7-8; peri-LC ablation, n = 8-9; RVLM activation, n = 9; RVLM ablation, n = 9. (D) Circulating phosphate levels after activation of peri-LC or RVLM neurons in C57BL/6J mice. Peri-LC activation, n = 8-10; RVLM activation, n = 9-10.

**Figure S10:**
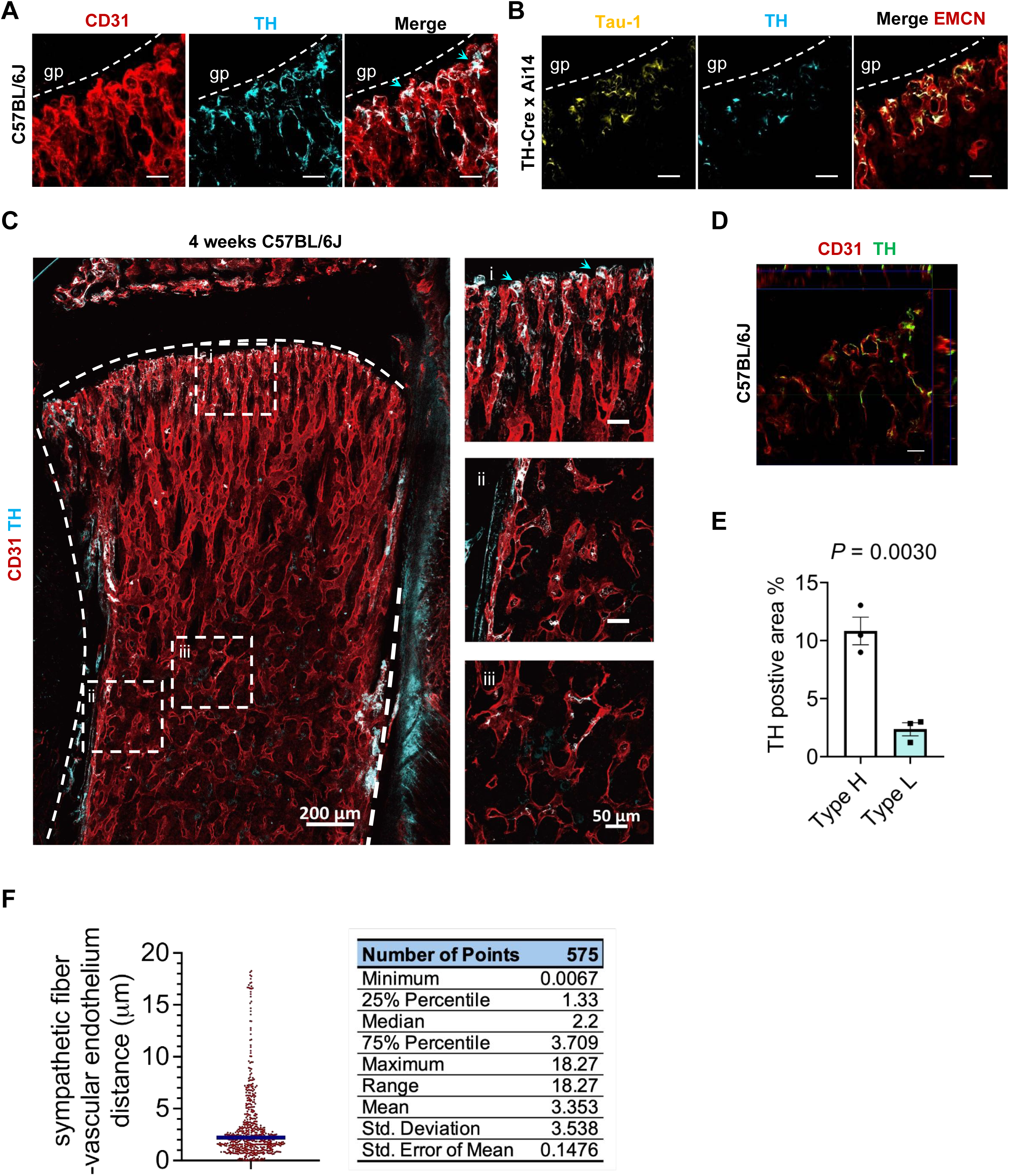
Heterogeneous distribution of sympathetic nerve terminals in long bones. (A) Representative images of EMCN and TH dual-immunostained femur sections from wild-type mice. The growth plate is marked. Cyan arrows indicate sympathetic fiber foot-like structures. Scale bars, 30 μm. (B) Representative images of Tau-1 and EMCN dual-immunostained femur sections from TH-Cre × Ai14 mice. The growth plate is marked. Scale bars, 30 μm. (C) Representative confocal images showing TH^+^ sympathetic fibers and vascular endothelial cells in femur sections from 4-week-old wild-type mice. Cyan arrows indicate sympathetic fiber foot-like structures. Scale bars, *left*, 200 μm; *right*, 50 μm. (D) Single optical section and orthogonal views corresponding to the confocal z-stack shown in A, illustrating the spatial relationship between sympathetic fiber structures and the vascular endothelial surface. (E) Quantification of relative distribution of sympathetic nerve terminals associated with two types of endothelial cells in long bones. n = 3. (F) Quantitative 3D proximity analysis of the distance between TH^+^ sympathetic fibers and the vascular endothelial surface. *Left*, the blue line indicates the median distance. *Right*, the accompanying table summarizes the distribution of measured distances using descriptive statistics.

**Figure S11:**
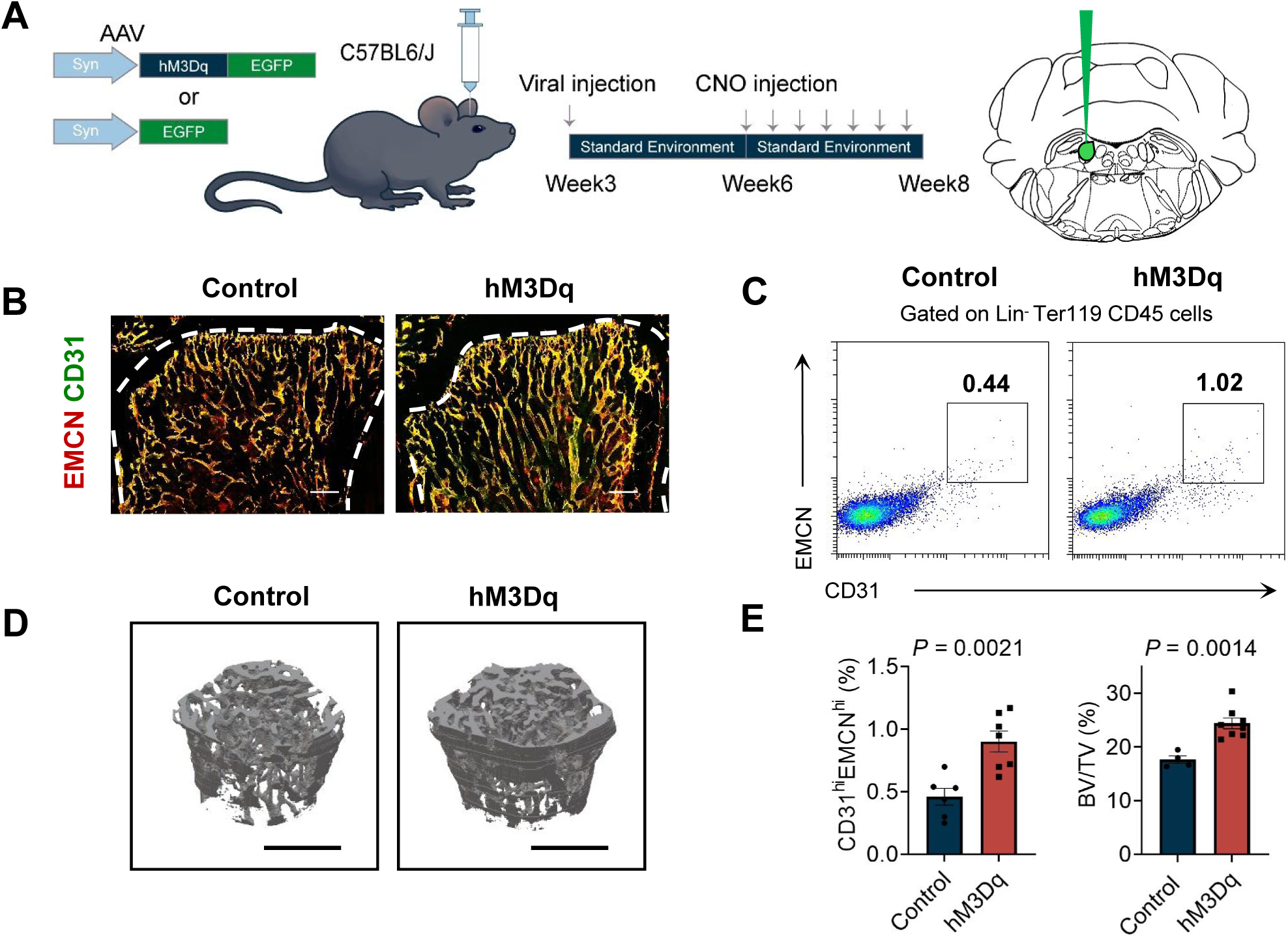
Chemogenetic regulation of endothelial cell heterogeneity in long bones via the peri-LC in wild-type mice. (A) Schematic representation of chemogenetic inactivation of the peri-LC in wild-type mice. CNO was administered every two days from week six to week ten. (B) Representative images of CD31 (green) and EMCN (red) dual-immunostained femur sections from EE-housed male mice with or without chemogenetic inactivation. Growth plate and endosteum are marked. Scale bars, 200 μm. (C) Representative flow cytometry plots showing CD31^hi^EMCN^hi^ endothelial cells from femurs and tibiae of EE-housed male mice with or without chemogenetic inactivation. (D) Representative μCT images of trabecular bone in the distal femur, Scale bars, 1 mm. (E) *Left*, Flow cytometry quantification of CD31^hi^EMCN^hi^ endothelial cells, n = 6–7, and *right*, bone volume/total volume (BV/TV) of EE-housed male mice with or without chemogenetic inactivation. Additional micro-CT parameters, including Ct.Th, Tb.Th, Tb.N, and Tb.Sp, are provided in Table 8. n = 5.

**Figure S12:**
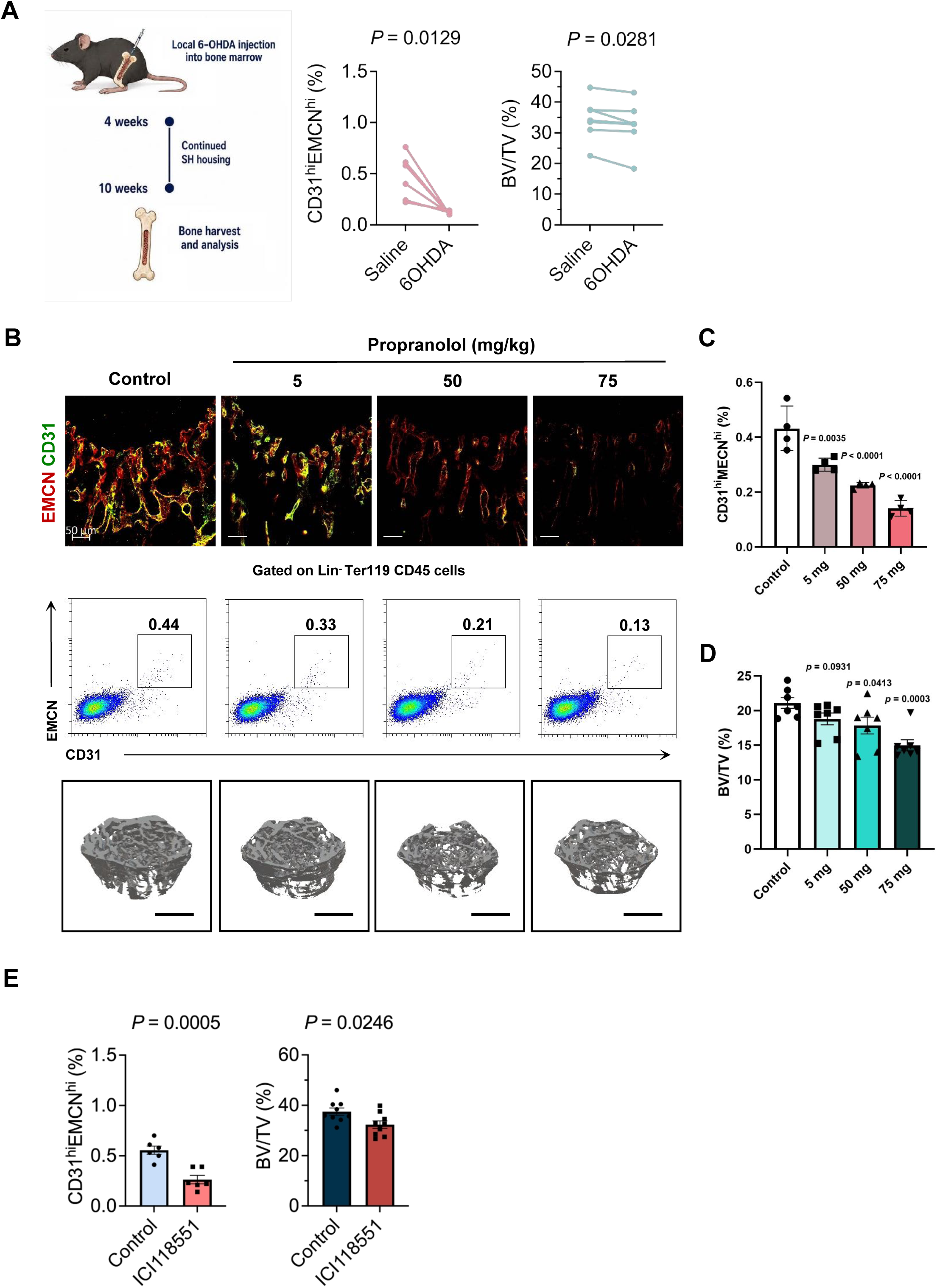
Reduction of CD31^hi^EMCN^hi^ endothelial cell levels and bone density in juvenile wild-type mice following treatment with beta-adrenergic receptor blocker propranolol. (A) *Left*, schematic representation of unilateral 6-OHDA injection into the femoral bone marrow of wild-type mice, with the contralateral femur serving as an internal control for analysis of skeletal endothelial cells and bone mass. *Middle*, flow cytometric quantification of CD31^hi^EMCN^hi^ endothelial cells, n = 6. *Right*, comparison of bone volume/total volume (BV/TV), n = 7. (B) *Upper*, representative images of CD31 (green) and EMCN (red) dual-immunostained femur sections from juvenile wild-type mice with or without propranolol gavage treatment. Propranolol was administered once daily for 4 weeks starting at 4 weeks of age. Scale bars, 50 μm. *Middle*, Representative flow cytometry plots showing CD31^hi^EMCN^hi^ endothelial cells from femurs and tibiae. *Lower*, Representative μCT images of trabecular bone in the distal femur, Scale bars, 1 mm. (C) Flow cytometry quantification of CD31^hi^EMCN^hi^ endothelial cells from propranolol-treated and control mice. n = 4. (D) Bone volume/total volume (BV/TV) of propranolol-treated and control mice. n = 7. (E) *Left*, flow cytometry quantification of CD31^hi^EMCN^hi^ endothelial cells from ICI118551-treated and control mice, n = 6. *Right*, bone volume/total volume (BV/TV) of ICI118551-treated and control mice. Additional micro-CT parameters, including Ct.Th, Tb.Th, Tb.N, and Tb.Sp, are provided in Table 17. n = 9.

**Figure S13:**
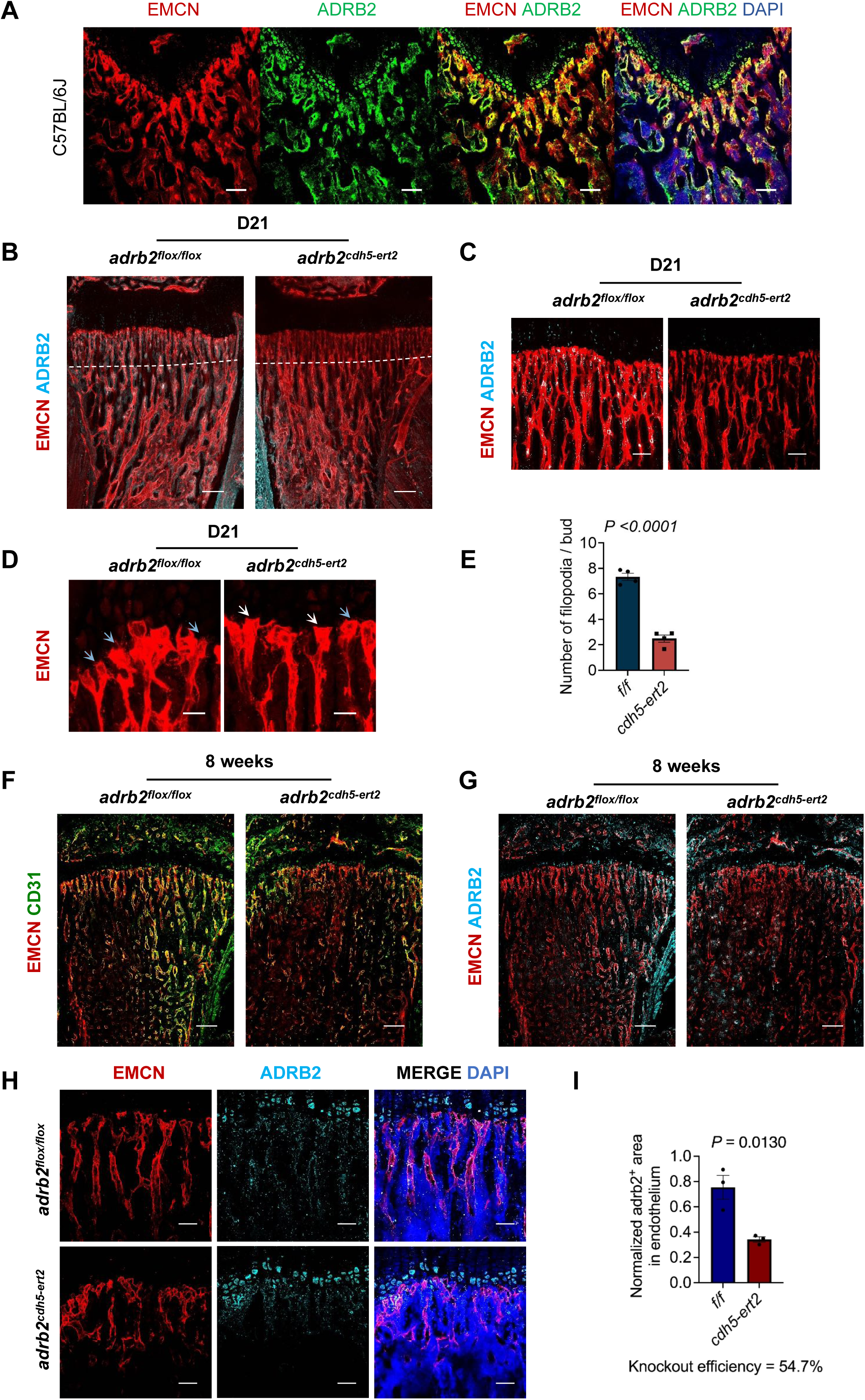
β2-adrenergic receptor expression and endothelial sprouting morphology in endothelial-specific Adrb2 knockout mice. (A) Representative images showing β2-adrenergic receptor expression around vascular endothelial cells near the growth plate in femur sections from 4-week-old mice. (B) and (C) Representative images of ADRB2 (cyan) and EMCN (red) dual-immunostained femur sections from D21 *adrb2*^flox/flox^ and *adrb2*^cdh5-ert2^ mice. Scale bars, B, 200 μm, C, 100 μm. (D) Representative images of D21 *adrb2*^flox/flox^ and *adrb2*^cdh5-ert2^ mouse femurs stained with EMCN. Blue arrows indicate normal endothelial sprouting, white arrows point to morphological changes, showing endothelial sprouts lacking filopodia. Scale bars, 30 μm. (E) Quantification of filopodia in endothelial sprouting. n = 4. (F) Representative images of CD31 (green) and EMCN (red) dual-immunostained femur sections from 8-week-old *adrb2*^flox/flox^ and *adrb2*^cdh5-ert2^ mice. Scale bars, 200 μm. (G) Representative images of ADRB2 (cyan) and EMCN (red) dual-immunostained femur sections from 8-week-old *adrb2*^flox/flox^ and *adrb2*^cdh5-ert2^ mice. Scale bars, 200 μm. (H) Representative images of ADRB2 (cyan), EMCN (red), and DAPI (blue) triple-immunostained femur sections from 8-week-old *adrb2*^flox/flox^ and *adrb2*^cdh5-ert2^ mice. Scale bars, 50 μm. (I) Quantification of endothelial β2-adrenergic receptor expression near the growth plate in femur sections from 8-week-old *adrb2*^flox/flox^ and *adrb2*^cdh5-ert2^ mice. n = 3.

**Figure S14:**
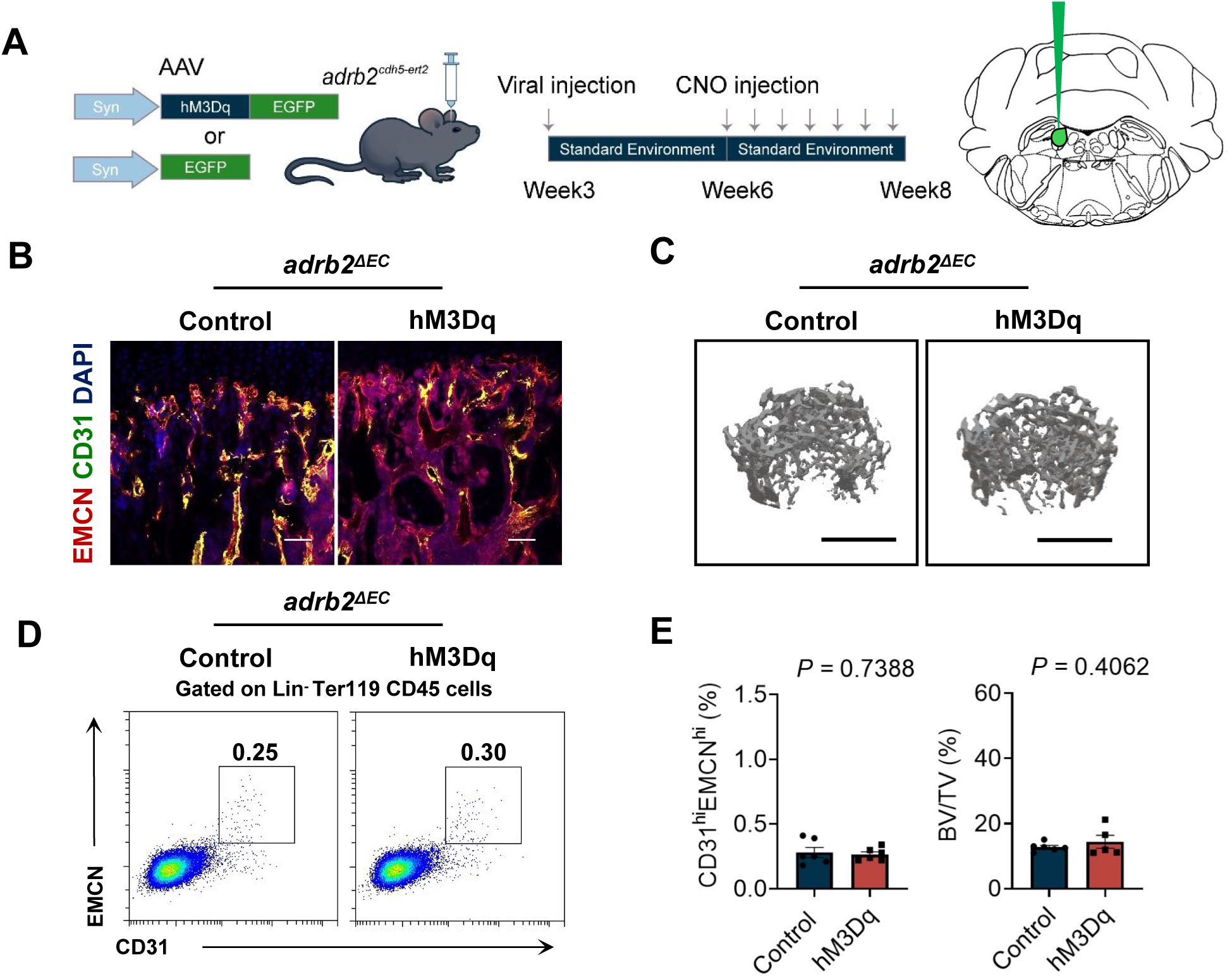
Endothelial-specific knockout of beta2-adrenergic receptors abolishes the effect of peri-LC activation. (A) Schematic representation of chemogenetic activation of the peri-LC in *adrb2*^cdh5-ert2^ mice. CNO was administered every two days from week six to week eight. (B) Representative images of immunostained femur sections from *adrb2*^cdh5-ert2^ mice with or without chemogenetic activation of the peri-LC, scale bar, 50 μm. (C) Representative μCT images of trabecular bone in distal femur, scale bars, 1 mm. (D) Representative flow cytometry plots showing CD31^hi^EMCN^hi^ endothelial cells from the femurs and tibiae of *adrb2*^cdh5-ert2^ mice with or without chemogenetic activation of the peri-LC. (E) *Left*, Flow cytometry quantification of CD31^hi^EMCN^hi^ endothelial cells, and *right*, bone volume/total volume (BV/TV) of *adrb2*^cdh5-ert2^ mice with or without chemogenetic activation of the peri-LC. n = 5-6.

**Figure S15:**
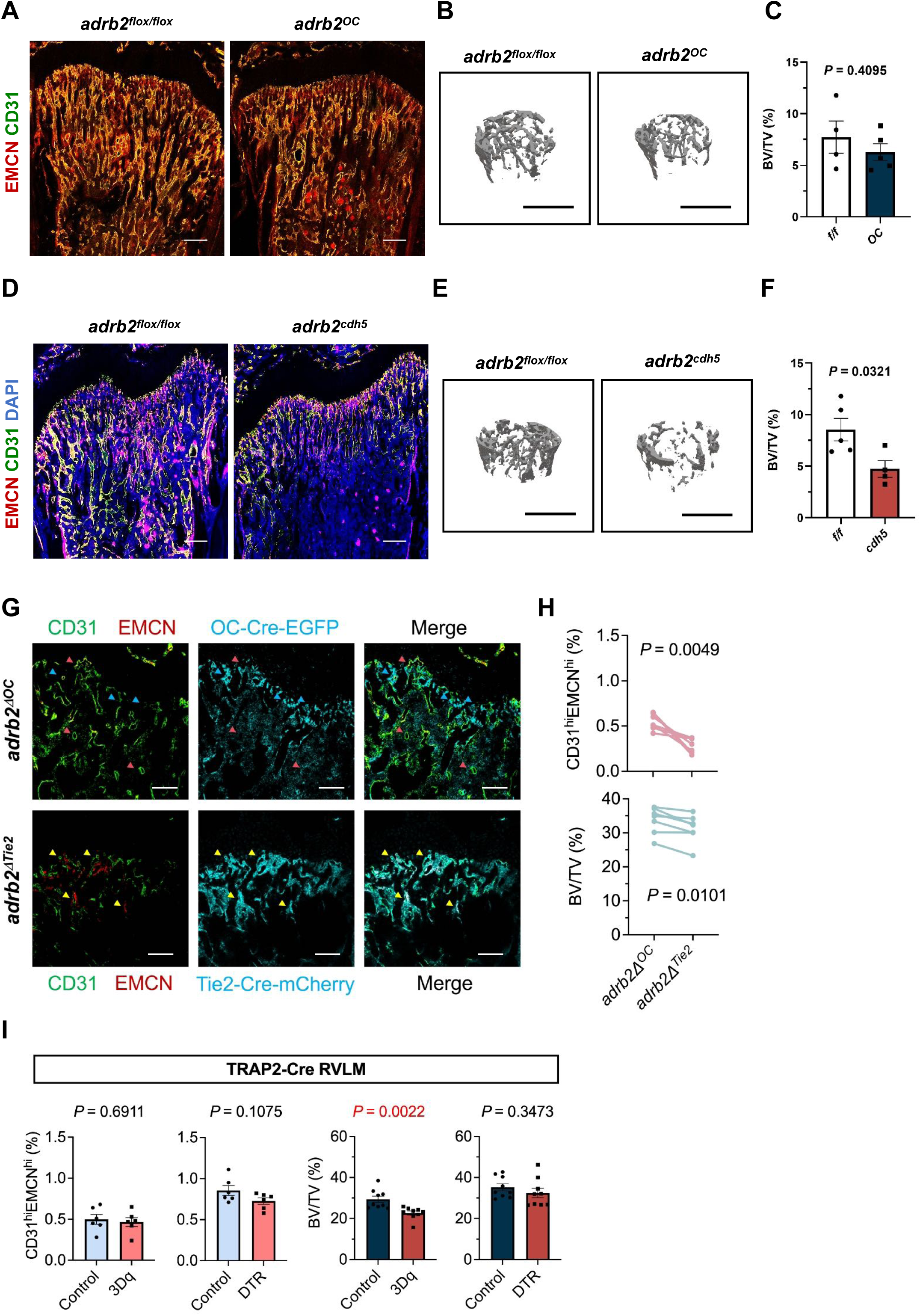
Comparison of endothelial cell heterogeneity and bone mass following specific knockout of *adrb2* in osteoblasts and endothelial cells. (A) Representative images of CD31 (green) and EMCN (red) dual-immunostained femur sections from 4-week-old *adrb2*^flox/flox^ and *adrb2*^oc^ mice. Scale bars, 200 μm. (B) Representative μCT images of trabecular bone in the distal femur, scale bars, 1 mm. (C) Bone volume/total volume (BV/TV) in 4-week-old *adrb2*^flox/flox^ and *adrb2*^oc^ mice. n = 4–5. (D) Representative images of CD31 (green) and EMCN (red) dual-immunostained femur sections from 4-week-old *adrb2*^flox/flox^ and *adrb2*^cdh5^ mice. Scale bars, 200 μm. (E) Representative μCT images of trabecular bone in the distal femur, scale bars, 1 mm. (F) Bone volume/total volume (BV/TV) of 8-week-old *adrb2*^flox/flox^ and *adrb2*^cdh5^ mice. n = 4–5. (G) Representative images of femur sections following local cell type-specific Adrb2 deletion in the same animal, with endothelial-specific Adrb2 deletion in one femur and osteoblast-specific Adrb2 deletion in the contralateral femur. Scale bar, 50 μm. (H) *Upper*, flow cytometric quantification of CD31^hi^EMCN^hi^ endothelial cells, n = 7; *lower*, micro-CT analysis of BV/TV in femurs with endothelial- or osteoblast-specific Adrb2 deletion. Additional micro-CT parameters, including Ct.Th, Tb.Th, Tb.N, and Tb.Sp, are provided in Table19. n = 7. (I) Quantification of CD31^hi^EMCN^hi^ endothelial cells and BV/TV following activation or DTR-mediated ablation of TRAP2-labeled RVLM neurons, Additional micro-CT parameters, including Ct.Th, Tb.Th, Tb.N, and Tb.Sp, are provided in Table 6-7. Activation, n = 6, DTR, n = 9.

**Figure S16:**
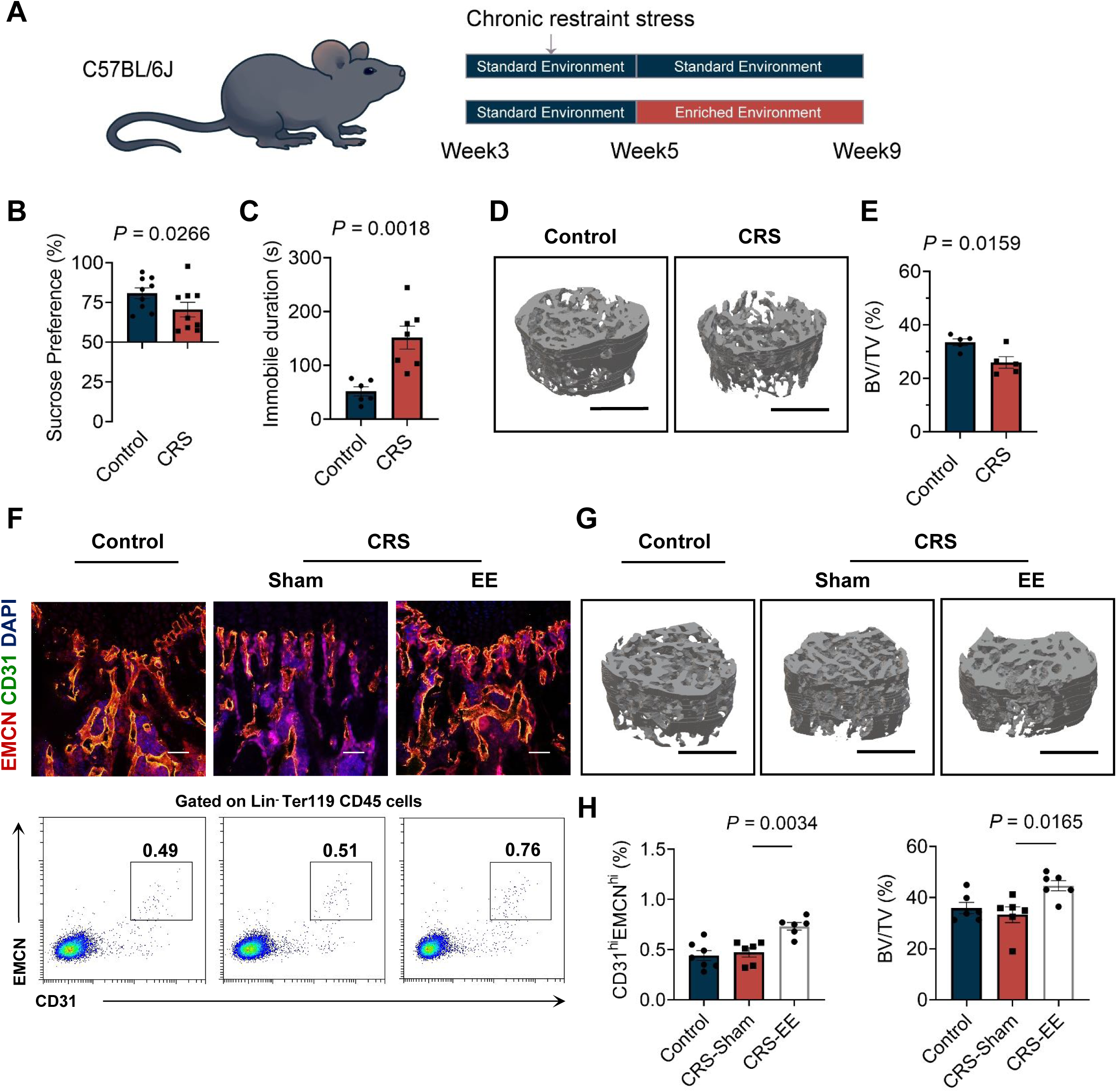
Increased bone density in CRS mice following environmental enrichment. (A) Schematic representations of standard- or EE-housed CRS mice. (B) Behavioral effects of CRS in mice in SPT. n = 9. (C) Behavioral effects of CRS in mice in TST. n = 7. (D) Representative μCT images of trabecular bone in the distal femur in control and CRS mice, scale bars, 1 mm. (E) Bone volume/total volume (BV/TV) of control and CRS mice. n = 5. (F) *Upper*, Representative images of immunostained-femur sections from standard- or EE-housed CRS mice, scale bar, 50 μm. *Lower*, Representative flow cytometry plots showing CD31^hi^EMCN^hi^ endothelial cells from the femurs and tibiae of standard- or EE-housed CRS mice. (G) Representative μCT images of trabecular bone in the distal femur, scale bars, 1 mm. (H) *Left*, Flow cytometry quantification of CD31^hi^EMCN^hi^ endothelial cells, and *right*, bone volume/total volume (BV/TV) of standard- or EE-housed CRS mice. n = 6.

**Figure S17:**
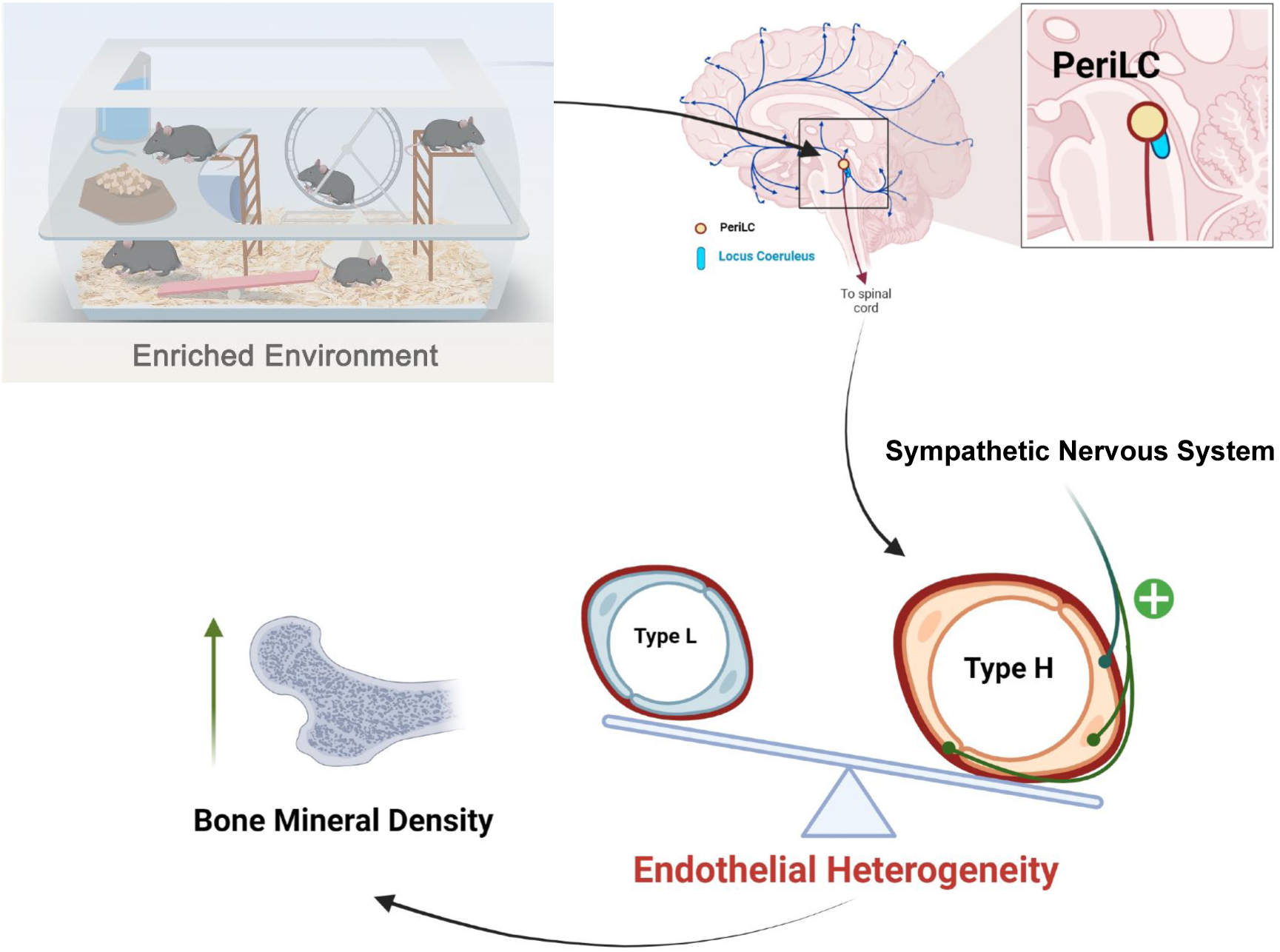
Summary graphs illustrating how enriched environments maintain vascular endothelial heterogeneity and enhance bone density via the peri-LC and sympathetic nervous system.

